# Bioen-OSMOSE: A bioenergetic marine ecosystem model with physiological response to temperature and oxygen

**DOI:** 10.1101/2023.01.13.523601

**Authors:** Alaia Morell, Yunne-Jai Shin, Nicolas Barrier, Morgane Travers-Trolet, Ghassen Halouani, Bruno Ernande

**Affiliations:** IFREMER, Unité halieutique Manche Mer du Nord Ifremer, HMMN, Boulogne sur mer, France; MARBEC, Univ. Montpellier, Ifremer, CNRS, IRD, Sète/Montpellier, France; DECOD (Ecosystem Dynamics and Sustainability), IFREMER, INRAE, Institut Agro, 44311 Nantes, France; International Institute for Applied Systems Analysis (IIASA), A-2361 Laxenburg, Austria

**Keywords:** Bioenergetic, Food web, Hypoxia, Marine ecosystem model, Phenotypic plasticity, Thermal tolerance

## Abstract

1. Marine ecosystem models have been used to project the impacts of climate-induced changes in temperature and oxygen on biodiversity mainly through changes in species spatial distributions and primary production. However, fish populations may also respond to climatic pressures via physiological changes, leading to modifications in their life history that could either mitigate or worsen the consequences of climate change.
2. Building on the individual-based multispecies ecosystem model OSMOSE, Bioen-OSMOSE has been developed to account for high trophic levels’ physiological responses to temperature and oxygen in future climate projections. This paper presents an overview of the Bioen-OSMOSE model, mainly detailing the new developments. These consist in the implementation of a bioenergetic sub-model that mechanistically describes somatic growth, sexual maturation and reproduction as they emerge from the energy fluxes sustained by food intake under the hypotheses of a biphasic growth model and plastic maturation age and size represented by a maturation reaction norm. These fluxes depend on temperature and oxygen concentration, thus allowing plastic physiological responses to climate change.
3. To illustrate the capabilities of Bioen-OSMOSE to represent realistic ecosystem dynamics, the model is applied to the North Sea ecosystem. The model outputs are confronted with population biomass, catch, maturity ogive, mean size-at-age and diet data of each species of the fish community. A first exploration of current species spatial variability in response to temperature or oxygen is presented in this paper. The model succeeds in reproducing observations, with good performances for all indicators.
4. This new model development opens the scope for new fields of research such as the exploration of seasonal or spatial variation in life history in response to biotic and abiotic factors at the individual, population and community levels. Understanding such variability is crucial to improve our knowledge on potential climate change impacts on marine ecosystems and to make more reliable projections under climate change scenarios.

## 1. Introduction

The development of increasingly realistic marine ecosystem models (MEMs) is needed to improve understanding and knowledge about marine ecosystems, which is one of the main challenges of the UN Decade of the Oceans (Heymans et al., 2020). MEMs are end-to-end models representing ecosystems from primary production to top predators, linking the species and/or functional groups via trophic interactions. These models also account for abiotic and human activity impacts on ecosystem dynamics (Rose et al., 2010; Steenbeek et al., 2021; Travers et al., 2007). MEMs are still being improved through the development of sub-models that increase their reliability in supporting ecosystem-based management (Pikitch et al., 2004; Rose et al., 2010).

The rates of ocean temperature rise and deoxygenation make urgent the development of mechanistic tools to forecast realistically their impacts from the physiology of marine organisms, to the population demographic impacts and to the consequence on marine trophic webs (Breitburg et al., 2018; Urban et al., 2016). Efforts to model the temperature impacts on marine biodiversity at the ecosystem level has so far focused mainly on the bottom-up effect of temperature on the ecosystem via changes in primary production (Lefort et al., 2015; Moullec et al., 2019) and on the distribution shift of species according to their preferred temperature (Albouy et al., 2014; Fernandes et al., 2013; Moullec et al., 2019; Serpetti et al., 2017). Mechanistic physiological response to temperature in MEM is modeled in size spectrum models and to our knowledge is not currently incorporated into an explicit multispecies model (Lefort et al., 2015; Maury, 2010). Although oxygen concentration is considered as a main pressure on marine biodiversity (Laffoley & Baxter, 2019), the oxygen physiological impact on marine ecosystems is still not explicitly modeled in MEMs.

The core of recent model developments linking environmental conditions and physiological response is primarily on single-species models. These frameworks mechanistically describe life history cycles and metabolic fluxes. The response of metabolic rates to temperature is used in several frameworks (Gillooly et al., 2002; Kooijman, 2010) which are applied to project future population dynamics and spatial distribution under climate change scenarios. The response of metabolic rates to oxygen through its impact on ingestion (Thomas et al., 2019) has been recently introduced in the Dynamic Energy Budget framework to study the impact of hypoxia on population dynamics (Lavaud et al., 2019).

The model Bioen-OSMOSE is a new framework that mechanistically describes the emergence of life history traits through an explicit description of the underlying bioenergetic fluxes and their response to food, temperature and oxygen variation in a multispecies food web model. It has been developed from the OSMOSE framework (Shin & Cury, 2004; www.osmose-model.org), which is an individual-based, spatially and temporally explicit, multispecies model for regional marine ecosystems. Designed to be possibly coupled to ocean and biogeochemical models, it includes the components of the entire ecosystem, from primary production to fish populations and human fishing activity, but the core of the model describes the dynamics of fish and macroinvertebrate species. In this paper, we provide a detailed description of the principles and equations of the Bioen-OSMOSE framework, as well as parameterization guidelines (detailed in Supporting Information). An application to the North Sea ecosystem is provided as a case study example. We then confront simulation outputs from the North Sea example to observed data to assess the consistency of the new model development and explore spatial variability in fish metabolic fluxes in response to temperature and oxygen.

## 2. Method

### 2.1. Model description

The Bioen-OSMOSE model (Fig. 1) represents fish individual physiological responses to temperature and oxygen variations and their consequences on fish communities in marine ecosystems. It is an individual-based, spatially and temporally explicit multispecies model accounting for trophic interactions. The main characteristics of the model are opportunistic predation based on size adequacy and spatiotemporal co-occurrence of predators and prey, the mechanistic description of individuals’ life-history traits emerging from bioenergetics. The aims of the model are to explore the functioning of marine trophic webs, the ecosystem impacts of individual physiological modifications due to temperature and oxygen, and the consequences of fishing pressure or climate change, from individual phenotype, to the population and to the community scale. The Bioen-OSMOSE model extends the existing OSMOSE model by (i) explicitly accounting for the mechanistic dependence of life-history traits on bioenergetics and (ii) describing intra- and inter-specific phenotypic variability originating from plastic responses to spatio-temporal biotic and abiotic factor variations. A process overview is given in Supporting Information S1.

**Figure 1:**
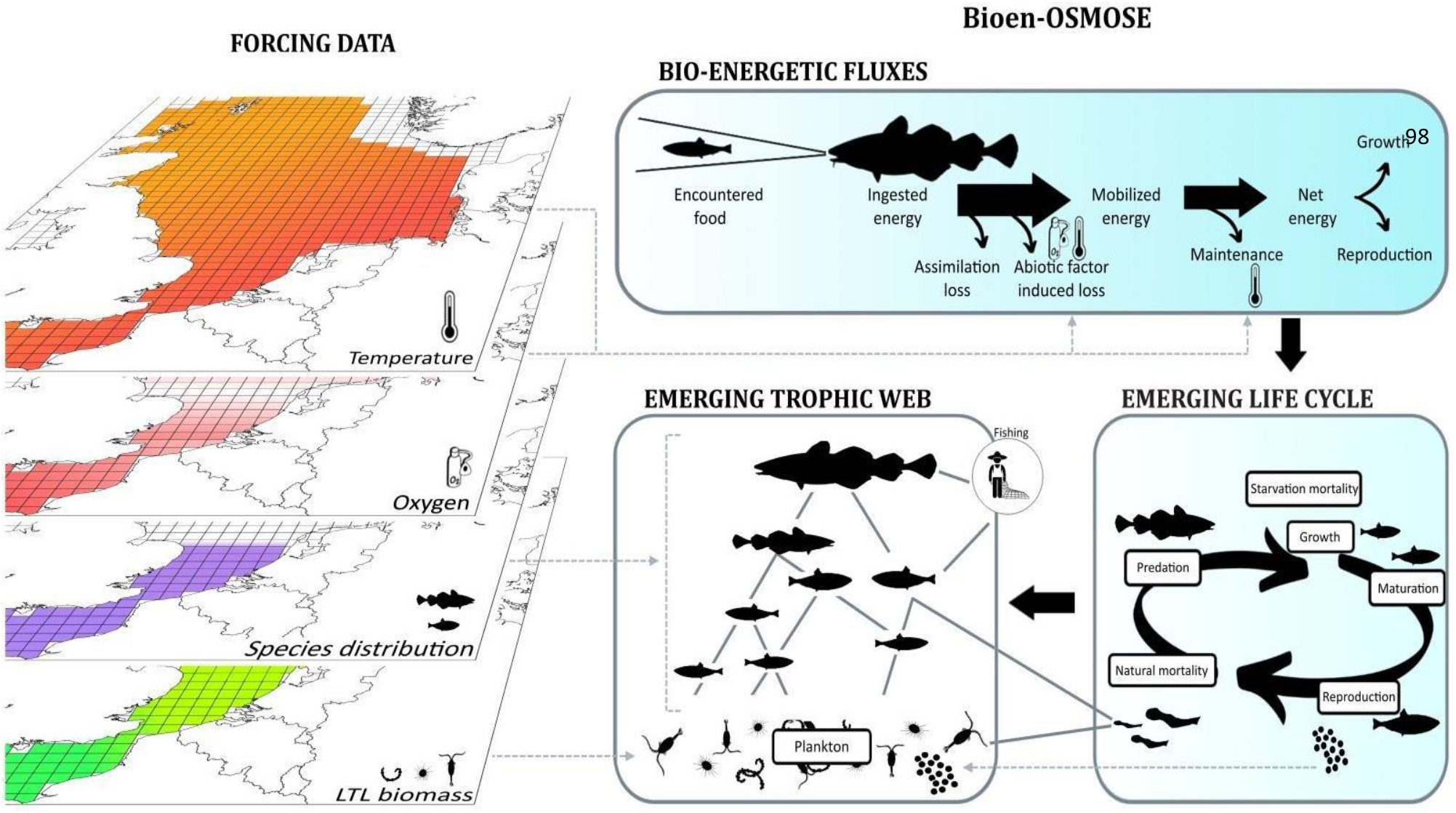
Graphical description of the Bioen-OSMOSE model. In the Bioen-OSMOSE model trophic relationships emerge from spatio-temporal co-occurrence and size adequacy between predators and prey, the former resulting from ontogenic spatial distributions and possibly LTL biomass distribution. The life cycle emerges from the underlying bioenergetic fluxes that describe the internal processes from energy ingestion (which relies on the encountered prey) to growth, maturation and reproduction. The internal fluxes are partly driven by environmental conditions, i.e, temperature and oxygen.

#### 2.1.1. Biological unit, state variables and spatial characteristics

The biological unit of the model is a school (a super-individual in individual-based modeling terminology). It is formed by individuals from the same species that are biologically identical. The state variables characterizing a school *i* at time step *t* belong to four categories (see Table 1 for state variable definitions and their units):

- Ontogenic state of individuals described by their age *a*(*i,t*), somatic mass *w*(*i,t*) and gonadic mass *g*(*i, t*);
- Abundance, namely the number of individuals in the school *N*(*i, t*);
- Spatial location, i.e., the grid cell *c*(*i, t*) where the school is located; and
- Taxonomic identity, i.e., the species *s*(*i*) to which the school belongs.

**Table 1:** Variables and functions of the bioenergetics and life-history sub-models. (Δ*t* t:time step duration)

| Symbol | Description | Units | Equations |
| --- | --- | --- | --- |
| **Entities: Fish schools** |  |  |  |
| **Ontogenic state** |  |  |  |
| *State variables* |  |  |  |
| $a(i, t)$ | Age of school $i$'s individuals at time step $t$ | y | 3, 13 |
| $w(i, t)$ | Somatic mass of school $i$'s individuals at time step $t$ | g | 2, 7, 10,11 |
| $g(i, t)$ | Gonadic mass of school $i$'s individuals at time step $t$ | g | 12, 14 |
| *Emerging individual variables* |  |  |  |
| $L(i, t)$ | Total length of school $i$'s individuals at time step $t$ | cm | 1,13 |
| $m(i, t)$ | Maturity state of school $i$'s individuals at time step $t$ | – | 10, 13 |
| $a_m(i)$ | Maturation age of school $i$'s individuals | y | |
| $w_m(i)$ | Maturation somatic mass of school $i$'s individuals | g | |
| $L_m(i)$ | Maturation length of school $i$'s individuals | cm | |
| $N_{eggs}(i, t)$ | Total fecundity of school $i$ at first time step $t$ of the breeding season | # | 14, 15 |
| *Abundance: State variable* |  |  |  |
| $N(i, t)$ | Number of individuals in school $i$ at time step $t$ | # | 1, 14,15 |
| $B(i, t)$ | Biomass of school $i$ at time step $t$ | g | 1 |
| *Spatial Location: State variable* |  |  |  |
| $c(i, t)$ | Grid cell of school $i$ at time step $t$ | – | 1 |
| *Taxonomic identity: State variable* |  |  |  |
| $s(i)$ | Species to which school $i$ belongs | – | |
| **Bioenergetics** |  |  |  |
| *Emerging individual variables* |  |  |  |
| $P(i, t)$ | Available prey biomass to an individual of school $i$ at time step $t$ | g | 1, 2 |
| $\psi(i, t)$ | Life-stage dependent multiplicative factor of maximum mass-specific ingestion rate of school $i$ at time step $t$ | – | 2, 3 |
| $I(i, t)$ | Ingestion rate of individuals in school $i$ at time step $t$ | $g \cdot \Delta t^{-1}$ | 2, 4 |
| $E_M(i, t)$ | Mobilized energy rate of individuals of school $i$ at time step $t$ | $g \cdot \Delta t^{-1}$ | 4, 9 |
| $E_m(i, t)$ | Maintenance cost rate of individuals of school $i$ at time step $t$ | $g \cdot \Delta t^{-1}$ | 7, 9 |
| $E_P(i, t)$ | Net energy available for new tissue production | $g \cdot \Delta t^{-1}$ | 9, 10, 11, 12 |
| $\rho(i, t)$ | Proportion of net energy allocated to gonadic growth | – | 10, 11, 12 |
| $\frac{dw}{dt}(i, t)$ | Somatic growth rate | $g \cdot \Delta t^{-1}$ | 11 |
| $\frac{dg}{dt}(i, t)$ | Gonadic growth rate | $g \cdot \Delta t^{-1}$ | 12 |
| *Functional responses* |  |  |  |
| $f(P)$ | Holling's type 1 functional response to prey biomass $P$ | – | 2 |
| $\lambda([O_2])$ | Dose-response function of energy mobilization to dissolved oxygen concentration $[O_2]$ | – | 4, 5 |
| $\varphi_M(T)$ | Response of energy mobilization to temperature $T$ | – | 4, 6 |
| $\varphi_m(T)$ | Response of maintenance rate to temperature $T$ | – | 7, 8 |
| **Spatial scales and units: grid cells** |  |  |  |
| Spatial coordinates: *State variables* |  |  |  |
| $x(c)$ | Longitude of grid cell $c$ | | |
| $y(c)$ | Latitude of grid cell $c$ | | |
| Physico-chemical factors: *State variables* |  |  |  |
| $pc_k(c, t, z)$ | Value of physico-chemical factor $k$ of grid cell $c$ at time step $t$ of layer $z$ | | |
| $T(c, t)$ | Temperature of grid cell $c$ at time step $t$ | K | 4, 6, 7, 8 |
| $[O_2](c, t)$ | Dissolved $O_2$ concentration of grid cell $c$ at time step $t$ | % | 4, 5 |
| Biomass of LTL: *State variables* |  |  |  |
| $B_{LTL}(c, j, t)$ | Biomass of *LTL* group $j$ of grid cell $c$ at time step $t$ | $g$ | |

Fish schools are distributed on a horizontal spatial grid that is composed of regular cells and that covers the geographical range of the ecosystem represented. A cell c is characterized by its spatial coordinates, longitude *x*(*c*) and latitude *y*(*c*), and several other variables: (i) the vertically-distributed values (z vertical layers) of k physico-chemical factors *pc_k_*(*c,t,z*) (such as temperature *T*(*c,t,z*) or the level of oxygen saturation (%) [*O*_2_](*c, t, z*)) and (ii) the biomass of all low trophic level (LTL) groups (indexed by *j*) *B_LTL_*(*c,j,t*) that are not explicitly modeled in Bioen-OSMOSE but provided as input from coupled hydrodynamic and biogeochemical models.

Below we describe the bioenergetic sub-model that we developed to describe individual life-history and its responses to environmental variations. The individuals described with this level of detail belong to high trophic level (HTL) species, mainly fish and macroinvertebrate species.

#### 2.1.2. Individual life history description

Individual life history emerges from underlying bioenergetic fluxes which are described according to a biphasic growth model (Fig. 1) (Andersen, 2019; Boukal et al., 2014; Quince et al., 2008). The body mass-dependent energy fluxes are allocated according to physiological tradeoffs between competing processes: maintenance, somatic growth and gonadic growth. The sexual maturation of individuals relies on the concept of maturation reaction norms that depicts how the process of maturation responds plastically to variation in body growth (Heino et al., 2002; Stearns & Koella, 1986). This combination of processes mechanistically describes how somatic growth, sexual maturation and reproduction emerge from energy fluxes sustained by food intake resulting from opportunistic size-based predator-prey interactions.

On top of the biphasic growth model, individuals’ energy mobilization and maintenance energetic costs depend on dissolved oxygen concentration and temperature so that the resulting metabolic rate (the net energy available for new tissue production) and thus somatic and gonadic growth vary with these abiotic parameters in a way that conforms to the oxygen- and capacity-limited thermal tolerance theory (OCLTT; Pörtner, 2001) and more generally to thermal performance curves (TPC; Angilletta, 2009).

In the following description, energetic fluxes are expressed in somatic mass unit equivalents under the assumption that the ratio of energy density between somatic and gonadic tissues *η* is independent of size. All the parameters of the bioenergetic and life-history sub-model are species-specific parameters except one parameter is constant across species, namely the Boltzmann constant (Table 2).

**Table 2:** Species-specific parameters of the bioenergetic and life-history sub-model. (Δ*t*: time step duration). One parameter is constant across species, namely the Boltzmann constant (*k_b_* = 8.62 *e*^-5^*eV.K*^-1^).

| Symbol | Description | Units | Equations | Source |
| --- | --- | --- | --- | --- |
| Ingestion, assimilation and mobilization |  |  |  |  |
| $R_{min}, R_{max}$ | Minimum and maximum predator to prey size ratio | – | 1 | Literature |
| $\gamma(i, j)$ | Accessibility coefficient of prey species $s(j)$ to predator species $s(i)$ according to life stage | – | 1 | Literature |
| $I_{max}$ | Maximum mass-specific ingestion rate | $g \cdot g^{-\beta} \cdot \Delta t^{-1}$ | 2 | Calibrated |
| $\beta$ | Scaling exponent of maximum ingestion rate and maintenance rate with body mass | – | 2, 7 | Assumed |
| $\theta$ | Multiplicative factor of maximum mass-specific ingestion rate at larval stage | – | 3 | Estimated<sup>1</sup> |
| $a_l$ | Age at the end of the fast growth period | y | 3 | Literature |
| $\xi$ | Assimilation efficiency | – | 4 | Literature |
| $c_{O,1}$ | Asymptote of energy mobilization dose-response function to dissolved oxygen saturation | – | 5 | Estimated<sup>2</sup> |
| $c_{O,2}$ | Slope of energy mobilization dose-response function to dissolved oxygen saturation | % | 5 | Estimated<sup>2</sup> |
| $\varepsilon_M$ | Increasing activation energy of the energy mobilization temperature function | eV | 6 | Estimated<sup>3</sup> |
| $\varepsilon_D$ | Declining activation energy of the energy mobilization temperature function | eV | 6 | Estimated<sup>3</sup> |
| $T_p$ | Temperature of peak value in energy mobilization | K | 6 | Estimated<sup>3</sup> |
| $\Phi$ | Normalization constant of the energy mobilization temperature function | – | 6 | Estimated<sup>3</sup> |
| Maintenance |  |  |  |  |
| $c_m$ | Mass-specific maintenance rate | $g \cdot \Delta t^{-1}$ | 7 | Literature |
| $\varepsilon_m$ | Activation energy of the maintenance temperature function | eV | 8 | Estimated<sup>3</sup> |
| Maturation |  |  |  |  |
| $m_0$ | Intercept of the maturity reaction norm | cm | 13 | Estimated<sup>1</sup> |
| $m_1$ | Slope of the maturity reaction norm | cm · y<sup>-1</sup> | 13 | Estimated<sup>1</sup> |
| New tissue production |  |  |  |  |
| $\eta$ | Energy density ratio between somatic and gonadic tissue | – | 10, 12 | Literature |
| $k$ | Allometric length-somatic mass relationship coefficient | $g \cdot cm^{-\alpha}$ | | Estimated<sup>4</sup> |
| $\alpha$ | Allometric length-somatic mass relationship exponent | — | | Estimated<sup>4</sup> |
| Reproduction |  |  |  |  |
| $r$ | Gonado-somatic index | — | 10 | Estimated<sup>1</sup> |
| $w_{egg}$ | Egg mass | $g$ | 14 | Estimated<sup>5</sup> |
| $sp(t)$ | Spawning seasonality expressed as a fraction of the gonad energy available at the time step $t$ | — | 14 | Literature |
| $n_{s(i)}$ | Number of new schools produced at each time step of the breeding season for the species $s$ of school $i$ | # | 15 | Arbitrary |

<sup>1</sup> from Size Maturity Age Length Key (SMALK) data (see Supporting Information S2.1)
<sup>2</sup> from ecophysiological data (see Supporting Information S2.4)
<sup>3</sup> from a combination of SMALK and thermal tolerance data (see Supporting Information S2.3)
<sup>4</sup> from length-mass data
<sup>5</sup> from fecundity data (see Supporting Information S2.5)

#### 2.1.3. Ingestion, assimilation and mobilization

For an individual in school *i*, the ingested food *I*(*i, t*) at time step *t* is described by a Holling’s type 1 functional response (Holling, 1959) that depends on its somatic mass *w*(*i, t*) (Christensen & Walters, 2004; Holt & Jorgensen, 2014; Shin & Cury, 2004) in two ways. First, it determines the prey biomass *P*(*i, t*) available to an individual of school *i*. All other fish schools and LTL organisms (from the forcing biogeochemical model) that are present in the same grid cell *c*(*i,t*) are potential prey if their body size is compatible with a minimum *R_min_* (Shin & Cury, 2004) and a maximum *R_max_* predator to prey size ratio based on individual total length 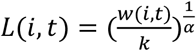 (Travers et al., 2009) so that:

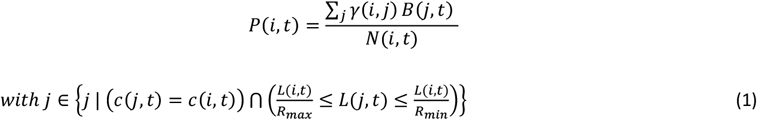

where *k* and *α* are the allometric length-somatic mass relationship coefficient and exponent, respectively, *γ*(*i,j*) is the accessibility coefficient of potential prey school *j* to school *i* that is essentially determined by the position in the water column of species relative to species according to their life stage, and is the biomass of prey school at time step *t*. The maximum possible food ingestion rate scales with the mass with a scaling exponent *β*. The ingested food can then be written as:

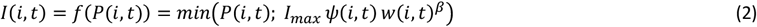

with *I_max_* the maximum ingestion rate per mass unit at exponent *β* (or mass-specific maximum ingestion rate) of individuals in school *i* and *ψ*(*i, t*) a multiplicative factor that depends on their life stage such that:

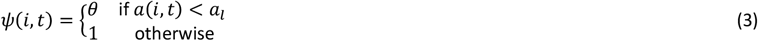

where *a_l_* is the age at the end of an early-life fast-growth period (e.g., larval period or the larval and post-larval period, defined according to data availability, see Supporting Information S2) and *θ* a multiplicative factor accounting for higher mass-specific ingestion rate at this stage. A portion of the ingested food *I*(*i, t*) is assimilated, (1 – *ξ*) being lost due to excretion and feces egestion.

Reserves are not modeled in Bioen-OSMOSE: the assimilated energy is directly mobilized. The difference between assimilated and mobilized energy depends on oxygen and temperature conditions (Fig. 2). Mobilized energy *E_M_*, referred to as active metabolic rate in the ecophysiology literature, fuels all metabolic processes such as maintenance, digestion, foraging, somatic growth, gonadic growth, etc... The mobilization of energy relies on the use of oxygen to transform the energy held in the chemical bonds of nutrients into a usable form, namely ATP (Clarke, 2019). In consequence, the maximum possible energy mobilized depends (i) directly on dissolved oxygen saturation that sets up an upper limit to mobilization at a given temperature and (ii) as temperature increases, on the capacity of individuals to sustain oxygen uptake and delivery for ATP production. The mobilized energy rate *E_M_* is thus described by:

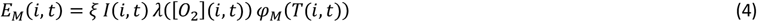

with *λ*([*O*_2_](*i,t*)) and *φ_M_*(*T*(*i,t*)) being the mobilization responses to dissolved oxygen saturation [*O*_2_](*i, t*) = [*O*_2_](*c*(*i, t*)) and temperature *T*(*i, t*) = *T*(*c*(*i, t*)), respectively, encountered by school *i* in the grid cell *c*(*i, t*). These are scaled between 0 and 1 such that, in optimal oxygen saturation and temperature conditions, all assimilated energy *E_M_*(*i,t*) = *ξ I*(*i,t*) can be mobilized and, in suboptimal conditions, only a fraction of assimilated energy can be mobilized *E_M_*(*i, t*) < *ξ I*(*i, t*).

**Figure 2:**
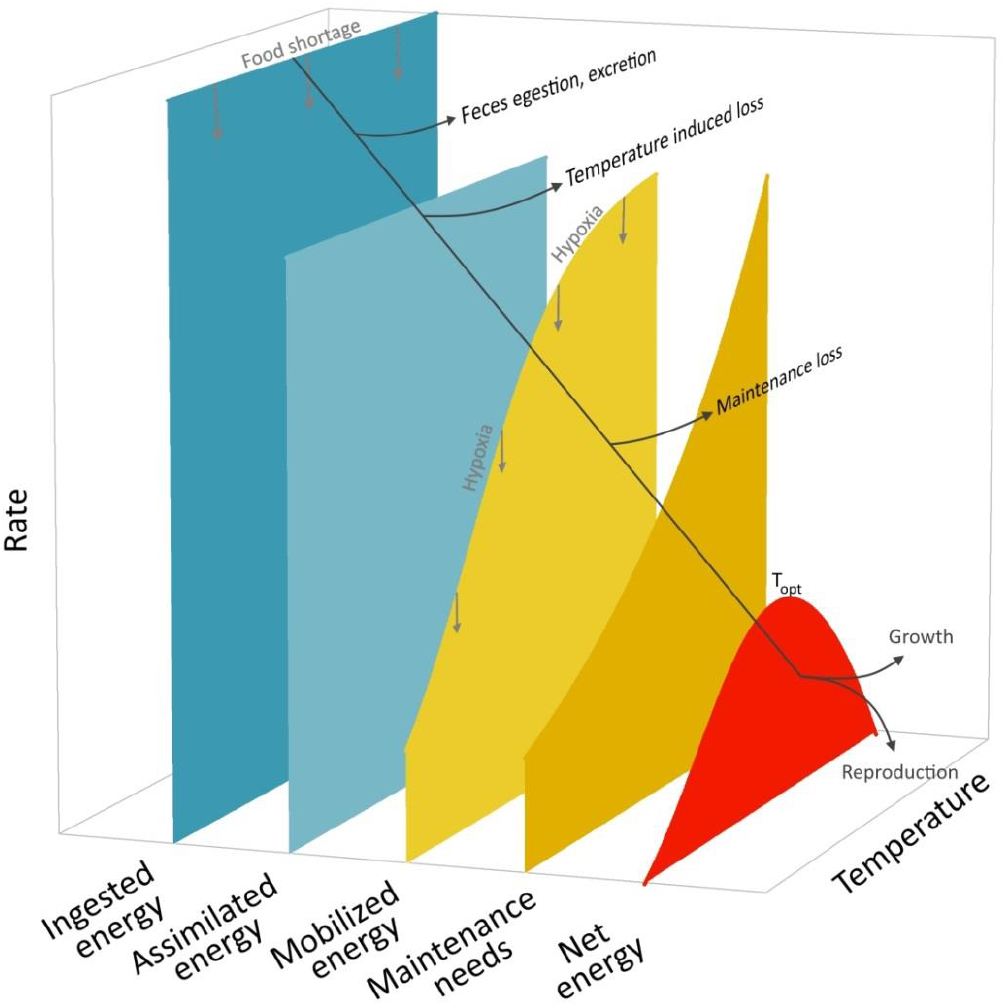
Thermal responses of the bioenergetic fluxes from ingestion to tissue growth in Bioen-OSMOSE. The net energy rate dome-shaped curve (in red) conforms to the OCLTT theory and the principle of TPC. Food shortage impacts ingested energy and downstream fluxes. Hypoxia impacts mobilized energy and downstream fluxes. The maximum of the net energy rate (red curve) is called *T_opt_* hereafter.

More precisely, the effect of dissolved oxygen is described by a dose-response function *λ*(·) (Thomas et al., 2019) which increases with the saturation of dissolved oxygen:

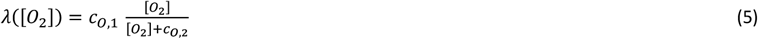

with parameters *c*_*o*,1_ and *c*_*o*,2_ the asymptote and the slope of the dose-response function. The effect of temperature *φ_M_*(·) is such that first, energy mobilization increases with temperature according to an Arrhenius-like law due to chemical reaction rate acceleration until reaching limitation in individuals’ ventilation and circulation capacity. Hence, oxygen uptake and delivery for energy mobilization saturates or even decreases at high temperatures, potentially due to temperature dependence of the rate of enzyme-catalyzed chemical reactions (Arcus et al., 2016) or enzyme denaturation (Pawar et al., 2015). This effect is described according to the Johnson & Lewin (1946) model (Pawar et al., 2015):

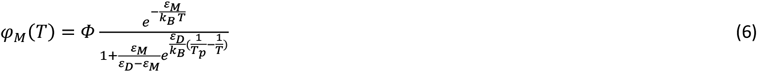

with *k_B_* the Boltzmann constant, *ε_M_* the activation energy for the Arrhenius-like increase in mobilized energy with temperature *T* before reaching its peak value at *T_p_, ε_D_* the activation energy when the energy mobilization declines with *T* above *T_p_*, and 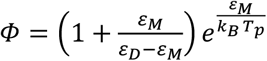 a standardizing constant ensuring that *φ_M_*(*T_p_*) = 1.

#### 2.1.4. Maintenance

The mobilized energy fuels all metabolic processes starting in priority with the costs of maintenance of existing tissues which is often referred to as the standard metabolic rate in the ecophysiology literature. Here, we also include in the maintenance costs, the routine activities of individuals, including foraging and digestion, so that they are actually best compared to the routine metabolic rate in the ecophysiology literature. The maintenance costs are explicitly modeled to describe the share of mobilized energy between maintenance and the production of new tissues (Charnov et al., 2001; Holt & Jorgensen, 2014), with precedence of the former over the latter, as well as to link mechanistically starvation mortality to energetic starvation when neither mobilized energy nor gonad energy reserves can cover the costs of maintenance (see next section on new tissue production for more details). The maintenance energy rate *E_m_* scales with the individual’s somatic mass *w*(*i,t*) with the same exponent *β* as the maximum ingestion rate. The maintenance rate also increases with the temperature experienced by individuals according to the Arrhenius law (Brown et al., 2004; Gillooly et al., 2002; Kooijman, 2010) and can be described as:

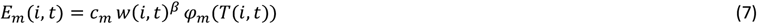

with *C_m_* the mass-specific maintenance rate and *φ_m_*(·) the Arrhenius function *m* defined as:

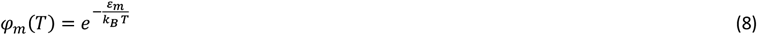

with *ε_m_* the activation energy for the increase of the maintenance rate with temperature.

#### 2.1.5. Net energy available for new tissue production

The net energy available for new tissues production *E_P_* is the difference between the mobilized energy *E_M_* and the maintenance costs *E_m_* defined as follows:

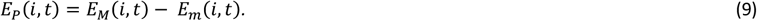

Given that the mobilized energy rate *E_M_* increases at a lower rate than the maintenance rate *E_m_* close to the species preferred temperature, it results that, all other things being equal, the emerging relationship between the net energy rate *E_P_* (and thus somatic and gonadic growth, see next section) and temperature is dome-shaped and conforms to the OCLTT theory and the principle of TPC (red curve in Fig. 2).

#### 2.1.6. New tissue production: somatic and gonadic growth

The net energy *E_P_* contributes to the production of new tissues with a proportion *ρ* being allocated to the gonadic compartment *g*(*i,t*) and a proportion (1 – *ρ*) to the somatic one *w*(*i,t*). This proportion depends on the sexual maturity status of the schools’ individuals and their somatic mass *w*(*i, t*). Before sexual maturation, i.e., when the maturity status *m*(*i,t*)=0, *ρ* is equal to 0 and, after maturation, i.e., when *m*(*i,t*)=1, *ρ* is determined such that the annual mean gonado-somatic index of individuals 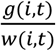 is constant throughout their adult life-stage and equal to *r* (Boukal et al., 2014; Lester et al., 2004; Quince et al., 2008):

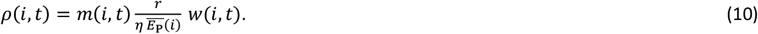

where, *η* is the ratio of energy density between somatic and gonadic tissues, 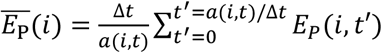 is the average net energy available per time step to individuals of school *i* since their birth, with Δ*t* being the duration of a time step. Eq. 10 differs from a deterministic continuous time version of the same model (Boukal et al., 2014; Lester et al., 2004; Quince et al., 2008) where the current net energy *E_P_*(*i,t*) would be used instead of the average 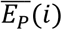. The averaging in a stochastic discrete time individual-based model such as Bioen-OSMOSE ensures a smooth increase of the proportion *p* as individuals grow by dampening strong variations in *E_P_*(*i,t*) and thus in *ρ*(*i,t*) due to the stochasticity of prey encounters and hence of the ingested energy *I*(*i,t*).

According to the definition of *ρ*, all net energy *E_P_* is allocated to somatic growth before maturation and it is shared between somatic and gonadic growth after, with the proportion *ρ* allocated to gonads increasing with somatic mass (Boukal et al., 2014), which limits somatic growth as individuals become bigger. In case the mobilized energy *E_M_* cannot cover the maintenance costs *E_m_*, i.e., when *E_P_* < 0, new tissue production is not possible and the gonadic compartment *g(i,t)* is resorbed to provide energy for sustaining maintenance. Somatic growth is then defined as follows:

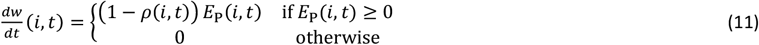

and gonadic growth as:

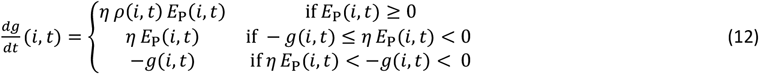

where the second and third conditional formulas account for maintenance coverage by energy reserves contained in gonads. In the former case, gonads’ energy can fully cover maintenance costs but in the latter it cannot, so that individuals undergo energetic starvation and incur additional starvation mortality (see Supporting Information S3).

#### 2.1.7. Maturation

Age and size at maturation vary strongly between individuals due to phenotypic plasticity. This plasticity in maturation is modeled by a deterministic linear maturation reaction norm (LMRN) that represents all the age-length combinations at which an individual can become mature (Stearns, 1992; Stearns & Koella, 1986). In this framework, individuals become sexually mature when their growth trajectory in terms of body length intersects the LMRN. The maturity status *m*(*i,t*) of individuals of school at time step is thus described as:

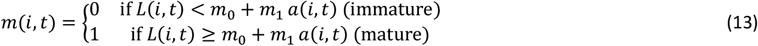

with *m*_0_ and *m*_1_ the intercept and slope of the LMRN, respectively.

#### 2.1.8. Reproduction

Mature individuals spawn during the breeding season, then a gonad portion is used to release eggs, what is represented by a gonad portion released *sp*(*t*′) above 0. The sex-ratio is assumed to be 1:1 for all species and the number of eggs produced by school *i* at time *t* is defined as follows:

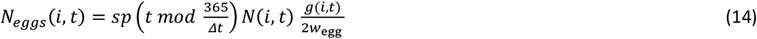

with 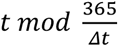 giving the time of the year at time step *t* for a time step size of Δ*t*, and *w_egg_* the mass of an egg.

At each time step *t* of the breeding season (i.e., *t* for which 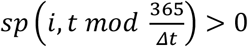, *n*_*s*(*i*)_ new schools are produced by species *s*(*i*), with the number of eggs, and thus individuals, per new school *i*′ calculated as follows:

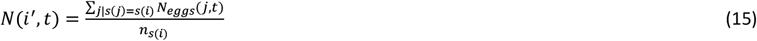

with ∑_*j*|*s*(*j*)=*s*(*i*)_ *N_eggs_*(*j, t*) the total number of eggs produced by schools of species *s*(*i*) at time step *t*, age of offspring set to 0, *a*(*i*′,*t*) = 0, their somatic mass to the mass of an egg, *w*(*i*′,*t*) = *w_egg_*, and their gonadic mass to 0, *g*(*i*′,*t*) = 0. The new schools are released randomly depending on the specific larvae habitat map.

#### 2.1.9. Mortality

At each time step, a school experiences several mortality sources. The total mortality of a school *i* is the sum of predation mortality caused by other schools, starvation mortality, fishing mortality, and additional mortalities (i.e. larval, senescence, diseases, and non-explicitly modeled predators). For a school *i*, the starvation mortality results from the encountered food, the environmental abiotic variables and its maintenance rate. If the mobilized energy *E_M_* covers the maintenance costs *E_m_*, there is no starvation. However, if the mobilized energy *E_M_* is lower than the maintenance costs, the school *i* has an energetic deficit. In this case, the gonad *g*(*i, t*) is used as a reserve. In case the gonad content does not cover the exceeding maintenance costs, the school *i* faces starvation mortality proportionally to the remaining energetic deficit. The equations and details about all mortality processes are in Supporting Information S3.

### 2.2. Application to the North Sea ecosystem: Bioen-OSMOSE-NS

#### 2.2.1. Area and explicit species

The Bioen-OSMOSE-NS area includes the North Sea (ICES area 4) and the eastern English Channel (ICES area 7d), excluding the area deeper than 200m (notably the Norwegian Trench), and extends from 49° N to 62° N and from 4° W to 8.5° E (red delimitation in Fig. 3). The model covers the area by 632 regular cells of 0.25° x 0.5°. 16 HTL species are modeled explicitly, accounting for 89% of the total fisheries landings in the area over the period 2010-2017 (ICES 4abc and 7d) and more than 90% of the scientific North Sea International Bottom Trawl Survey (NS-IBTS-Q1, DATRAS) catches. There are five pelagic species, seven demersal species, three benthic flatfish and one shrimp functional group. The species and their input parameters are listed in Supporting Information S4, Table S3, and the data sources, references, and/or methodology to estimate these parameters are presented in the Supporting Information S6. The species spatial distributions are described by presence/absence maps, and informed per life stages (egg-larvae, juvenile, and adult) whenever information was available (Supporting Information S7). As individuals are represented in a 2D horizontal environment, a predator-prey accessibility matrix *Γ*, used to determine the accessibility coefficient *γ*(*i,j*) (see Eq. 1 and Table 2), is defined according to the vertical distribution overlap between potential predator and prey species possibly per life stage (Supporting Information S4, Table S5). The gonad portion released *sp*(*t*) is estimated from the seasonality of eggs’ release (see Supporting Information S2.5). The seasonality of eggs’ release data are taken from the literature and presented in Supporting Information S8. The fishing mortality rates are size-dependent due to fisheries size-selectivity. The calibrated maximum fishing mortality rates *F_max_* are in Supporting Information S4, Table S3. The species-specific fishing selectivity curves are in Supporting Information S11. The larval and additional mortality are in Supporting Information S4, Table S3.

**Figure 3:**
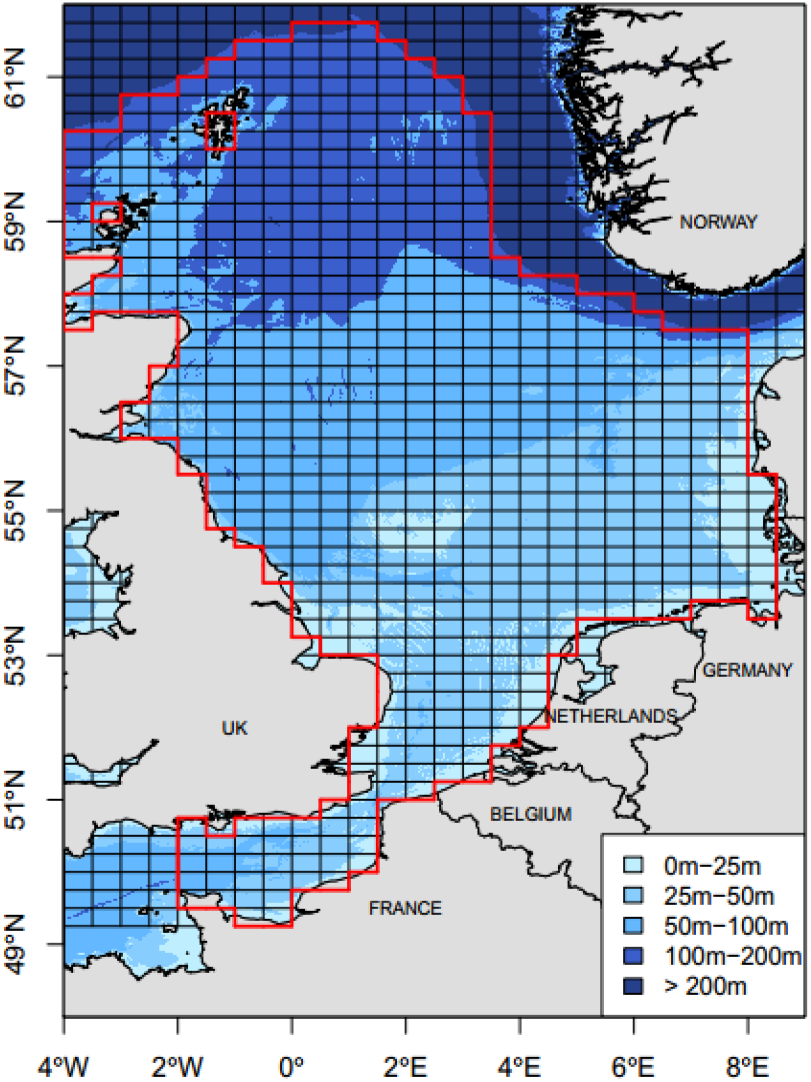
Case study area map. The Bioen-OSMOSE-NS area is delimited with the red line. The area is divided in 632 regular cells of 0.25° x 0.5° delimited with black lines.

#### 2.2.2. Forcing variables: low trophic levels and physical variables

The Bioen-OSMOSE model is forced by temperature and oxygen variables and by LTL biomass fields. The forcing data come from the regional biogeochemical model POLCOMS-ERSEM applied to the North Sea ecosystem (Butenschön et al., 2016). The modeled period is 2010-2019. There are five pelagic (micro-phytoplankton, diatoms, heterotrophic flagellates, micro-zooplankton, meso-zooplankton) and three benthic (suspension feeders, deposit feeders and meiobenthos) LTL groups (Supporting Information S5): the biomass of the former is available in three dimensions and therefore integrated vertically, while the biomass of the latter is available in two-dimensions. Two other groups of large and very large benthos are set as homogeneous prey fields in space and time due to the absence of data and model output for these LTL groups. For the temperature and oxygen variables, their values are integrated over the 43 vertical layers of POLCOMS-ERSEM to force pelagic and demersal HTL species. Only the values in the deepest layer are used for benthic species. Monthly maps for each LTL group and temperature and oxygen variables are shown in Supporting Information S9.

#### 2.2.3. Calibration

The model is calibrated, i.e., parameters for which an independent estimator is unavailable are estimated, using maximum likelihood estimation based on an optimization method adapted to high-dimensional parameter space, namely an evolutionary algorithm available in the package calibraR in R (Oliveros-Ramos & Shin, 2016). The algorithm explores the space of unknown parameters (referred to as “calibrated” in Supporting Information S4, Table S3) so as to maximize the likelihood obtained by comparing model outputs to observed data. Data used to calibrate Bioen-OSMOSE-NS are fisheries landings (ICES, 2019a), size-at-age from scientific surveys (NS-IBTS-Q1, North Sea International Bottom Trawl Survey (2010-2019), available online at http://datras.ices.dk) and estimated biomasses for assessed species (ICES, 2016, 2018a, 2018b, 2018c, 2019b). The discard rate of assessed species is low except for dab and plaice: the data used as landings and biomass for these species includes estimated discards from stock assessments. The biomasses estimated for stocks entirely located within the study area are directly used (herring, sandeel, sprat, sole, and whiting). For stock with a wider distribution than the study area, the biomass data is taken proportional to total stock biomass according to the ratio between landings in the study area and total landings (mackerel, norway pout, plaice, saithe, cod, haddock, dab, hake). There is no biomass target value for unassessed species (horse mackerel, grey gurnard, hake, shrimp). The calibration is performed for an average state of the ecosystem for the period 2010-2019 by using observed data averaged over the period as target values (Supporting Information S10). The calibration is run using four phases with a new set of parameters to be estimated added at each phase for better convergence of the optimization: the first phase calibrates the LTL group accessibility coefficients only (Supporting Information S5), the larval mortalities are added for the second phase (Supporting Information S4, Table S3), the maximum ingestion rates are added on phase three (Supporting Information S4, Table S3), and the maximum fishing mortality rates and the additional mortality rates are added in the last phase (Supporting Information S4, Table S3).

The calibrated configuration is run for 80 years. The first 70 years is the spin-up period, a period during which the system stabilizes. The results presented hereafter are the years after the spin-up period. 28 replicates of the model are run with the same parameterization to account for Bioen-OSMOSE stochasticity.

## 3. Results and discussion

In this paper, we present the Bioen-OSMOSE framework with its first application to the North Sea ecosystem, involving the coupling of the POLCOMS-ERSEM model for the physical and LTLs model with the HTLs Bioen-OSMOSE model. The North Sea trophic network has been intensively studied and modeled, either considering the whole ecosystem (Blanchard et al., 2014; Cormon et al., 2016; Heath, 2012; Lewy, 2004; Mackinson & Daskalov, 2007) or part of it (including the English Channel) (Girardin et al., 2018; Stäbler et al., 2016; Travers-Trolet et al., 2019).

This is the first time that the Bioen-OSMOSE model is used, i.e the OSMOSE model (Shin & Cury, 2004) augmented with a mechanistic description of the emergence of life history from underlying bioenergetics and its response to temperature and oxygen seasonal and spatial variations. It is also, to our knowledge, the first application of a marine ecosystem model considering the impacts of physiologically-induced trait changes in response to food, oxygen, and temperature at the individual, population, and community levels.

### 3.1. Model evaluation

The calibration procedure allowed us to estimate unknown parameters to obtain a model configuration that fairly accurately represents the North Sea ecosystem. Particular attention was paid to obtaining satisfactory results for indicators at different biological levels, the final result being a compromise between indicators used as target (size-at-age, catch and biomass) and emerging variables (maturation and diet).

#### 3.1.1. The individual level: size structure and maturity ogives

The simulated mean sizes-at-age correctly reproduce the observed ones (Fig. 4), supporting the credibility of the growth process described by the new bioenergetic sub-model. The Von-Bertalanffy-like shape and the indefinite growth are two realistic properties reproduced with our model. As observed in the data, it can be noted that growth is faster during the first years of life and the sizes at older ages slowly tend to an infinite size. The simulated and observed sizes-at-age interquartile ranges overlap for almost all age classes of the species. The simulated size hierarchy between species is consistent with the observed one, which is a key expected property for a size-based model.

**Figure 4:**
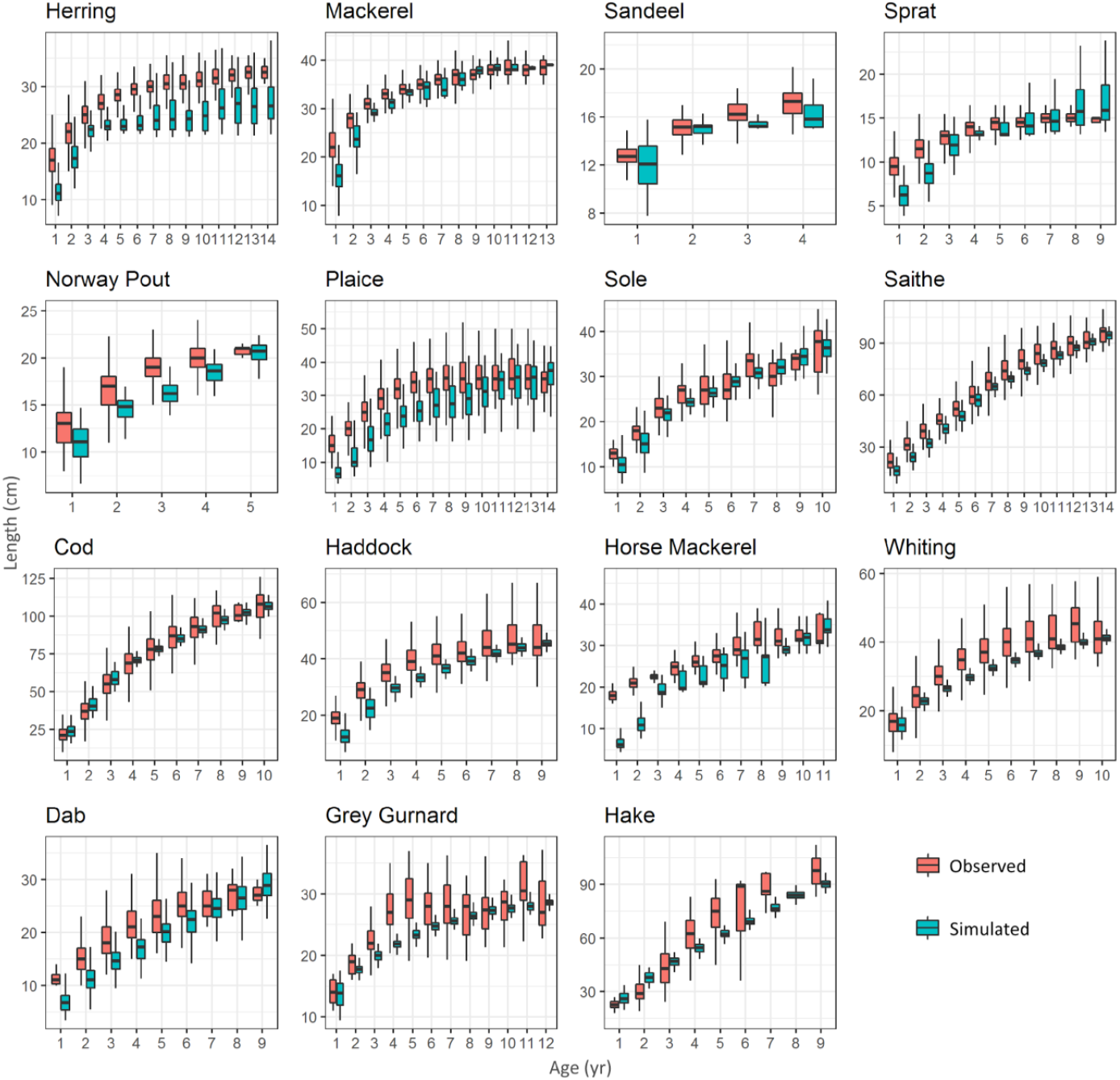
Boxplot of size-at-age per species for observed (pink) and simulated (blue) individual data. Horizontal bars represent the first, second and third quartiles. The whiskers’ extremities represent 1.5 times the interquartile space (the distance between the first and third quartile). The shrimp group was not represented in this graph, as available observed data are not sufficiently taxonomically resolved to be relevant for this functional group.

The simulated sizes-at-age 1 have generally the poorest fit to observed data. Size-at-age class 1 partly inherits uncertainties linked to size at hatching and to the growth rate at very early stages, notably the larval one, that is imperfectly accounted for by the multiplicative factor of maximum mass-specific ingestion rate for the larvae at larval stage *θ.* In addition, growth during the first year is mainly driven by food limitation implying that the size-at-age class 1 is the result of a complex model adjustment between growth rate, competition and prey accessibility.

The variance in size-at-age differs between observed and simulated data, mainly for demersal species. The observed variance in sizes-at-age is the result of macro-environmental variations, i.e, in the abiotic environment (Brown et al., 2004; Gislason et al., 2010; Thomas et al., 2019) and food availability (Brosset et al., 2016), micro-environmental variations, i.e., in undetectable or unaccounted for environmental conditions, and genetic variability in energy allocation inducing variability in size-at-age (Enberg et al., 2012). In contrast, the variance of simulated size-at-age only results from macro-environmental variations. Thus, the species with observed variance higher than the simulated variance is because genetic and micro-environmental variances are not modeled here.

Comparison of observed and simulated age and size maturity ogives demonstrate the ability of Bioen-OSMOSE to correctly reproduce maturation patterns (Fig. 5). The simulated mean age at first maturation perfectly matches that observed for three species (cod, norway pout, and whiting). In observed data, age is given with yearly resolution. Therefore, we consider that a correct pattern is obtained for seven additional species for which the difference between simulated and observed age at maturity is less than one year (grey gurnard, haddock, hake, herring, mackerel, plaice and sole). The worst deviation is obtained for sprat and saithe with simulated maturation occuring at later ages and larger sizes than that observed. Saithe and sprat have lower mean sizes at early ages than observed ones (Fig. 4) which can explain the late simulated maturation. However, both species have simulated maturation ages that stand within observed ranges, being lower than the upper bounds of observed maturation ages, namely 9 years for saithe in the North Sea (Cohen et al., 1990) and 4 years for sprat (Ojaveer and Aps, 2003) in the Baltic Sea (only mean values were reported for the North Sea population).

**Figure 5:**
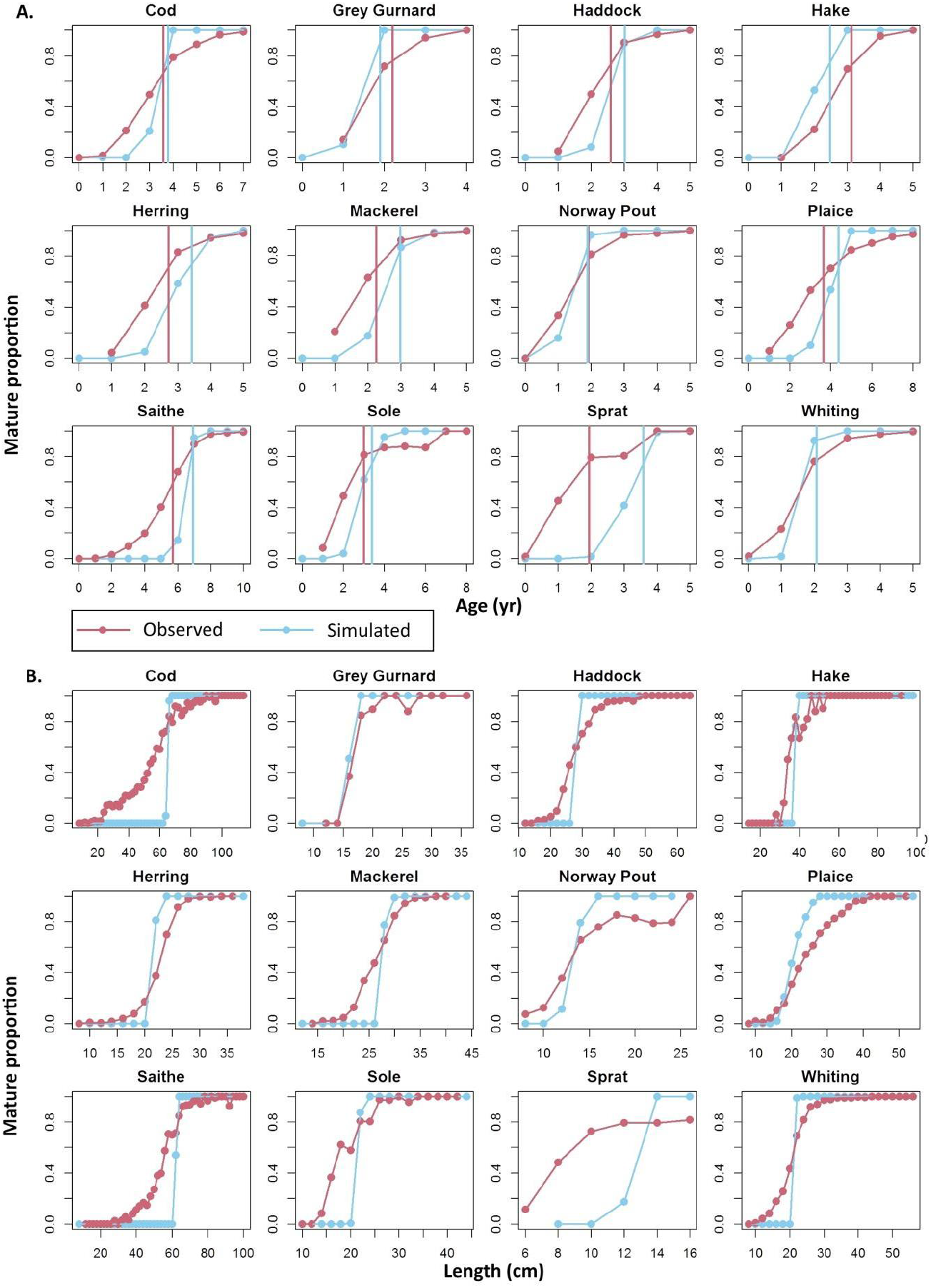
Age (A) and size (B) maturity ogives per species for observed (pink) and simulated (blue) data. Results are shown for species for which there is enough data to estimate and plot the observed age and size maturity ogives. Age data have a yearly resolution and size data a 2-centimeter resolution. The simulated (blue) and observed (pink) mean age at maturity are represented by vertical lines (A). The mean size at maturity is not represented. The observed size maturity ogive is not strictly increasing and does not allow a reliable estimation of the maturation size.

The use of LMRNs improves the description of the variability in the maturation process compared to the use of fixed age or size at maturity, as is most commonly done in marine ecosystem models, such as previously in the OSMOSE framework (Shin & Cury, 2004) or in other models (Audzijonyte et al., 2019). Consequently, individuals from the same size or age class do not necessarily have the same maturity state (Fig. 5), which increases the realism of the life cycle description. For the majority of the species, the slope of simulated age and size maturity ogives is higher than the observed one, meaning that the observed maturation process is more variable than in simulations. As for size-at-age, part of the observed variability in maturation is determined by genetic and/or micro-environmental variability (Law, 2000; van Wijk et al., 2013) and is not modeled here.

#### 3.1.2. The population level: fisheries catches and species biomass

Given the high statistical confidence in catch data, a greater weight was given to the corresponding likelihood component in the calibration process which resulted in reaching targets, i.e., simulated catches were within the range of observed values, for the majority of species (Fig. 6A). Plaice and dab are the two species for which the simulated catches were the farthest from their targets. Plaice and dab are two by-catch species that are largely discarded. These discards are estimated to be up to 40% for plaice and 90% for dab (ICES, 2018c). The catch data used as target here is reconstructed from the landings and discards estimates, which, in case of overestimation of discards, could explain the discrepancy between the target and the simulated values. The poor fit of plaice could also be partially explained because the migrations between the Eastern and Western English Channel stocks are not taken into account in Bioen-OSMOSE-NS (ICES, 2021).

**Figure 6:**
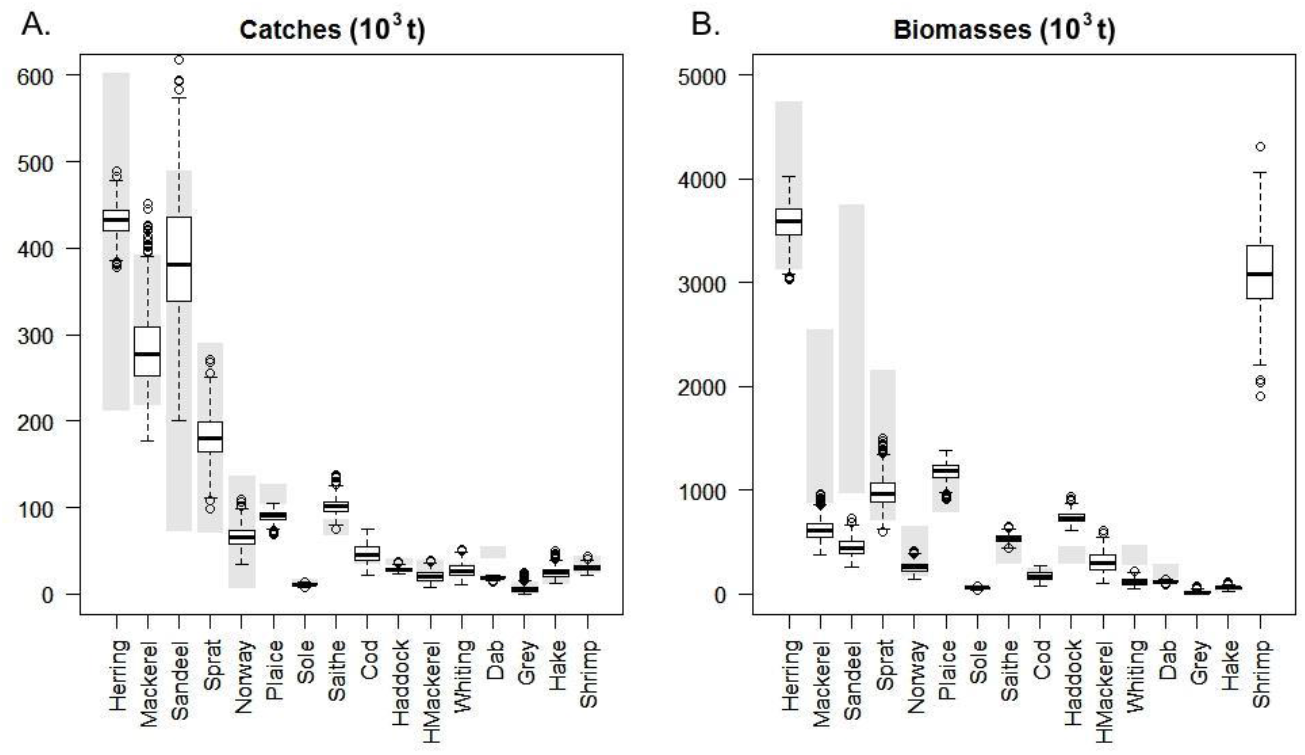
Fisheries catches (A) and biomasses (B), in thousand tons, per species for stock assessment estimates and simulated data. The boxplots represent the simulated data for 28 replicated simulations (stochastic model) for the catches and biomasses per species, with the first, second and third quartiles represented horizontally in each plot. The gray bars show the minimal and maximum values observed for catch and biomass estimates from stock assessment for the 2010-2019 period. The species without gray bars for biomasses are not assessed in the area.

The simulated biomasses are within acceptable ranges (Fig. 6B) and the resulting dominance ranking between species groups respects the ranking based on stock assessment estimations: small pelagic fish are the dominant group with herring as the main species. Demersal species have lower biomasses with saithe and haddock as the most abundant ones. Flatfish are the minority group in the system, which is dominated by plaice.

The simulated biomasses of mackerel and sandeel are underestimated compared to the stock assessment biomass estimates (Fig. 6B). Mackerel is a widely distributed stock in the North East Atlantic area and our biomass estimate in the North Sea (proportional to the total biomass in the North East Atlantic according to the ratio between North Sea landings and total landings) relies on the assumption of a uniform fishing effort in the assessed area. A higher fishing effort within the North Sea would lead to an overestimation of the target biomass for the area, which is credible since the North Sea is a historically heavily fished area. An ecopath model of the North Sea (Mackinson & Daskalov, 2007) also estimated a lower biomass for this species (980 400 t). Underestimation for sandeel is more troublesome as it is a key forage species in the North Sea (Engelhard et al., 2014), for which the stock assessment is considered to be very detailed with 7 stocks in the area (ICES, 2016). The fact that the Bioen-OSMOSE-NS model does not describe the peculiar overwintering behavior of this species, which buries itself in sand and thus is less vulnerable to fishing and predation in winter (Henriksen et al., 2021), may explain the underestimation of its biomass. The sandeel may also be over-consumed by higher tropic level species in our model, indicating a missing forage species or an over-consumption of sandeel over LTL forced prey.

In Bioen-OSMOSE-NS, flatfish are represented by only the three main species of the North Sea ecosystem. However, there are other flatfish species each with low biomass levels *(Scophthalmus maximus, Microstomus kitt, Scophthalmus rhombus, Platichthys flesus*...) (NS-IBTS-Q1, DATRAS, Piet et al., 1998) but whose total biomass is not negligible (NS-IBTS-Q1, DATRAS, Mackinson & Daskalov, 2007). Thus, the overestimation of plaice biomass may compensate for the absence of these other flatfish in the model that may leave an empty trophic niche.

The high biomass of the shrimp functional group, dominated in the ecosystem by the species *Crangon crangon* and *Pandalus borealis,* may seem surprising. However, as the micro- and meso-zooplankton groups described by the biogeochemical model POLCOMS-ERSEM represent pelagic prey of sizes smaller than 0.5 cm only, we suggest that the shrimp functional group has a broader ecological role in the Bioen-OSMOSE-NS model by actually representing all LTL prey larger than 0.5 cm in the water column, whose biomass is critical to sustain the food web. These prey include demersal crustaceans with diel vertical migration such as *Crangon crangon* or *Pandalus borealus* as well as more pelagic species such as large amphipods (large *Bathyporeia elegans)* or euphausiids (*Thysanoessa sp.*, *Meganyctiphanes norvegica*).

In Bioen-OSMOSE, the diet emerging from opportunistic predation reflects the species’ relative abundances, their sizes and their spatio-temporal overlap. There are no pre-established predator-prey diet matrix in the parameterization of the model so confronting the output diets to observed ones, especially in terms of species composition, is a way to validate the model properties.

The simulated diets show patterns that are consistent with observations (Fig. 7). The model reproduces correctly observed ontogenetic diet shifts (Timmerman et al., 2020). The prey composition shifts between pelagic early-life stages (size class 0-10 cm for fish species and 0-3 cm for the shrimp group) and the older life stages for all species. There are different emerging diet patterns depending on the predator’s position in the water column. The pelagic species diet is dominated by phyto- and especially zoo-planktonic prey, which is consistent with studies on sprat and herring (De Silva, 1973; Last, 1989; Raab et al., 2012). The benthic species diet is composed of benthic LTL groups and the shrimp functional group, similarly to results obtained by an isotopic study (Timmerman et al., 2021), by plaice and sole stomach content studies for adults (Rijnsdorp & Vingerhoed, 2001) and for juveniles (Amara et al., 2001) with smaller prey for sole than for plaice and dab of the same size (Amara et al., 2001). The demersal species have an intermediate diet composition with a high degree of piscivory for the larger fish. There is a steady increase in piscivory with size, mainly for demersal species, as shown empirically for whiting, cod, saithe and haddock in the area (Robb & Hislop, 1980; Timmerman et al., 2020) but not for norway pout (Robb & Hislop, 1980). In addition, there is a significant part of benthic prey in the pelagic species diet, which correctly represents the strong pelagic-benthic coupling in this area (Giraldo et al., 2017; Timmerman et al., 2021): the pelagic piscivorous fish (mackerel and horse mackerel) also feed on benthic prey which represents half of their diet (Giraldo et al., 2017).

**Figure 7:**
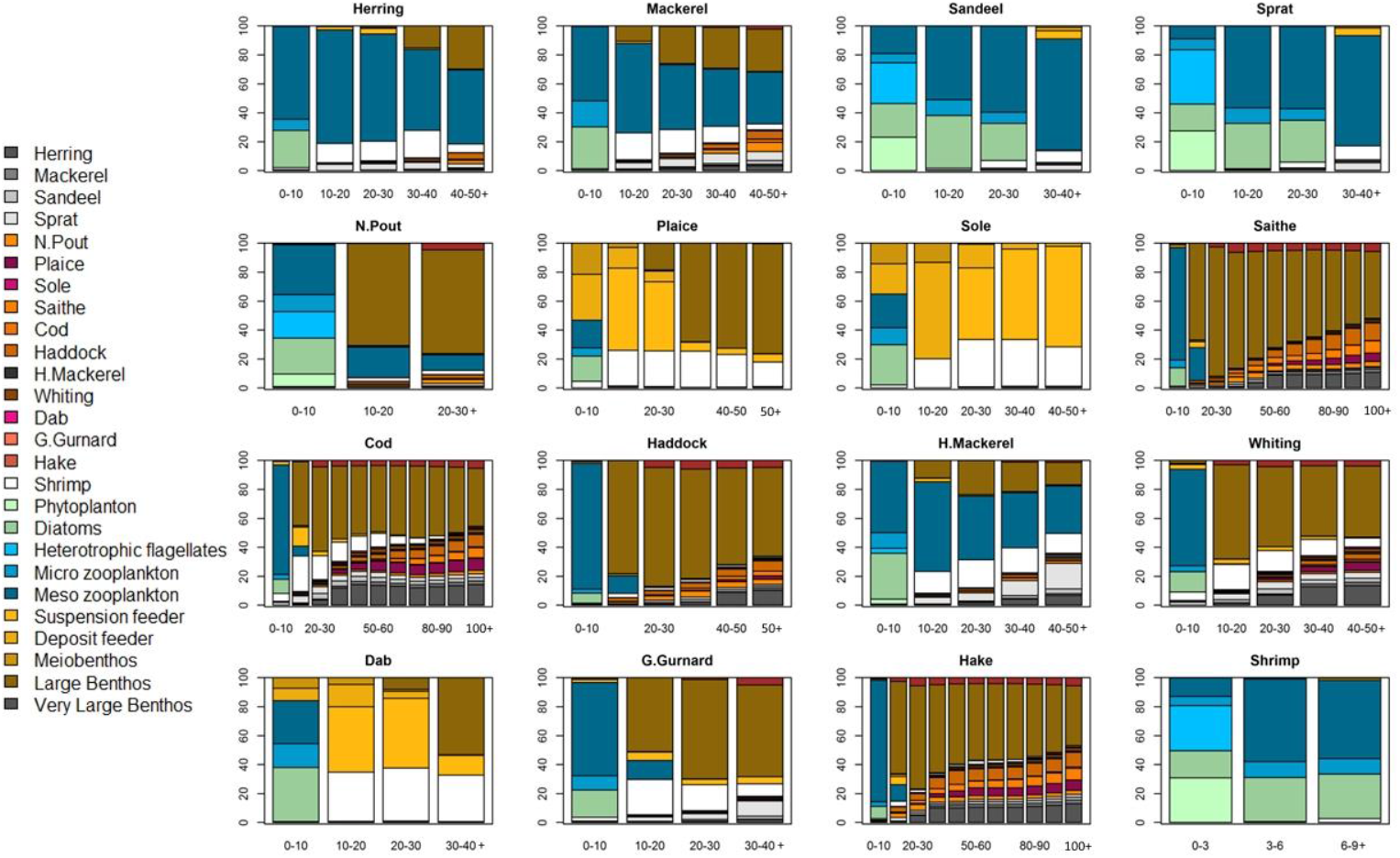
Diet in percent of biomass eaten of prey species per size class of the predator species in cm. For each predator species, the last size class (x-axis) includes all the larger individuals.

### 3.2. The physiological level: spatial pattern

New outputs and original questions emerge from the physiological responses of metabolic rates to biotic and abiotic variables and can be explored with the Bioen-OSMOSE model. The representation of emergent spatially and seasonally varying bioenergetic fluxes is an example of the new features brought by Bioen-OSMOSE that can help improve our understanding of the relationship between temperature and ecosystem dynamics, which is crucial in the context of global warming (Lindmark et al., 2022). This spatial and seasonal variability of metabolism in relation to temperature variation is often under-studied.

The simulated adult mean mass-specific net energy rate for new tissue production 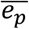, is the ratio between the population mean mass-specific net energy rate (see Eq. 9) and the weight at the exponent *β* and it drives the energy allocated to growth and reproduction. It is spatially represented as an output example (Fig. 8). This spatial representation of emerging bioenergetic fluxes highlights the high variability of mean mass-specific net energy rate for three widely distributed species in the North Sea ecosystem with contrasted thermal preferences: cod, herring and dab (quoted by increasing physiological optimum temperature *T_opt_*, defined in Fig. 2). The spatial pattern of mean mass-specific net energy is mainly explained by species thermal preferences. The species with the lowest thermal preference (cod) has a greater net energy acquisition in the northern part of the area where the water is colder on average. The opposite pattern emerges for the species with the highest thermal preference (dab). There is a better energy acquisition in the south where the average temperature is higher than in the north. A similar spatial pattern for growth rate was predicted as outputs of a single-species bioenergetic model for two thermophilic flatfish in the North Sea (Teal et al., 2012). Herring, which has an intermediate thermal preference, exhibits a more spatially homogeneous emerging mean mass-specific net energy rate. During the period described by our model (2010-2019), temperature is the main driver of spatial variability in bioenergetic fluxes for these three species. The spatial distribution of food has little or no impact on adult energy acquisition for these examples as we observe that simulated food ingestion frequently reaches the maximum at the adult stage (results not shown). Likewise, oxygen saturation has no impact on the emerging spatial pattern because oxygen saturation is not low enough to become a primary driver of bioenergetic fluxes (Vaquer-Sunyer & Duarte, 2008, Supporting Information S10).

**Figure 8:**
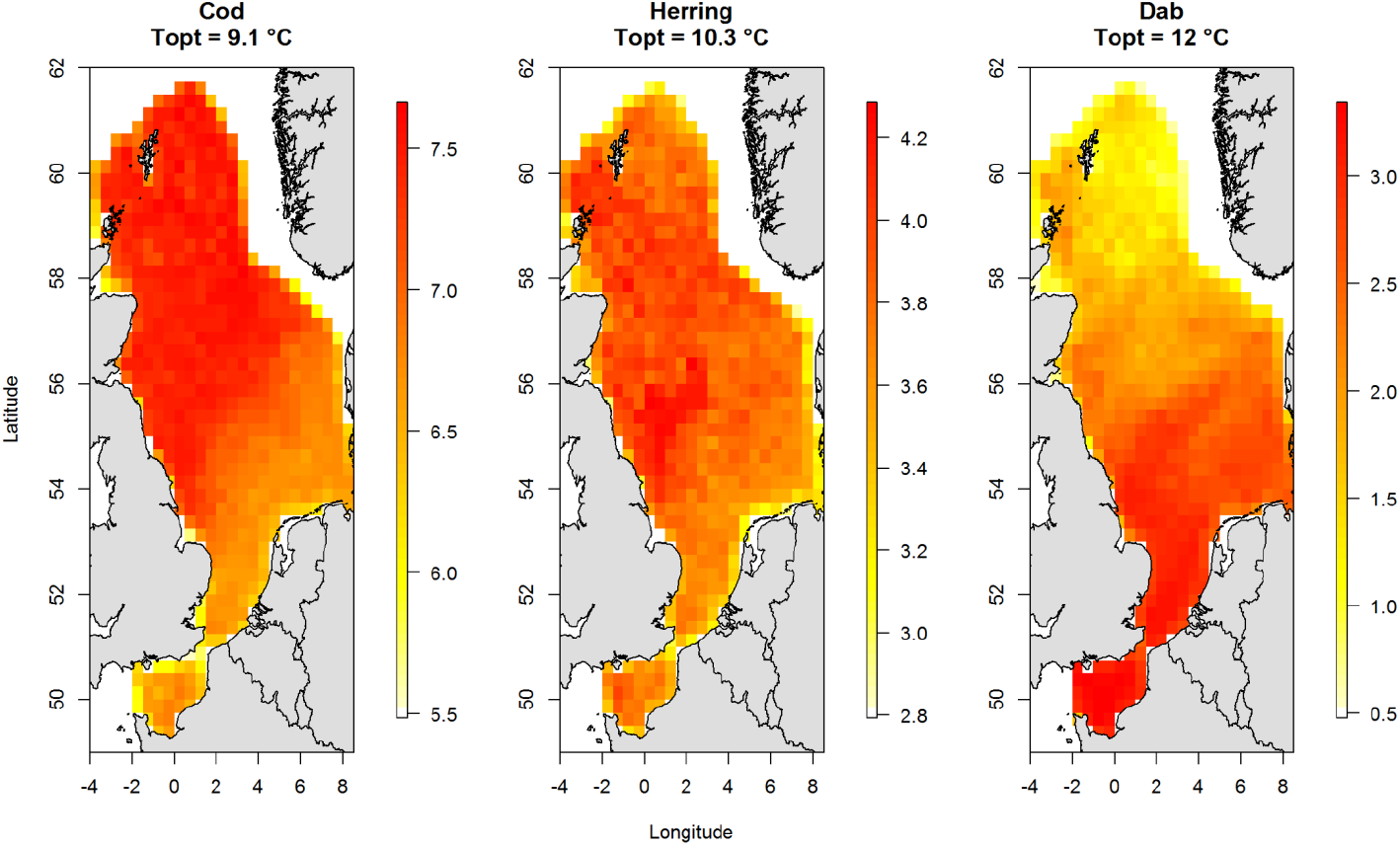
Spatial variability of the mean adult mass-specific net energy rate available for new tissue production, per model cell for cod, herring and dab. These three species are distributed over the whole modeled area although they have different optimum temperatures *T_opt_*. Cod, herring and dab mean adult mass-specific net energy available rate for new tissue production averaged over the area are 7.4, 3.3, and 2.7 *g.g^β^*, respectively. Spatial variations of this bioenergetic flux can be driven by temperature, oxygen and food variation.

## 4. Conclusion

Applying Bioen-OSMOSE to the North Sea allows demonstrating the feasibility of its parameterization for several species with different levels of available knowledge and allows evaluating the framework capabilities. Bioen-OSMOSE-NS simulates many different outcomes that convincingly reproduce observations such as biomasses, catches, sizes at age, maturation ogives, and diets. The model also produces compelling spatial responses of the bioenergetic fluxes to temperature variations.

The Bioen-OSMOSE framework is also intended to be used for hindcast or forecast simulations. Hindcasting could help disentangling the effects of temperature increase and/or oxygen depletion on the historical trends in life-history traits. Hindcasting with Bioen-OSMOSE could also be useful to understand the contribution of temperature- and oxygen-induced physiological changes in population and community dynamic alterations that were observed in past periods. Given the increasing need to reliably forecast biodiversity under future climate change scenarios, we believe that Bioen-OSMOSE will also allow improving projections of regional ecosystem dynamics by taking into account future individual-level physiological changes and their consequences at the population and community levels.

## Acknowledgements

This work has been partially funded by the BiodivErsA and Belmont Forum project SOMBEE (BiodivScen programme, ANR contract n°ANR-18-EBI4-0003-01). Alaia Morell was supported by a PhD grant from Ifremer and Région Hauts-de-France. The POLCOMS-ERSEM projections were produced with funding from the European Union Horizon 2020 research and innovation programme under grant agreement No 678193 (CERES, Climate Change and European Aquatic Resources). The authors acknowledge the Pôle de Calcul et de Données Marines (PCDM, http://www.ifremer.fr/pcdm) for providing DATARMOR storage, data access, computational resources, visualization and support services. Yunne-Jai Shin acknowledges funding support from the European Union’s Horizon 2020 research and innovation programme under grant agreement No 869300 (FutureMARES) and the Pew marine fellows programme.

## Author Contributions

Yunne-Jai Shin and Bruno Ernande conceived and supervised the project. Bruno Ernande and Alaia Morell conceived the concepts of the new model developments. Nicolas Barrier and Alaia Morell developed the code and validated the model functioning. Alaia Morell gathered the data for the model parameterization. Bruno Ernande and Alaia Morell conceived and developed the scripts for the parameter estimation. Ghassen Halouani, Morgane Travers and Alaia Morell parameterized the model. All authors interpreted the model outputs. Nicolas Barrier, Ghassen Halouani, Morgane Travers and Alaia Morell performed the model calibration. Alaia Morell wrote the first paper draft. All authors contributed critically to the revisions of the manuscript and gave final approval for submission.

## Data Archiving

The Bioen-OSMOSE-NS configuration and its associated version of the Bioen-OSMOSE model executable will be deposited on Zenodo. Model code will be available on Github. The scripts developed to estimate Bioen-OSMOSE-NS parameters are available on Github.

