## Supplementary material for "Bioen-OSMOSE: A bioenergetic marine ecosystem model with physiological response to temperature and oxygen"

### Supporting Information

Supporting Information S1: **Process overview**

Incoming flux

School initialization

LTL update, physical file update

Spatial distribution

Mortality

Ingestion, assimilation, mobilization

Maintenance

Tissue growth

Reproduction

Remove dead schools

Save outputs

Figure S1: Process execution order during a Bioen-OSMOSE time step, green processes are the biological process; white processes are the IT processes. (LTL: low trophic level)

Within each time step $t$, the following processes take place in the order described in Fig. S1. Mortality is divided into sub-time steps to take into account the simultaneity of mortality processes (see details of mortality sub-time steps within each time stepat <http://documentation.osmose-model.org/>).

1. *Income flux of biomass*

Some species might realize only part of their life cycle in the model area (e.g., ontogenetic migration in the area) or might immigrate into the model area (e.g., invasive species, climate-induced range extension), which is described by an influx of schools over time.

2. *School initialization*

All schools are initialized for the new time step, i.e., state or emerging individual variables are reset to their initial value (e.g., number of dead individuals, ingested food) or updated (e.g., age $a\left( i,t \right)$) as required.

3. *Environment and resource update*

The value of physico-chemical factors ${pc}_{k}(c,t,z)$ and the biomass of low trophic level (LTL) organisms $B_{LTL}(c,j,t)$ are updated for each cell$c$ of the spatial grid, using data from forcing files.

4. *Spatial distribution*

The model is forced with maps of species distribution (either presence/absence or densities, resulting directly from survey data or from climate niche models). Ontogenic and/or seasonal migration can be parameterized, by specifying as input of the model distinct distribution maps depending on stage, age and/or season. At each time step, schools move to an adjacent grid cell according to a random walk within their distribution map. For species that do not spend their entire lifecycle in the simulated area (due to, e.g., seasonal reproduction migration or ontogenic habitat shift), schools can emigrate from or immigrate to the model area during specified time steps, generating an outgoing or incoming flux of biomass.

5. *Mortality*

Schools experience mortality (modeled as a decrease of school abundance) related to predation, starvation, foraging, fishing and background causes (non-explicit additional predation, senescence and diseases) whose value differs between larvae stage (larval mortality) and older stages (additional mortality). Starvation mortality is directly linked to the maintenance process (see below) and relies thus on the amount of net energy at the previous time step (t-1).

6. *Ingestion, assimilation and mobilization*

Food ingestion based on the predation process on other schools and/or LTL groups, assimilation and mobilization for metabolic processes take place for each school. The school order is randomly selected.

7. *Maintenance*

Part of schools’ mobilized energy is used for the maintenance of existing tissues and basal activities such as foraging. If maintenance needs cannot be covered, the gonadic compartment is used as an energy reserve to cover them and no further metabolic process takes place. If the gonadic compartment cannot cover the maintenance costs, schools are in a state of “energetic” starvation which results in starvation mortality applied at the next time step.

8. *New tissue production: somatic and gonadic growth*

When maintenance costs are covered from mobilized energy, the remaining net energy of a school is used for new tissue production. Sexually immature individuals allocate all energy to somatic growth whereas mature ones share energy between somatic and gonadic growth.

9. *Reproduction*

During time steps corresponding to the species’ breeding period, sexually mature schools reproduce panmictically. Eggs are produced from adult gonad mass and are released in new schools distributed randomly across the larval area at the beginning of the next time step.

Schools go through these processes depending on their life-history stage. During the egg stage and the endogenous-feeding larval stage (typically the first time step of a school life), no food ingestion occurs so processes 6 to 9 do not apply and mortality of process 5 is restricted to predation and background causes. After transition to exogenous feeding, no energy is allocated to reproduction during the larval, post-larval and juvenile stages; therefore, only processes 1 to 8 are prosecuted. Schools at the adult stage go through all processes.

Processes 1 to 3 are strictly similar to those in the existing OSMOSE model and the reader is referred to its description (<http://documentation.osmose-model.org/>) for more details. Process 5 includes a new mortality term linked to foraging —and processes 6 to 10 are new and described in the main text of this article.

Supporting Information S2**: Parametrization of the new sub-model**

**1 - Estimation of the bioenergetic sub-model parameters from Size Maturity Age Length Key (SMALK) data**

The bioenergetic sub-model implemented in Bioen-OSMOSE requires ten parameters per species in input (without the oxygen and temperature parameter responses) plus three two intermediate parameters: there are two parameters for the length-somatic mass allometric relationship, nine parameters for the processes of growth and reproduction, and two parameters for the linear maturation reaction norm. Intermediate parameters are parameter that are not input parameters of Bioen-OSMOSE but that are used to estimated directly (see Supporting Information S2) or indirectly through calibration process (see Section 2.2.3) input parameters of Bioen-OSMOSE. Most of these parameters can be estimated from a common set of individual-level data classically referred to as Sex Maturity Age Length Key (SMALK) data. Sex data are used to test for sexual dimorphism in the species. If sexual dimorphism is present, only female data are retained for all parameter estimates.

To estimate the parameters $k$ and $\alpha$ of the allometric relationship between length and somatic mass, a nonlinear regression was fitted to the length-mass data.

The estimation of growth and reproduction parameters uses the reformulation by Boukal et al. (2014) of the general biphasic growth model proposed by Quince et al. (2008) and formerly derived by Lester et al. (2004) for the specific case of a constant gonado-somatic index.

This model predicts individuals’ size as a function of their age and maturity status based on the average age at maturation $a_{mat}$ in the population and their initial size $w_{1}$ estimated from the SMALK data.

Under the assumption of a constant net energy mass-specific per year during the individual lifespan and across the population spatial distribution, our model is equivalent to the model of Boukal et al. (2014). We note the annual mean mass-specific net energy acquisition rate $\bar{e_{p}}$: this parameter is equivalent to c of the Boukal model. The individual’s lifetime growth trajectory is described according to the following set of equations:

$w(a)=\left\{ \begin{aligned} \left( w_{a_{0}}^{1-\beta}+(1-\beta)\bar{e_{p}} (a-a_{0}) \right)^{\frac{1}{1-\beta}}\mathrm{if}m(a)=0 \\ \left( \frac{\eta\bar{e_{p}}}{r}-\left( \frac{\eta\bar{e_{p}}}{r}-{(w}_{a_{0}}^{1-\beta}+(1-\beta)\bar{e_{p}}(a_{m}-a_{0})) \right)\left( \frac{1}{1+(1-\beta)\frac{r}{\eta}} \right)^{a-a_{m}} \right)^{\frac{1}{1-\beta}} \mathrm{if}m(a)=1 \end{aligned} \right.$	(S1)

where $a$ is individual’s age, $a_{0}$ the age for which initial mass is observed and all other parameters are as in the main text.

In our case, the initial mass observed is for age 1 for all the species. The mean age at maturity $a_{mat}$ was determined from the age maturity ogive estimated by fitting a logistic regression to age and maturity data. For all species, the scaling exponent of maximum ingestion rate and maintenance rate with body mass $\beta$ was set to 0.75 following the classical power law for metabolic rates (West et al., 1997) and the ratio $\eta$ of gonad energy density to soma energy density was assumed to be 1 by default: the gonado-somatic index (GSI) $r$ was then the energetic GSI. Alternative values of $\beta$ and *q* can be used for other applications whenever necessary data to estimate them are available. To estimate $r$ and the annual mass-specific net energy acquisition rate $\bar{e_{p}}$, the biphasic growth model was fitted on mass-at-age or length-at-age data (thanks to the allometric length-somatic mass relationship), using a nonlinear regression. The same statistical weight was attributed to all age classes by weighting each data point of a given age class by the inverse of its abundance. To estimate the multiplicative factor of maximum mass-specific net energy acquisition rate for the larval stage$\theta'$, the model was fitted between age 0 and 1 using the mass at age 0 ($w_{egg}$) and the mass at age 1 $w_{1}$ as data points (see section 2.4 for $w_{egg}$ estimation). $\theta'$ is an intermediate parameter to estimate $\theta$ the multiplicative factor of the maximum ingestion rate per mass unit for the first year of life (see Eq. 3, main text). $\theta$ is calculated as follow:

$\theta=\frac{I_{max}+\left( \theta^{'}-1 \right)\bar{e_{p}}}{I_{max}}$	(S2)

with $I_{max}$the maximum ingestion rate mass-specific.

The parameterization of the maturation process is based on the probabilistic maturation reaction norm (PMRN), estimated from size, age and maturity data following the method described in Barot et al. (2004) and summarized hereafter. The first step is to estimate an age and size maturity ogive, i.e., the proportion of mature individuals as a function of age and size, with or without interactions, through logistic regression on age, size and maturity data. The second step is to estimate the length increment at each age using size and age data. The third step is to build the PMRN by estimating the probability of becoming mature for each possible age-length combination using the two previous ingredients (see Barot et al. (2004) for further details).

To obtain a linear deterministic maturation reaction norm (LMRN) from the PMRN, a logistic regression is fitted to the estimates of the probabilities of becoming mature as a linear function of age and length. The resulting age-length iso-probability line for which there is a 50% probability of maturing is then taken as the estimate of the LMRN and its intercept and slope are used as estimates of $m_{0}$ and$m_{1}$, respectively. The LMRN is considered deterministic in the sense that when an individual reaches an age-length combination located on the reaction norm, it becomes mature with probability 1 (Eq. 13 in Main Text).

**2 - Estimating the egg mass**

There is not much reliable data on the wet mass of the eggs of marine fish species in the literature. In contrast, relative fecundity, i.e., the number of eggs laid per gram of mature female$\varphi_{S}$ is a much more common data. With $r$, the GSI estimated from the SMALK data (see Supporting Information S2.1), we estimate the mass of an egg needed to obtain a relative fecundity that emerges from the model similar to that observed:

$w_{egg}=\frac{r}{\varphi_{S}}$	(S3)

The number of eggs laid per gram of mature female$\varphi_{S}$ values for the North Sea species are in Supporting Information S4, Table S4 and the reference are in Supporting Information S6 in the column fecundity.

**3 - Physiological thermal response curves parameters estimation**

The thermal response curves in Bioen-OSMOSE are detailed in the eq. 6, 7, 8 and 9 in the main text. The mobilization and maintenance response parameter estimation uses the properties of the resulting net energy available for new tissues $E_{P}$ (Eq. 9 in the main text). $E_{P}$ has the classic bell-shaped response to temperature consistent with the OCLTT theory. Under the assumption of maximum oxygen saturation and combining Eq. 4, 7, and 9 in the main text, mass-specific net energy available for new tissue production $E_{P}(i,t){/w\left( i,t \right)}^{\beta}=e_{P}(i,t)$ can be written

$e_{P}(i,t)=\xi\iota(i,t) \varphi_{M}\left( T(i,t) \right)-c_{m} \varphi_{m}(T(i,t))$.	(S4)

with $\iota(i,t)$ denoting mass-specific ingestion rate $I(i,t)/{w(i,t)}^{\beta}$. Averaging over an individual’s lifetime and under the assumption of a constant inter annual ingestion, we obtain the average mass-specific net energy over an individual’s lifespan $\bar{e_{p}}$ as:

$\bar{e_{p}}=\frac{1}{\tau}\int_{0}^{\tau} e_{P}(i,t)dt=\frac{\xi}{\tau} \bar{\iota}\int_{0}^{\tau} \varphi_{M}\left( T(i,t) \right)dt -\frac{c_{m}}{\tau} \int_{0}^{\tau} \varphi_{m}(T(i,t))dt$.	(S5)

with $\tau$ the individual’s lifespan in years and $\bar{\iota}$ the annual mass-specific ingestion rate.

Under the additional assumption of a constant temperature $T$ throughout an individual’s lifetime, the equation simplifies to

$\bar{e_{p}}(T)=\xi\bar{\iota}\varphi_{M}\left( T \right) -c_{m}\varphi_{m}(T)$	(S6)

with $\bar{\iota}=\frac{1}{\tau}\int_{0}^{\tau} \iota(i,t) dt$ the lifetime-mean annual mass-specific ingestion rate. In these conditions, i.e. maximum oxygen saturation and constant temperature over an individual’s lifetime, our bioenergetic model is strictly equivalent to the biphasic growth model of Boukal et al. (2014).

There are five unknown parameters for the thermal response of mobilization and maintenance, namely $\varepsilon_{M}$, $\varepsilon_{D}$,	and$T_{p}$ for $\varphi_{M}\left( \cdot\right)$ and $\varepsilon_{m}$ and $c_{m}$ for $\varphi_{m}(\cdot)$ (see Table S1). These can be (owing to the various assumptions underlying Eq. S6) estimated together with the mean annual mass-specific ingestion rate $\underline{\iota}$ as a by-product from the numerical resolution of a system of six nonlinear equations resulting from six particular data points. Four data points correspond to remarkable points of the function $\bar{e_{p}}(T)$.The first and second point, are related to the physiological minimum ($T_{min}$), and maximum ($T_{max}$) temperatures that are respectively defined as the lower and upper temperature thresholds at which $\bar{e_{p}}$becomes null, $\bar{e_{p}}(T_{min})=\bar{e_{p}}(T_{max})=0$. The third point relates to the optimum temperature ($T_{opt}$) that corresponds to the temperature at which $\bar{e_{p}}$is maximum and thus its derivative according to temperature is null, ${\frac{\partial\bar{e_{p}}(T)}{\partial T}|}_{T=T_{opt}}=0$. The fourth data point is estimated from observations: at the mean temperature in the distribution area over the year, $T_{a}$, the population of the area acquires energy at the rate per unit of body mass (see 1.1 for estimation procedure for $\bar{e_{p}}$). $T_{min}$, $T_{opt}$, $T_{max}$ and $T_{a}$ are species dependent. The mobilization is a physiological temperature-dependent flux: hypothetically, the value of the flux reaches 0 at a temperature of absolute zero. This fifth data point is species independent. The sixth data point is the value of the maintenance rate $c_{SMR}$ at $T_{ref}$ from the literature. The six data points are used to estimate six parameters with the numerical resolution of a mathematical nonlinear system.

$\left\{ \begin{aligned} \bar{E_{p}}\left( T_{min} \right)=0 \\ \bar{E_{p}}(T_{max})=0 \\ \begin{aligned} \bar{E_{p}}'(T_{opt})=0 \\ \bar{E_{p}}\left( T_{a} \right)=c \\ \varphi_{M}\left( 0 \right)=0 \\ C_{m}=c_{SMR}.e^{\frac{\varepsilon_{m}}{k_{B}}.\left( \frac{1}{T_{ref}} \right)} \end{aligned} \end{aligned} \right.$	(S7)

Table S1: Description of the parameters of the physiological thermal response curves and the data required to estimate them.

|  | Parameter | Description | Source |
| --- | --- | --- | --- |
| Mobilization response | $T_{p}$ | The temperature at which the mobilization reaches its maximum | Estimated |
|  | $\varepsilon_{M}$ | The activation energy for the Arrhenius-like increase with temperature before $T_{p}$ | Estimated |
|  | $\varepsilon_{D}$ | The activation energy for the energy mobilization decline with temperature after $T_{p}$ | Estimated |
|  | $k_{B}$ | The Boltzmann constant | 1.38064852 |
|  | $\underline{\iota}$ | Mean annual mass-specific ingestion rate | Estimated |
| Maintenance response | $C_{m}$ | The absolute mass-specific maintenance rate | Estimated |
|  | $\varepsilon_{m}$ | The activation energy for maintenance rate increase with temperature | Estimated |
|  | $c_{smr}$ | The mass-specific maintenance rate at the temperature $T_{ref}$ | Literature |
|  | $T_{ref}$ | The temperature at which $c_{smr}$ is measured | Literature |
| Net energy response | $T_{min}$ | Temperature at which the net energy reaches 0 for a temperature under $T_{opt}$ | Literature, experimental or from climatic niche |
|  | $T_{opt}$ | Temperature at which $E_{P}$ is maximum |  |
|  | $T_{max}$ | Temperature at which the net energy reaches 0 for a temperature above $T_{opt}$ |  |
|  | $T_{a}$ | Mean temperature in the distribution area over the year | Calculated from forcing files |
|  | $\bar{e_{p}}$ | Mass-specific net energy acquisition rate. Given the $T_{a}$ definition, is the value at $T_{a}$. | Estimated (Supporting Information S2.1) |

A nonlinear equation solver available in R within the package “nleqslv” was used to solve the system (4). To parameterize temperature preference, the use of physiological preference data is advised. In NS-Bioen-OSMOSE, $T_{min}$ and $T_{max}$ are parameterized with Dahlke et al. (2020) database. In parallel, an estimation of $T_{min}$ and $T_{max}$ is done from thermal niches derived from specific global distribution occurrences from databases: the Ocean Biogeographic Information System (<http://www.iobis.org/>), the Global Biodiversity Information Facility (<https://www.gbif.org/>), the Vertebrate biodiversity Networks, <http://vertnet.org/>, and the UC Berkeley's Natural History Data, (<https://bnhm.berkeley.edu/informatics/ecoengine/>). The spatial distribution model method used to obtain the thermal niches is from Ben Rais Lasram et al., (2020). A linear relationship is fitted between the physiological $T_{min}$ and $T_{max}$ and the thermal niches-derived $T_{min}$ and $T_{max}$ respectively, for the species with available data. This linear relationship is used to estimate the physiological $T_{min}$ and $T_{max}$ for the species without data (shrimp, grey gurnard, whiting, mackerel, norway pout and dab) in the database from Dahlke et al. (2020). $T_{opt}$ is estimated from thermal niches estimated with global distribution occurrences and temperature at the occurences, similarly to $T_{min}$ and $T_{max}$.

**4 - Parameter estimation of the mobilization response curve to oxygen**

To estimate the parameters $c_{O,1}$ and $c_{O,2}$ of the mobilization response to oxygen, a nonlinear least square regression (nls function in R) of the response function (Eq. 5 in the main text) was fitted on the respiration rate relative to oxygen concentration or saturation. The data were extracted from the literature (Supporting Information S6). The parameters for species without data were filled by the parameter of the closest species in the phylogenetic tree.

**5 - Parameterization of the reproduction seasonality**

In Bioen-OSMOSE, the spawning seasonality is represented by a set of parameters $sp\left( t \right)$ indicating the proportion of the current energy available in the gonad used to produce and release eggs at the time step $t$.

The proportion of the gonad energy used at a time step $t$, $sp\left( t \right)$, is calculated from the proportion of eggs $p_{e;t}$ released at time step $t$ compared to the annual total released eggs. The seasonality of eggs’ release can be found in the literature and was classically used in previous versions of OSMOSE. The proportion of gonad energy used to release eggs $sp\left( t \right)$at the time step $t$ is linked with $p_{e;t}$ and with the cumulative proportion of the eggs already released within this reproductive season (released at the time steps anterior to the time step $t$ in the reproductive season):

$sp\left( t \right)=\left\{ \begin{aligned} {n_{r}.p}_{e;t} if t=1 \\ \frac{{n_{r}.p}_{e;t}}{\prod_{j=1}^{j=t-1} 1-p_{g;j}} if t>1 \end{aligned} \right.$	(S8)

with $j$ the number of time steps in the reproductive season before the time step$t$ and $n_{r}$ the number of reproductive seasons in the year.

Supporting Information S3: **Mortality process description**

At each time step, a school experiences several mortality sources. The total mortality of a school $i$ is the sum of predation mortality caused by other schools, foraging mortality, starvation mortality, fishing mortality, larval mortality and additional mortalities (i.e. senescence, diseases, and non-explicitly modeled predators). Within each time step $t$, the different mortality sources impact a school $i$ in a random order (across both sources of mortality and schools) and are implemented through iterations over sub-time steps (here 10 sub-time steps occur within a time step) so as to simulate the simultaneous nature of these processes (see <http://documentation.osmose-model.org/> for more details).

The mortality induced by predation emerges from the ingestion process previously described (see section 1.3.6 *Ingestion, assimilation and mobilization*) and thus is an explicit stochastic size-dependent process depending on the spatial co-occurrence between predators and prey. The predation-induced loss of biomass experienced by school $i$ is simply the sum of biomass losses due to the ingestion of all predator schools $j$ present in the same grid cell $c\left( i,t \right)$ at time step $t$ and with adequate minimum $R_{min}$ and maximum $R_{max}$ predator to prey size ratios,

$\frac{dB(i,t)}{dt}= \sum_{j} \frac{\gamma\left( j,i \right)B\left( i,t \right)}{P\left( j,t \right)}I\left( j,t \right)$

with $j\in\left\{ j | \left( c\left( i,t \right)=c\left( j,t \right) \right)\bigcap\left( \frac{L(j,t)}{R_{max}}\leq L(i,t)\leq\frac{L(j,t)}{R_{min}} \right) \right\}$,	(S9)

where $B\left( i,t \right)$ is the biomass of school $i$, $P\left( j,t \right)$ is the prey biomass for school $j$,$R_{min}$ and $R_{max}$ the minimum and maximum predator to prey size ratio based on individual total length, $c\left( i,t \right)$ the cell of the school $i$ and $\gamma\left( j,i \right)$ is the accessibility coefficient of potential prey school $i$to school $j$ (see Eq. 1 and Table 2 in the main text). Then, the ratio $\frac{\gamma\left( j,i \right)B\left( i,t \right)}{P\left( j,t \right)}$ is a proportion of school $i$ in the predator $j$’s ingestion $I\left( j,t \right)$. Change in the number of individuals of school $i$ due to predation during a sub-time step d$t$ is then given by:

$N\left( i,t+dt \right)= N\left( i,t \right)(1-\frac{\sum_{j} \frac{\gamma\left( j,i \right)B\left( i,t \right)}{P\left( j,t \right)} I\left( j,t \right)}{w(i,t)})$

with $j\in\left\{ j | \left( c\left( i,t \right)=c\left( j,t \right) \right)\bigcap\left( \frac{L(j,t)}{R_{max}}\leq L(i,t)\leq\frac{L(j,t)}{R_{min}} \right) \right\}$.	(S10)

Starvation occurs when an individual cannot cover its maintenance needs, i.e., when net energy is negative $E_{P}(i,t)<0$, even by drawing energy from its gonadic reserves, i.e., when $\eta E_{P}(i,t)< -g(i,t)$ (see sub-section “New tissue production: somatic and gonadic growth” for details). In this case, schools undergo a decrease in biomass equaling the energetic deficit that remains after accounting for the energy reserves contained in the gonadic compartment:

$B\left( i,t+\Delta t \right)=B\left( i,t \right)-N\left( i,t \right)\left( \left| E_{P}\left( i,t \right) \right|-g\left( i,t \right)/\eta\right) if \eta E_{P}(i,t)< -g(i,t)$,	(S11)

The energetic debt of time step t+$\Delta t$ is equally shared between the sub time step dt, so that change in the number of individuals of school $i$ during the sub-time step d$t$ due to energetic starvation during time step d$t$ is given by:

$N\left( i,t+dt \right)=\frac{N\left( i,t \right).dt}{\Delta t}(1-\frac{\left| E_{P}\left( i,t \right) \right|-g\left( i,t \right)/\eta}{w\left( i,t \right)}) if \eta E_{P}\left( i,t \right) < -g\left( i,t \right).$	(S12)

As the mortality process is implemented before ingestion (see SI 1), starvation mortality at time step t relies on the net energy amount obtained at the previous time step. During the first time step of a school, i.e., at the egg life stage, the school does not feed to model the yolk reserve. Therefore, starvation does not occur at the first time step nor at the second time step as starvation relies on the energy acquired at the previous time step.

Schools experience size-selective fishing that generates an instantaneous fishing mortality rate $F(i,t)$ for a school $i$ at time step $t$ defined as follows:

$F\left( i,t \right)=F_{max} q\left( L(i,t) \right)$	(S13)

with $q(L(i,t))$ the size-selectivity of the fishery that varies between 0 and 1 with individual length, and Fmax is the maximum instantaneous fishing mortality rate. Change in the number of individuals in school $i$ due to fishing during time sub time step $dt$ is then obtained as follows:

$N\left( i,t+dt \right)=N\left( i,t \right) e^{-F\left( i,t \right) dt}$	(S14)

Finally, in order to account for the additional mortality sources that are not represented by the previous mortality terms (e.g. senescence, diseases, or non-explicitly modeled predators), an instantaneous life-stage specific additional mortality rate $M(i,t)$ is defined as follows:

$M(i,t)=\left\{ \begin{aligned} \mu_{l^{'}}if a\left( i,t \right)<a_{l^{'}} \\ \mu otherwise \end{aligned} \right.$	(S15)

with $\mu_{l^{'}}$ the instantaneous larval additional mortality rate and $\mu$ its counterpart after $a_{l^{'}}$, the first feeding age, and for the rest of the life cycle. Change in the number of individuals in school $i$ due to additional mortality during sub-time step $dt$ is given by:

$N\left( i,t+dt \right)=N\left( i,t \right) e^{-M\left( i,t \right) dt}$	(S16)

Table S2: Description of the parameters of the mortality sub-model

| **Symbol** | **Description** | **Units** | **Equations** | **Source** |
| --- | --- | --- | --- | --- |
| $\Delta t$ | Time step | $y$ |  | - |
| $dt$ | Sub-time step |  |  |  |
| $F_{max}$ | Maximum instantaneous fishing mortality rate | y^-1^ | S13 | Calibrated |
| $a_{l^{'}}$ | First feeding age | $y$ | S15 | Literature |
| $\mu_{l^{'}}$ | Instantaneous larval additional mortality rate | y^-1^ | S15 | Calibrated |
| $\mu$ | Instantaneous additional mortality rate for individuals older than first-feeding agee | y^-1^ | S15 | Calibrated |

Supporting Information S4: **Input parameters and intermediate parameters of Bioen-OSMOSE-NS for the 16 species modeled explicitly**.

**1 – Species specific input parameters**

Table S3: Input parameters parameters of Bioen-OSMOSE-NS for the 16 species modeled explicitly. $\alpha, k, r, m_{0},$ $m_{1}$, $w_{egg}$ and $\theta$ are estimated (see Supporting Information S2.1). $a_{max}$ drawn from the literature (see Supporting Information S6). When $R_{min}$ and $R_{max}$ shift with ontogeny, the size threshold of the shift is provided in the predation threshold column. $R_{min}$, $R_{max}$ and their threshold are from literature. Mobilization and maintenance parameters are estimated (see Supporting Information S2.3)._._ $I_{max}$ and mortality parameters are calibrated (see 2.2.3 in main text for details about the calibration process). Some parameters are assumed equal for all the species: $\eta$ is set to 1, $\xi$ is set to 0.8, $a_{l}$ is set to 1, and $\beta$ is set to 0.75. The references of parameters from literature are in Supporting information S6.

|  | **GROWTH, REPRODUCTION AND CONDITION** | | | | | | | | | | **PREDATION** | | |
| --- | --- | --- | --- | --- | --- | --- | --- | --- | --- | --- | --- | --- | --- |
|  | $\alpha$ | $k$ | $r$ | $I_{max}$ | $m_{1}$ | $m_{0}$ | $w_{egg}$ | $a_{max}$ | $\theta$ | $n_{s(i)}$ | $R_{min}$ | $R_{max}$ | Threshold |
| Species | *-* | *g.cm^-𝛼^* | *-* | *g.g^-β^*${\Delta t}^{-1}$ | *cm.y^-1^* | *cm* | *g* | *year* | *-* | *-* | *-* | *-* | *cm* |
| Herring (*Clupea harengus)* | 3.41 | 0.002 | 0.77 | 14 | -0.9 | 25.36 | 6.67e-04 | 17 | 1.56 | 25 | 250 | 5 | - |
| Mackerel (*Scomber scombrus)* | 3.06 | 0.007 | 0.75 | 16.4 | 0.78 | 25.76 | 5.86e-04 | 17 | 1.66 | 38 | 150 | 2.5 | - |
| Sandeel (*Ammodytes spp)* | 3.44 | 0.001 | 1.2 | 9.5 | 0 | 15 | 9.97e-04 | 10 | 1.66 | 38 | 1000 | 8 | - |
| Sprat (*Sprattus sprattus)* | 3.15 | 0.005 | 0.56 | 12.1 | 0 | 12.96 | 2.80e-04 | 10 | 1.56 | 38 | 1000 | 8 | - |
| Norway pout (*Trisopterus esmarkii)* | 3.1 | 0.006 | 0.5 | 9.9 | -3.54 | 19.22 | 1.57e-04 | 7 | 1.92 | 30 | 500 | 1.5 | - |
| Plaice (*Pleuronectes platessa)* | 3.02 | 0.009 | 0.52 | 10.9 | -4.99 | 43.51 | 2.04e-03 | 25 | 1.85 | 50 | 100/50 | 5 | 8 |
| Sole (*Solea solea)* | 3.26 | 0.004 | 0.6 | 9.9 | 0.69 | 19.17 | 5.81e-04 | 17 | 1.6 | 50 | 100/50 | 10 | 8 |
| Saithe (*Pollachius virens)* | 3.24 | 0.003 | 0.21 | 14.5 | -1.75 | 74.1 | 3.30e-04 | 21 | 1.82 | 30 | 50/15 | 2 | 20 |
| Cod (*Gadus morhua)* | 3.21 | 0.005 | 0.54 | 21.6 | -2.74 | 75.26 | 4.55e-04 | 17 | 1.38 | 25 | 50/15 | 2/1.3 | 12 |
| Haddock (M*elanogrammus aeglefinus)* | 3.24 | 0.004 | 0.59 | 15.8 | -1.62 | 32.85 | 1.36e-03 | 18 | 1.56 | 38 | 50/15 | 1.7 | 12 |
| Horse Mackerel (*Trachurus trachurus)* | 2.7 | 0.027 | 0.55 | 13.4 | 0 | 18.5 | 7.35e-04 | 15 | 1.51 | 38 | 150 | 2.5 | - |
| Whiting (*Merlangius merlangus)* | 3.26 | 0.003 | 0.73 | 17.8 | -0.33 | 21.72 | 4.29e-04 | 16 | 1.39 | 38 | 100/20/10 | 1.5/1.1 | 10/40 |
| Dab (*Limanda limanda)* | 3.21 | 0.006 | 0.5 | 8.8 | -2.47 | 19.03 | 1.72e-04 | 10 | 1.68 | 50 | 100/50 | 5 | 30 |
| Grey gurnard (*Eutrigla gurnardus)* | 3.15 | 0.005 | 0.87 | 13.6 | -6.81 | 28.89 | 9.92e-04 | 15 | 1.48 | 30 | 100/30 | 2 | 12 |
| Hake (*Merluccius merluccius)* | 3.16 | 0.004 | 0.29 | 17.3 | 1.47 | 34.15 | 6.72e-04 | 15 | 1.84 | 20 | 50/15 | 2/1.1 | 12 |
| Shrimp | 2.81 | 0.01 | 1.65 | 8.9 | 0 | 5.37 | 2.03e-03 | 3 | 1.07 | 25 | 1000 | 8 | - |

| **MAINTENANCE** | | **MOBILIZATION** | | | | | **MORTALITY** | | |
| --- | --- | --- | --- | --- | --- | --- | --- | --- | --- |
| **TEMPERATURE** | | **TEMPERATURE** | | | **OXYGEN** | | **FISHING** | **LARVAL** | **ADDITIONAL** |
| $\varepsilon_{m}$ | $c_{m}$ | $\varepsilon_{M}$ | $\varepsilon_{D}-\varepsilon_{M}$ | $T_{p}$ | $c_{O,1}$ | $c_{O,2}$ | $F_{max}$ | $\mu_{l'}$ | $\mu$ |
| $eV$ | $g\cdot{\Delta t}^{-1}$ | $eV$ | $eV$ | *°C* | *-* | *%* | *y^-1^* | *y^-1^* | *y^-1^* |
| 0.6 | 1.49E+11 | 1.93 | 1.01E-04 | 43.1 | 1 | 8.1 | 0.8 | 6.1 | 0.17 |
| 0.33 | 5.03E+06 | 1.67 | 1.02E-04 | 46.85 | 1 | 8.1 | 2.86 | 4.65 | 0.3 |
| 0.47 | 5.40E+08 | 1.96 | 1.01E-04 | 43.29 | 1 | 7.3 | 1.79 | 2.26 | 0.19 |
| 0.27 | 2.57E+05 | 1.15 | 1.06E-04 | 68.31 | 1 | 8.1 | 0.33 | 0.94 | 0.12 |
| 0.17 | 5.82E+03 | 3.21 | 9.43E-05 | 23.8 | 1.48 | 54.6 | 0.58 | 3.76 | 0.31 |
| 0.38 | 1.54E+07 | 1.8 | 1.02E-04 | 46.59 | 2 | 99 | 0.13 | 5.22 | 0.07 |
| 0.42 | 5.28E+07 | 2.54 | 9.87E-05 | 35.12 | 2.6 | 179.4 | 0.26 | 7.01 | 0.34 |
| 0.26 | 2.74E+05 | 2.64 | 9.59E-05 | 27.87 | 1.48 | 54.6 | 0.53 | 2.73 | 0.6 |
| 0.48 | 2.07E+09 | 2.27 | 9.84E-05 | 34.77 | 1.48 | 54.6 | 0.31 | 9.34 | 0.58 |
| 0.31 | 1.96E+06 | 2.43 | 9.69E-05 | 30.67 | 1.48 | 54.6 | 0.08 | 2.82 | 0.8 |
| 0.25 | 1.04E+05 | 2.28 | 9.89E-05 | 39.39 | 1 | 8.1 | 0.1 | 1.2 | 0.27 |
| 0.3 | 1.36E+06 | 1.93 | 1.01E-04 | 41.91 | 1.48 | 54.6 | 0.61 | 8.56 | 0.16 |
| 0.46 | 2.86E+08 | 2.55 | 9.86E-05 | 34.92 | 2.3095 | 139.189 | 0.18 | 4.63 | 0.25 |
| 0.31 | 1.47E+06 | 1.98 | 1.00E-04 | 41.49 | 1.48 | 54.6 | 0.36 | 5.15 | 0.07 |
| 0.25 | 1.74E+05 | 2.92 | 9.60E-05 | 30.06 | 1.48 | 54.6 | 0.42 | 7.72 | 0.32 |
| 0.4 | 3.10E+07 | 2.25 | 9.97E-05 | 38.54 | 1.475 | 47 | 0.02 | 0.06 | 0.21 |

**2 – Species specific intermediate parameters**

Table S4: Species specific intermediate parameters. Intermediate parameters are parameter that are not input parameters of Bioen-OSMOSE-NS but that are used to estimated directly (see Supporting Information S2) or indirectly through calibration process (see Section 2.2.3) input parameters of Bioen-OSMOSE-NS.

|  | **Fecundity** | **Larval growth** | **Maintenance** | | **Temperature preference** | | |
| --- | --- | --- | --- | --- | --- | --- | --- |
|  | $\varphi_{s}$ | $\theta'$ | $T_{ref}$ | $c_{smr}$ | $T_{min}$ | $T_{opt}$ | $T_{max}$ |
| Species | *g.(g female)^- 1^.y^-1^* | **-** | **°C** | *g.g^-^*^α^.*y^-1^* | **°C** | **°C** | **°C** |
| Herring | 1159 | 3.36 | 9.6 | 5.51 | -1.3 | 10.3 | 20.1 |
| Mackerel | 1275 | 3.93 | 1 | 5.20 | -0.5 | 10.7 | 24.3 |
| Sandeel | 1202 | 3.2 | 10 | 2.84 | -0.7 | 12.5 | 27.1 |
| Sprat | 1993 | 5.52 | 9.6 | 5.51 | -0.1 | 12.1 | 24.8 |
| Norway pout | 3199 | 5.66 | 10 | 2.70 | 1.4 | 7.7 | 25.1 |
| Plaice | 255 | 4.46 | 14 | 3.95 | 1 | 12.6 | 26.5 |
| Sole | 1040 | 2.85 | 10 | 1.07 | 1.6 | 12.9 | 31.3 |
| Saithe | 630 | 4.47 | 10 | 2.79 | -0.2 | 7.2 | 21.8 |
| Cod | 1188 | 2.1 | 10 | 2.22 | -2 | 9.1 | 23 |
| Haddock | 430 | 2.91 | 10 | 2.70 | -0.9 | 7.9 | 23.3 |
| Horse Mackerel | 1655 | 3.36 | 14 | 6.25 | 4.8 | 14.5 | 33.5 |
| Whiting | 1700 | 2.48 | 10 | 3.09 | 1.1 | 11.6 | 27 |
| Dab | 2931 | 3.2 | 14 | 3.95 | 1.8 | 12.1 | 27 |
| Grey gurnard | 873 | 2.79 | 10 | 2.70 | 1.5 | 12 | 27.7 |
| Hake | 426 | 4.11 | 10 | 2.70 | 3.7 | 11.3 | 28.9 |
| Shrimp | 815 | 1.28 | 8 | 1.85 | 1.5 | 12.5 | 28.1 |

**3 – Predator-prey accessibility matrix.**

Table S5: Accessibility coefficients $\gamma$ (Eq. 1) between potential predators and prey depend on their water-column position and represent the proportion of prey biomass accessible to a predator. The vertical position of individuals depends on their life stage, which is indicated in the last column. Shrimp (>0.25 year) and sandeels are exceptions due to their day/night migrations.

|  |  | Predator | | | | |  |
| --- | --- | --- | --- | --- | --- | --- | --- |
|  |  | Pelagic | Demersal | Benthic | Sandeel (>0.25 year) | Shrimp |  |
| Focus Prey | Pelagic | 0.8 | 0.4 | 0 | 0.8 | 0 | Norway pout (<0.25 year), Sandeel (<0.25 year), Cod (<0.25 year), Saithe (<0.25 year), Grey gurnard (<0.25 year), Whiting (<0.25 year), Haddock (<0.25 year), Dab (<0.25 year), Sole (<0.25 year), Plaice (<0.25 year), Hake (<0.25 year), Sprat, Horse mackerel, Mackerel, Herring |
|  | Demersal | 0.4 | 0.8 | 0 | 0.4 | 0 | Norway pout (>0.25 year), Cod (>0.25 year), Saithe (>0.25 year), Grey gurnard (>0.25 year),Whiting (>0.25 year), Haddock (>0.25 year), Hake (>0.25 year) |
|  | Benthic | 0 | 0.4 | 0.8 | 0 | 0.8 | Dab (>0.25 year), Sole (>0.25 year), Plaice (>0.25 year) |
|  | Sandeels (>0.25 year) | 0.4 | 0.8 | 0.4 | 0.8 | 0.4 | Sandeels (>0.25 year) |
|  | Shrimp | 0.2 | 0.4 | 0.8 | 0.2 | 0.8 | Shrimp |
| Ressource prey | Pelagic LTL | 1 | 0.5 | 0 | 1 | 0.5 | Dinoflagellates, Diatoms, Micro zooplankton, Meso zooplankton, Macro zooplankton |
|  | Benthic LTL | 0.04 | 0.5 | 1 | 0.04 | 0.5 | Suspension Feeders, Deposit Feeders, Meiobenthos |

Supporting Information S5: **Parameters of the low trophic level (LTL) groups.** The trophic levels are set based on expert knowledge, except for the meso-zooplankton, the suspension feeder, the large and very large benthos for which trophic levels are taken from Kopp et al. (2015). The accessibility coefficients to fish are estimated through the calibration process (see 2.2.3 Calibration).

|  | LTL groups | Size range (cm) | Trophic level | Accessibility coefficient to fish |
| --- | --- | --- | --- | --- |
| PELAGIC PREY | Micro-phytoplankton | 0.0002 – 0.002 | 1 | 0.133 |
|  | Diatoms | 0.001 – 0.02 | 1 | 0.028 |
|  | Hetero-trophic flagellates | 0.0002 – 0.002 | 2 | 0.187 |
|  | Micro-zooplankton | 0.002 – 0.02 | 2 | 0.032 |
|  | Meso-zooplankton | 0.02 – 0.5 | 2.3 | 0.073 |
| BENTHIC PREY | Suspension feeders | 2 - 5 | 2.2 (bivalves) | 0.002 |
|  | Deposit feeders | 0.1 - 2 | 2 | 5.7e-5 |
|  | Meio benthos | 0.0045 - 0.5 | 3 | 0.001 |
|  | Large benthos | 5-10 | 2.6 (Small Crab) | 0.018 |
|  | Very large benthos | 10-15 | 3.6 (Large Crab) | 0.014 |

Supporting Information S6**: References for the life cycle parameters used in the North Sea model.** The length-mass parameters were estimated from SMALK data, except for shrimp (Oh et al., 2001). $R_{min}$ and $R_{max}$ based on Pinnegar (2014) and manually adjusted on diet.

| Species | Fecundity | Spawning season | Maintenance | Oxygen response |
| --- | --- | --- | --- | --- |
| Herring | Muus & Nielsen, 1999 | Muus & Nielsen, 1999, Baxter, 1958 | Same as sprat | Same as mackerel |
| Mackerel | Macer, 1976 | Muus & Nielsen, 1999 | Dickson, 2002 | (Dickson, 2002) |
| Sandeel | Bergstad et al., 2001 | Muus & Nielsen, 1999 | Behrens & Steffensen, 2007 | Behrens & Steffensen, 2007 |
| Sprat | De Silva, 1973 | Koli, 1990; MacKenzie & Köster, 2004 | Meskendahl, 2013 | Same as mackerel |
| Norway pout | Christiansen et al., 1997 | Muus & Nielsen, 1999 | Teleost value, Clarke & Johnston, 1999 | Same as cod |
| Plaice | van Damme et al., 2009 | Fox et al., 2000 | Clarke & Johnston, 1999 | Steffensen et al., 1982 |
| Sole | Witthames et al., 1995 | Quéro et al., 1986 | Lefrançois & Claireaux, 2003 | Lefrançois & Claireaux, 2003 |
| Saithe | Cohen et al., 1990 | ICES, 2012 | Steinhausen et al., 2005 | Same as cod |
| Cod | Yoneda & Wright, 2004 | Narberhaus et al., 2012; Cohen et al., 1990 | Claireaux et al., 2000 | Claireaux et al., 2000 |
| Haddock | Sonina, 1971 | Muus & Nielsen, 1999 | Average of cod, saithe and whiting rate | Same as cod |
| Horse Mackerel | Eltink & Vingerhoed, 1989 | Muus & Nielsen, 1999 | Geist et al., 2013 | Same as mackerel |
| Whiting | Hislop, 1966 | Hehir, 2003;  Cohen et al., 1990 | Steinhausen et al., 2005 | Same as cod |
| Dab | Bohl, 1956 | Muus & Nielsen, 1999 | Same as sole | Average of sole and plaice response |
| Grey gurnard | Muus & Nielsen, 1999 | Muus & Nielsen, 1999 | Teleost value, Clarke & Johnston, 1999 | Same as cod |
| Hake | Mehault et al., 2010 | Mehault et al., 2010 | Average of cod, saithe and whiting rate | Same as cod |
| Shrimp | Bilgin & Samsun, 2006 | WGCRAN, 2015 | Dupont-Prinet et al., 2013 | Dupont-Prinet et al., 2013 |

Supporting Information S7: **Species distribution relative to ontogenetic stages.**

The threshold between larvae and juvenile stage is set to 0.25 year for all the species. The threshold between juvenile and adult stage is species dependent and corresponds to the mean age at maturation (a_mat_). The distribution was obtained from IBTS data for period 2010-2019, Coull et al. (1998), Sundbly et al. (2017), Van der Land (1991) and Mackinson & Daskalov (2007).

| Stage\  Species | Egg and larvae (<0.25y) | Juvenile (0.25y< - <a_amat)_ | Adult (>a_mat_) |
| --- | --- | --- | --- |
| Herring  (a_mat_ = 2.9y) | 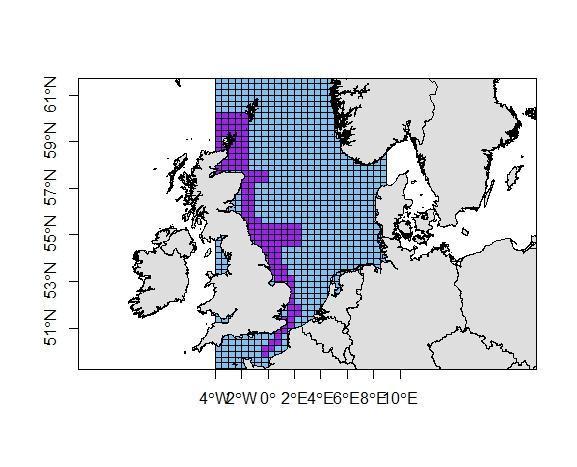 | 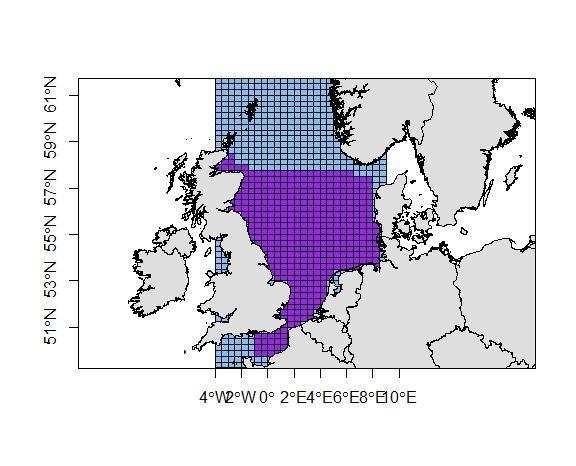 | 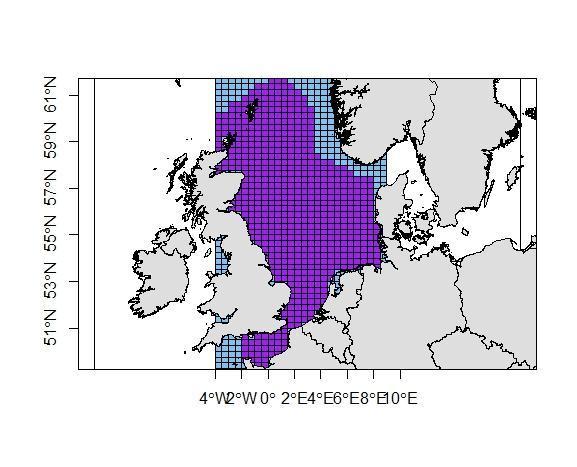 |
| Mackerel  (a_mat_ = 2.5y) | 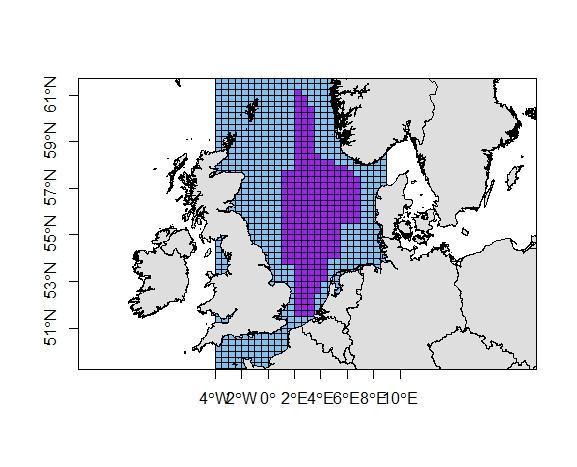 | 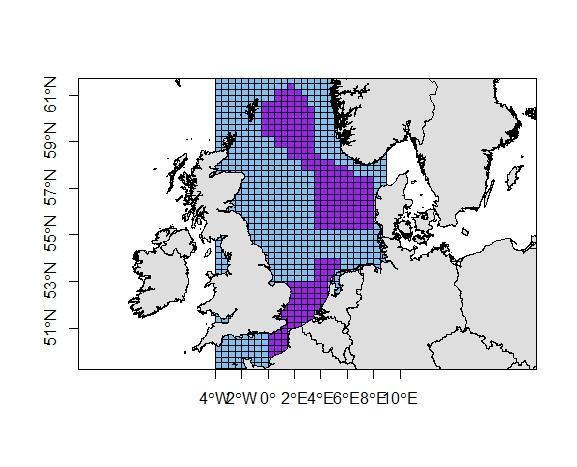 | 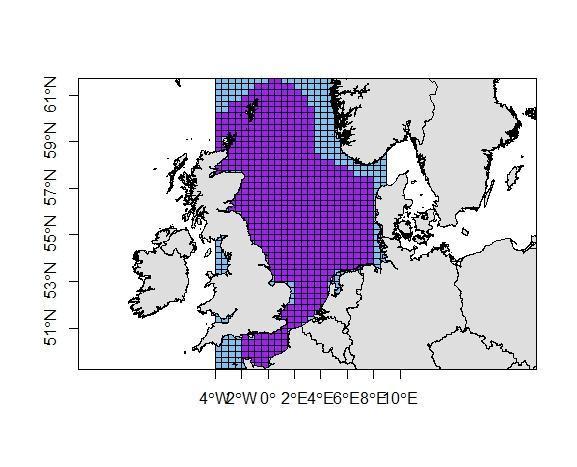 |
| Sandeel  (a_mat_ = 2.2y) | 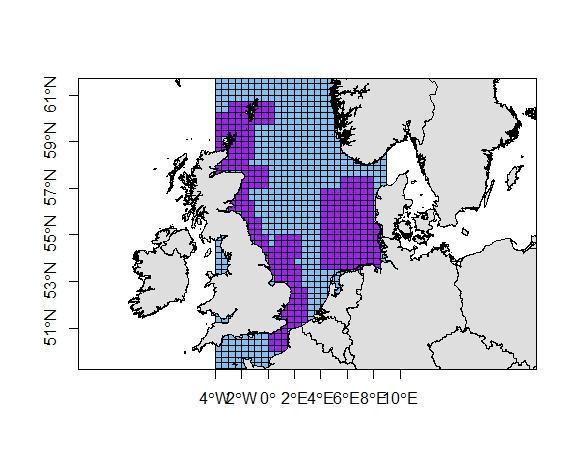 | 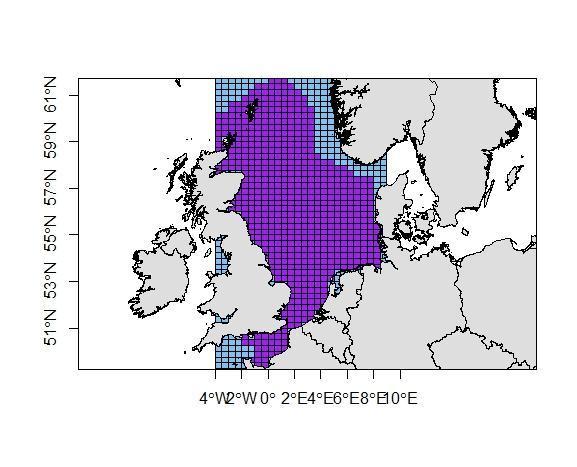 | |
| Sprat  (a_mat_ = 2.3y) | 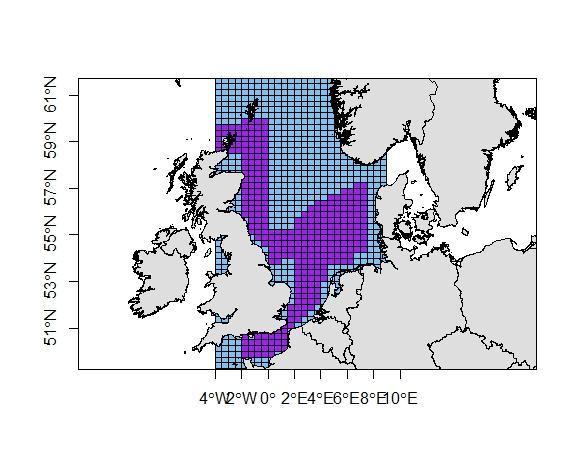 | 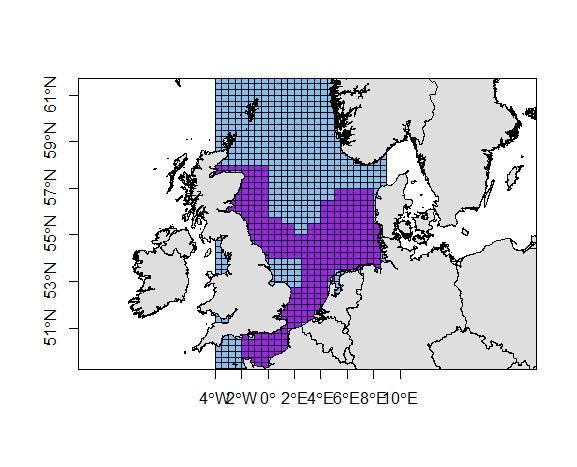 | 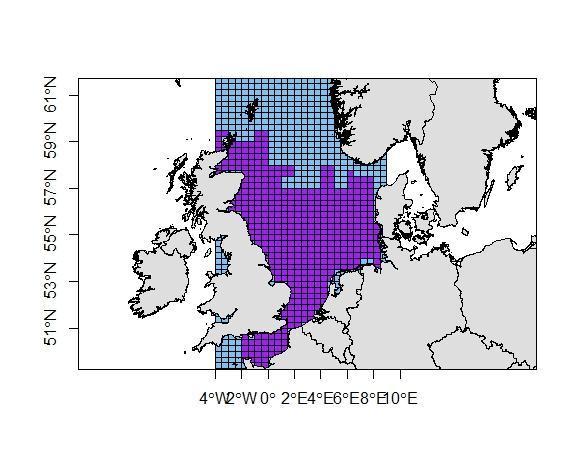 |
| Norway pout  (a_mat_ = 2.0y) | 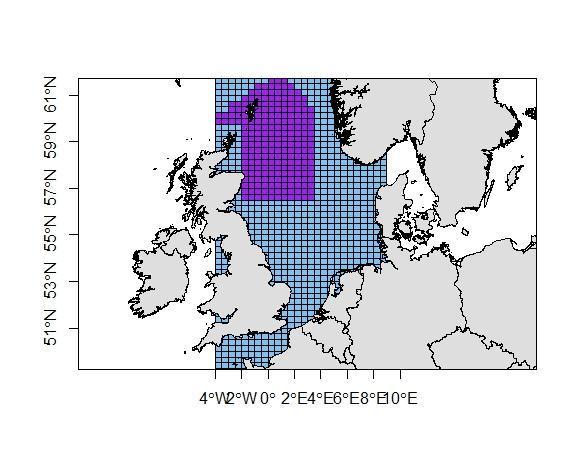 | 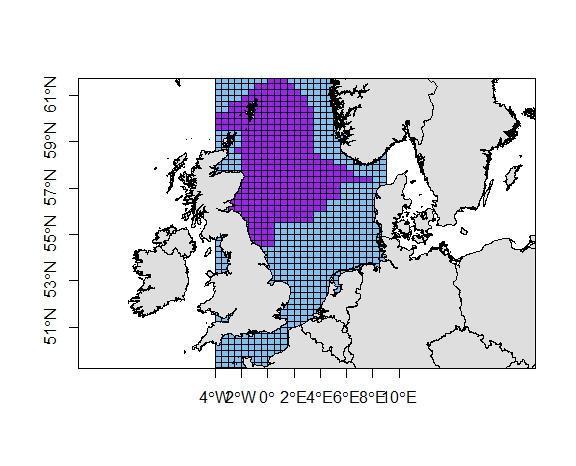 | 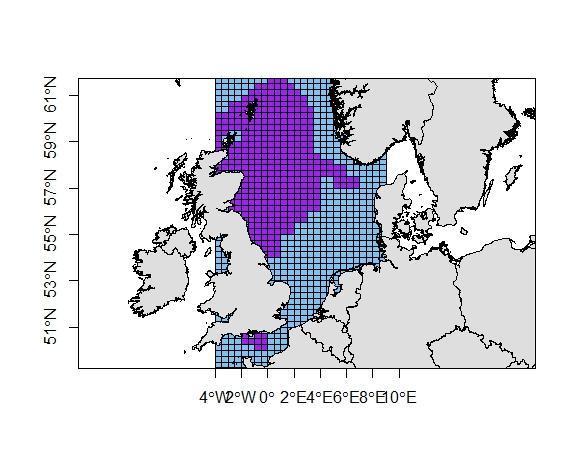 |
| Plaice  (a_mat_ = 3.8y) | 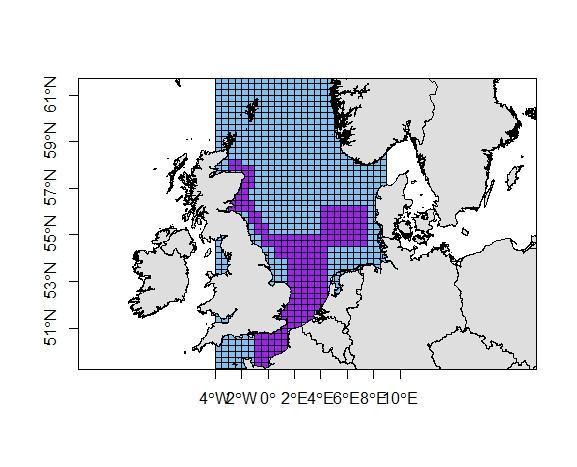 | 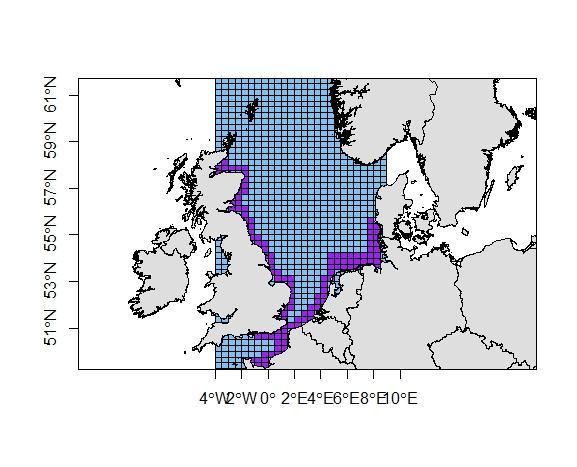 | 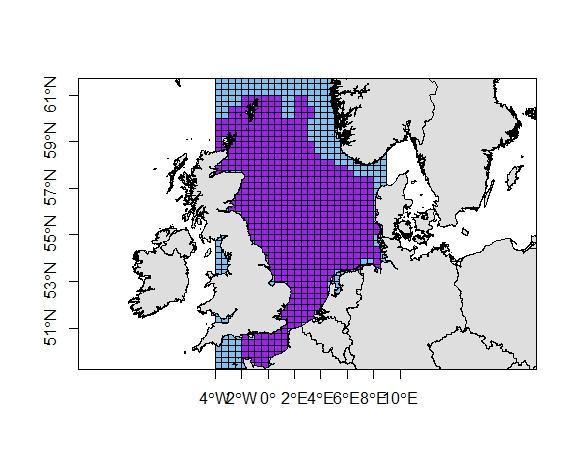 |
| Sole  (a_mat_ = 3.0y) | 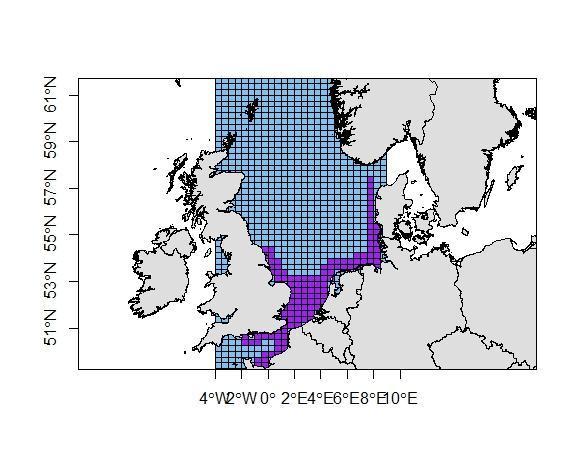 | 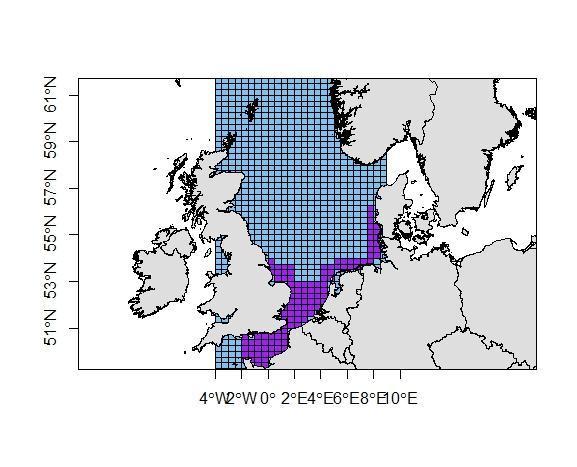 | 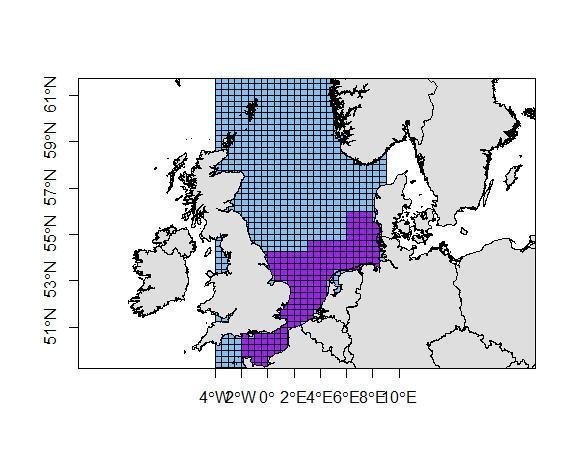 |
| Saithe  (a_mat_ = 5.8y) | 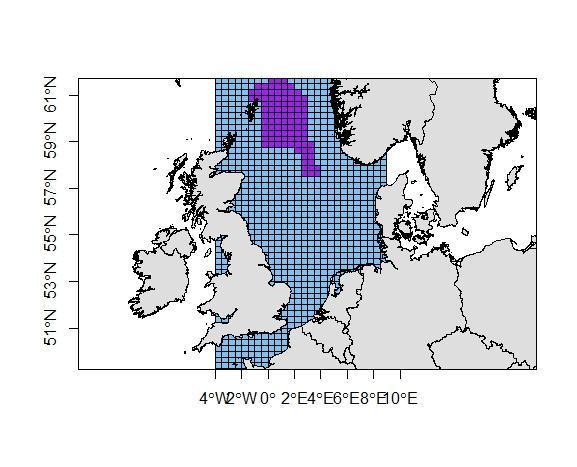 | 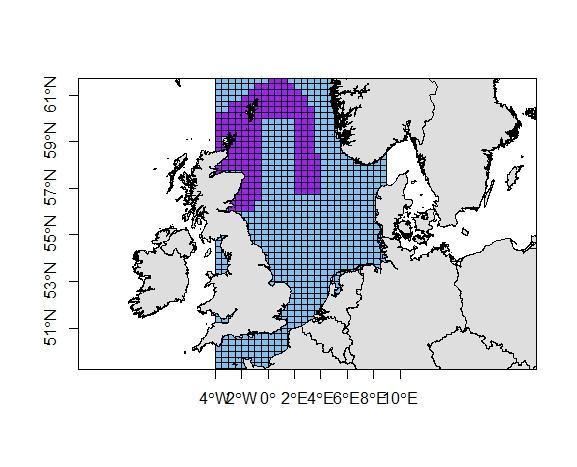 | 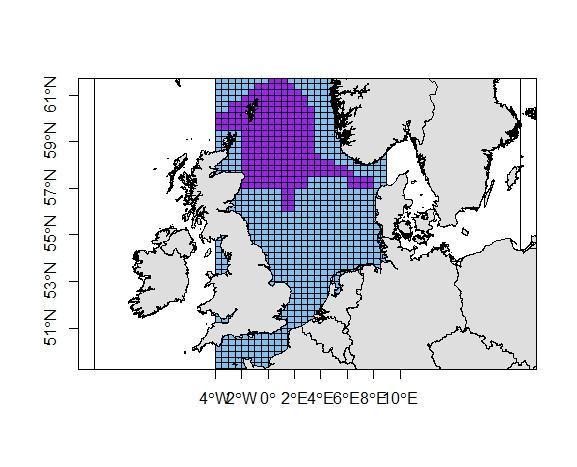 |
| Cod  (a_mat_ = 3.9y) | 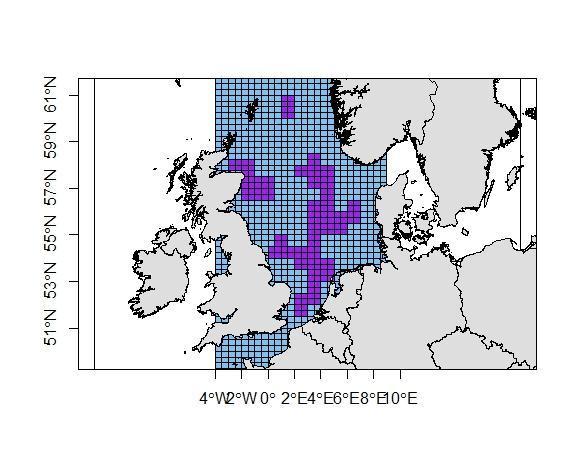 | 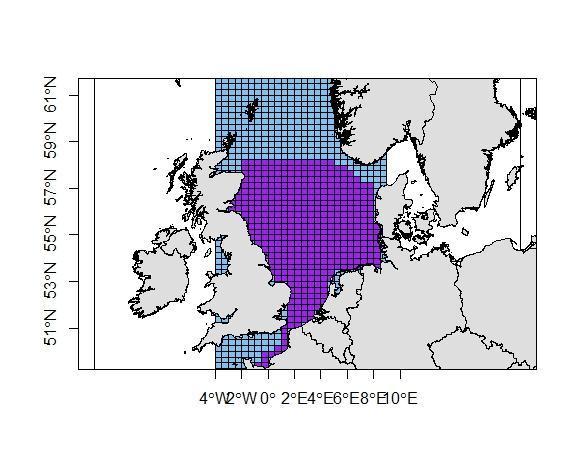 | 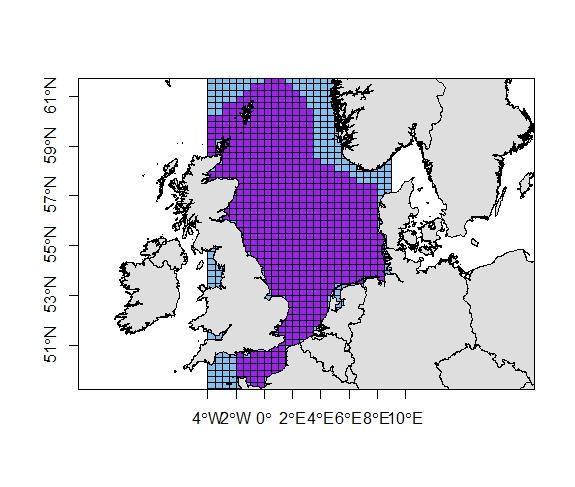 |
| Haddock  (a_mat_ = 2.6y) | 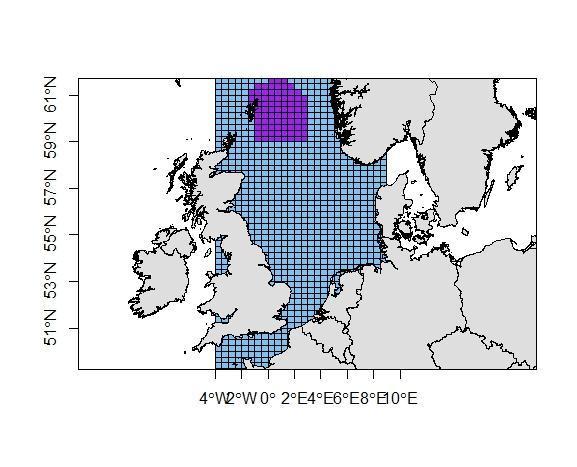 | 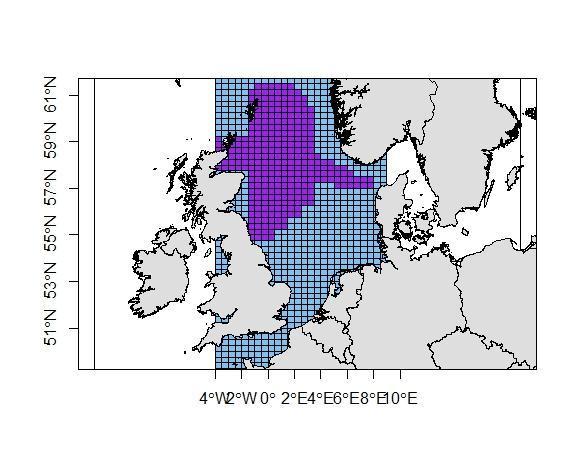 | 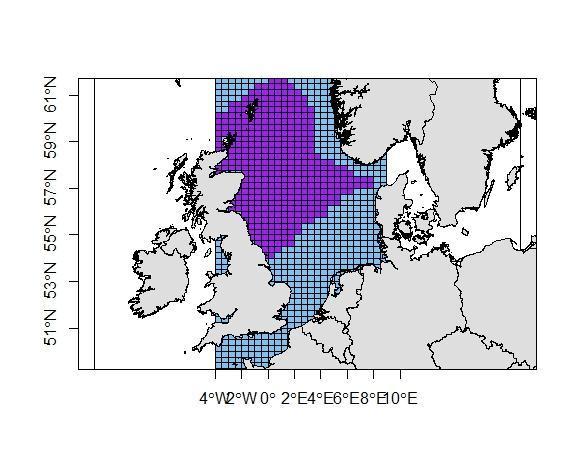 |
| Horse Mackerel  (a_mat_ = 1.8y) | 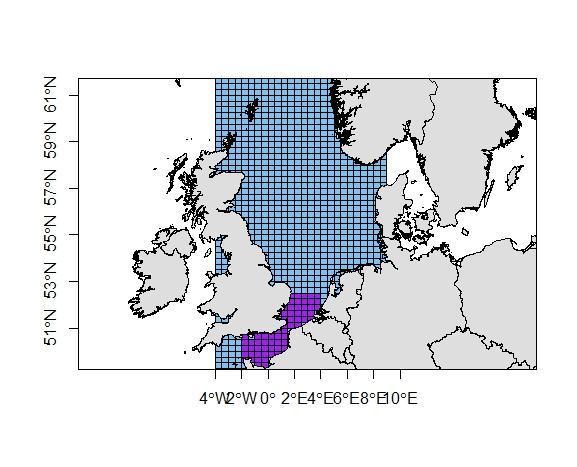 | 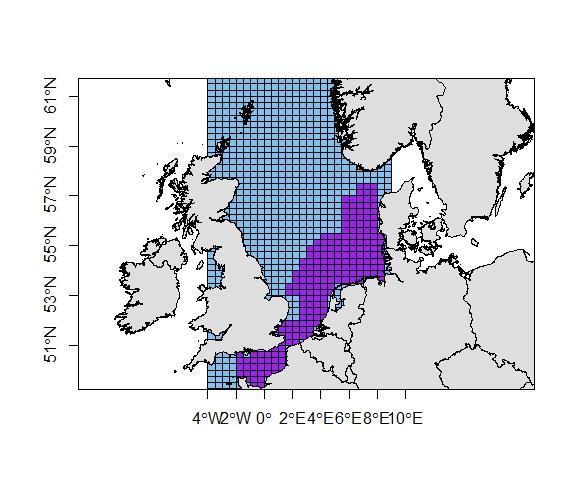 | 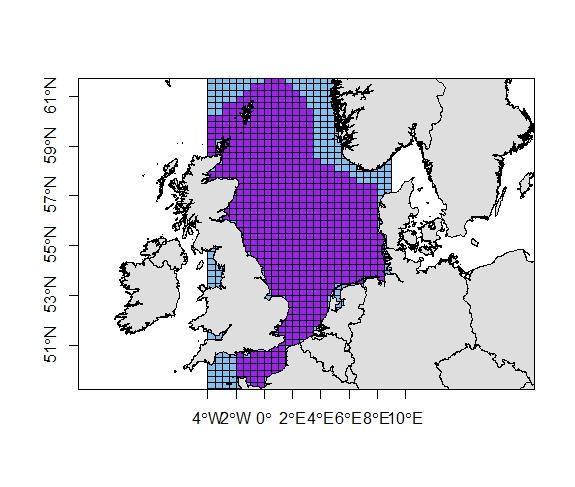 |
| Whiting  (a_mat_ = 2.2y) | 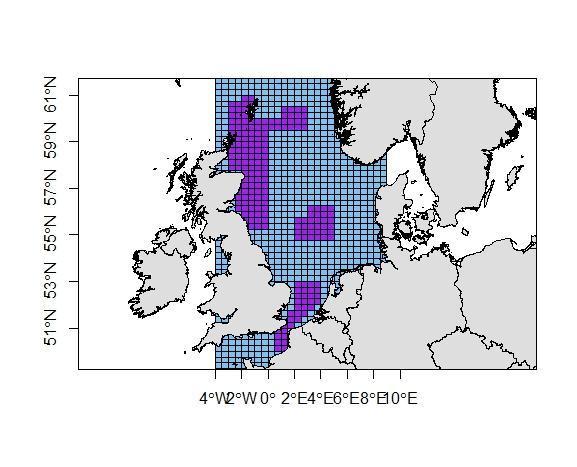 | | 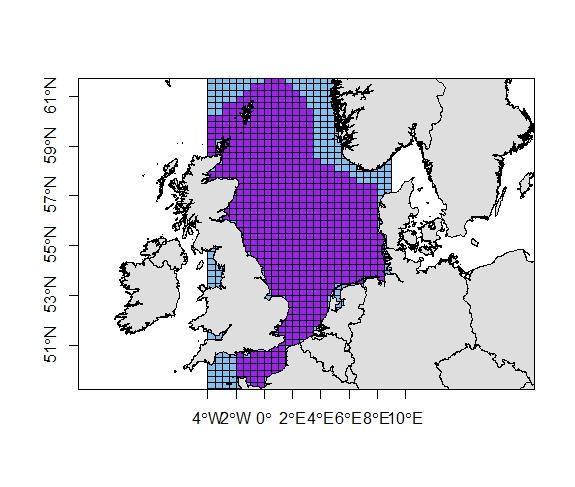 |
| Dab  (a_mat_ = 1.7y) | 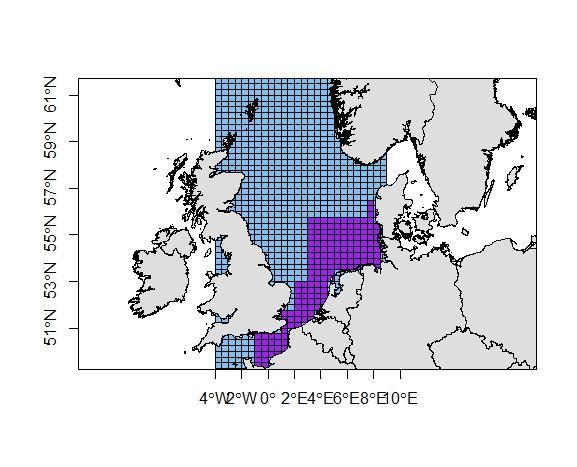 | | 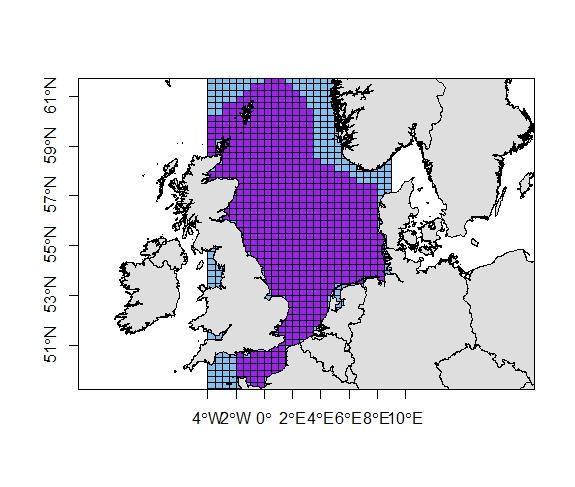 |
| Grey gurnard  (a_mat_ = 2.2y) | 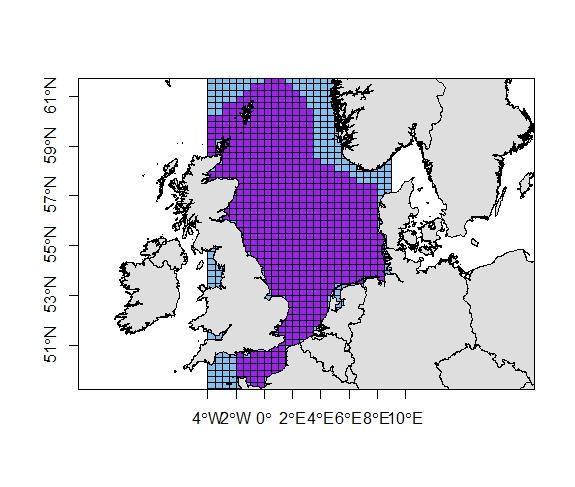 | | |
| Hake  (a_mat_ = 3.4y) | 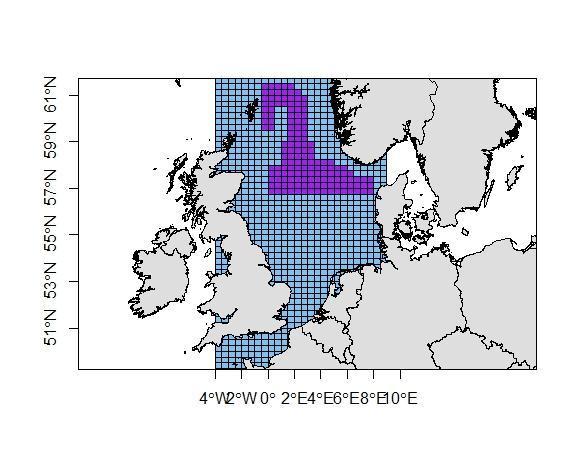 | 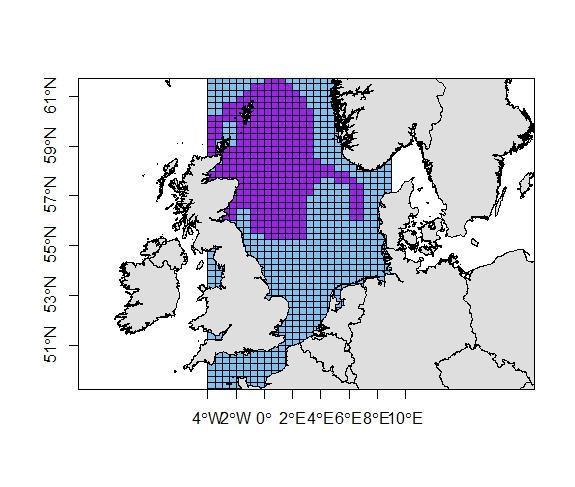 | 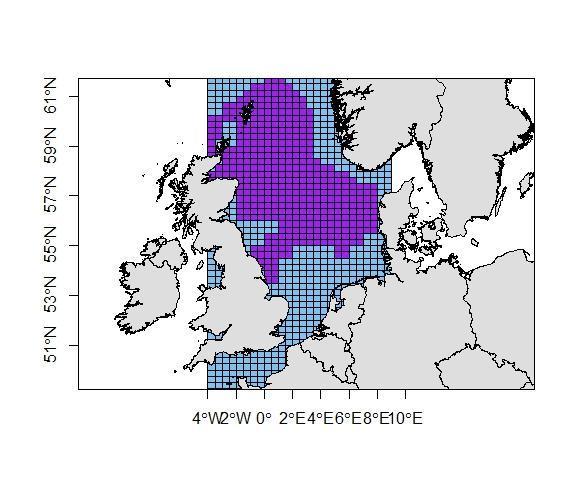 |
| Shrimp  (a_mat_ = 1y) | 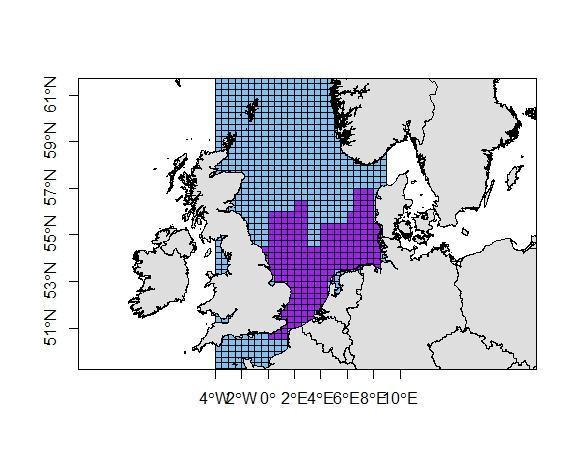 | | |

Supporting Information S8: **Reproductive season.** The diagram is the proportion of released eggs per time step $t p_{e;t}$ relative to the annual total released eggs (see Eq. S8). The references are given in Supporting information S6.

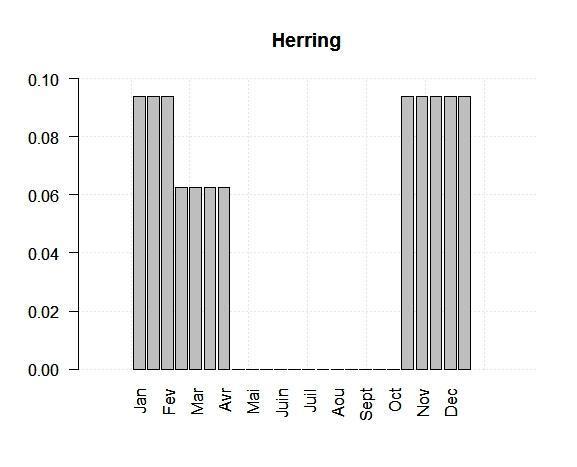

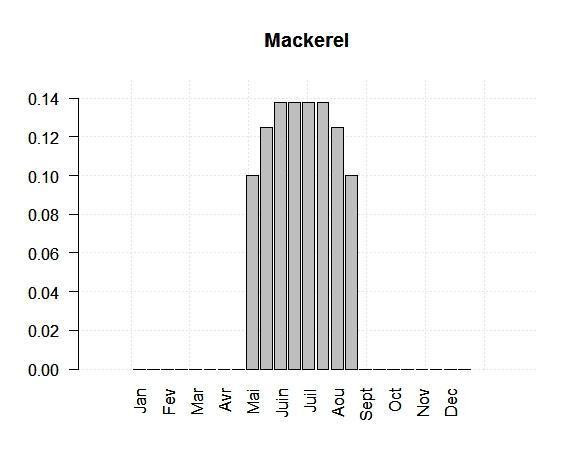

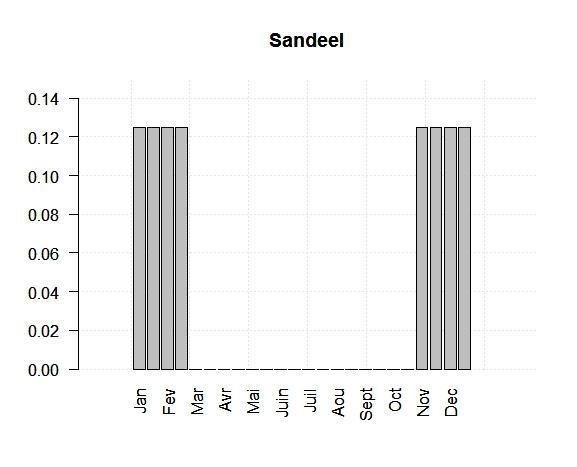

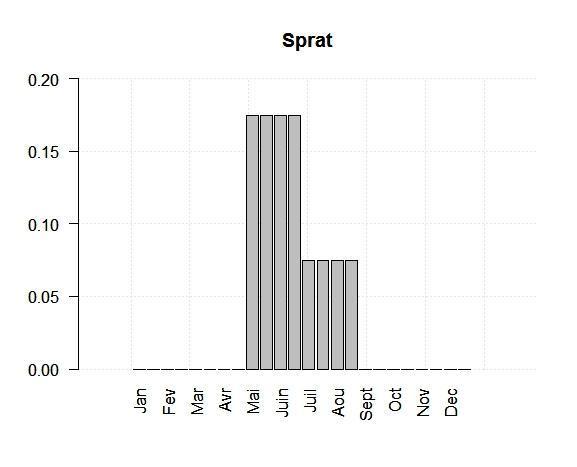

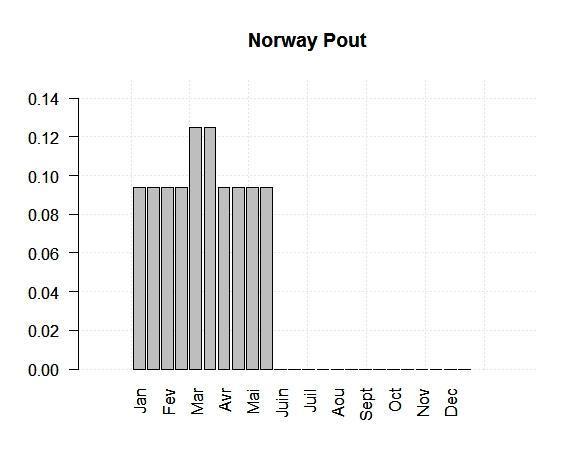

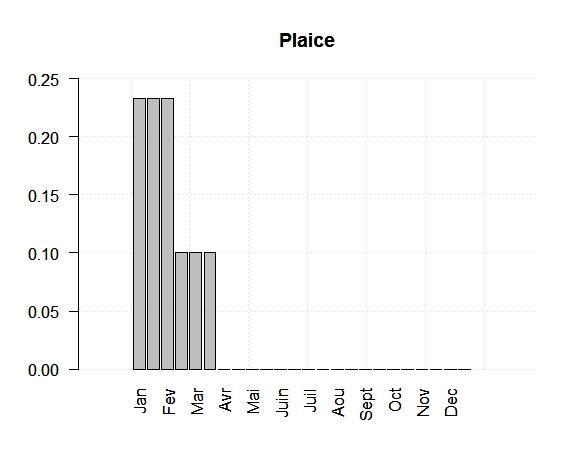

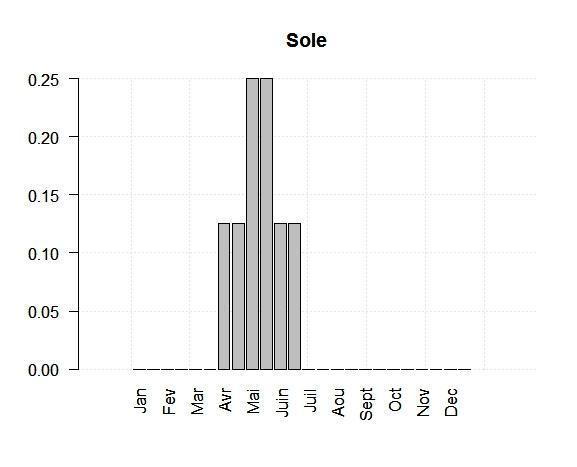

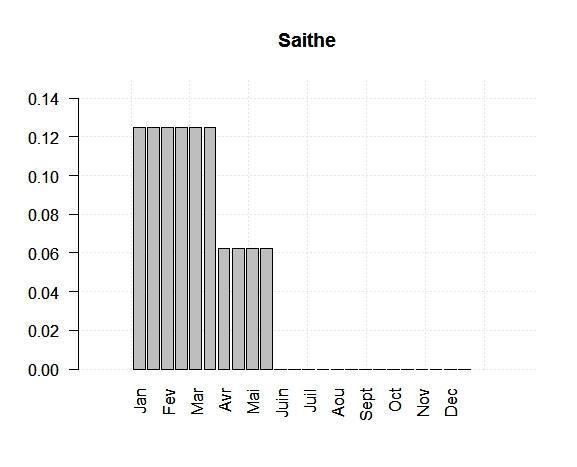

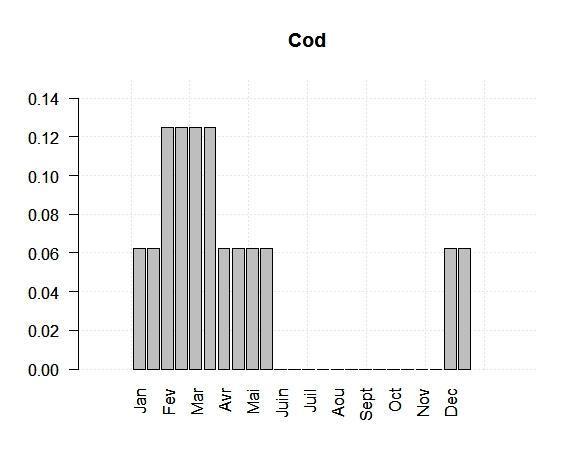

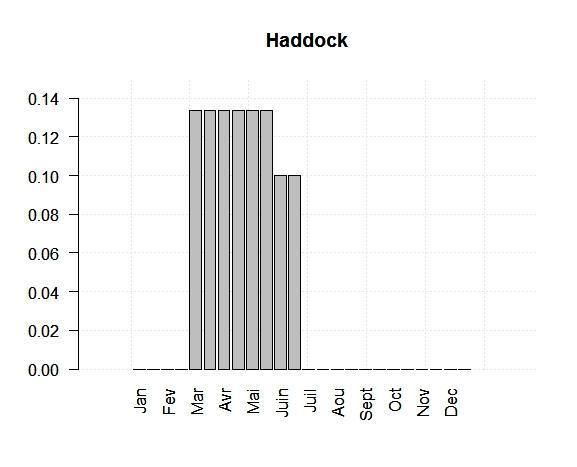

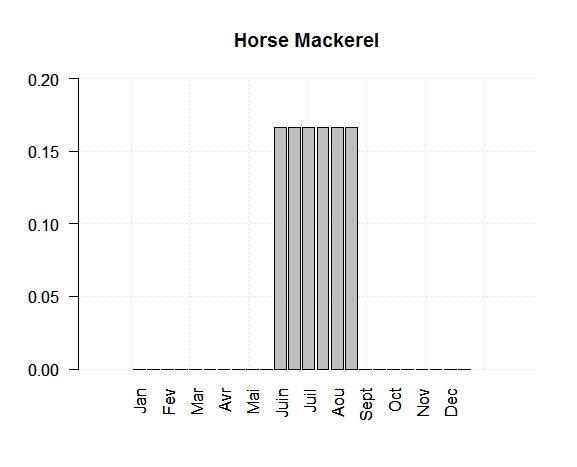

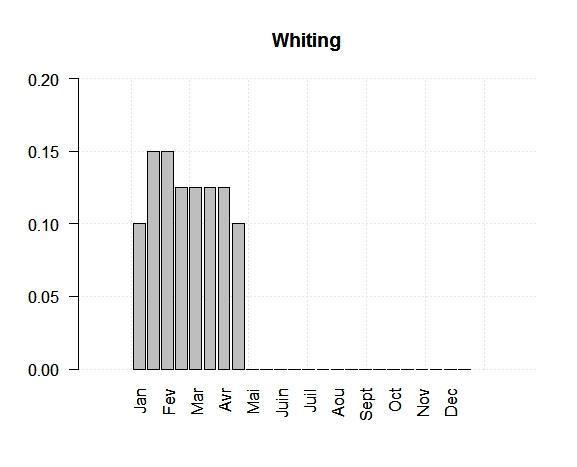

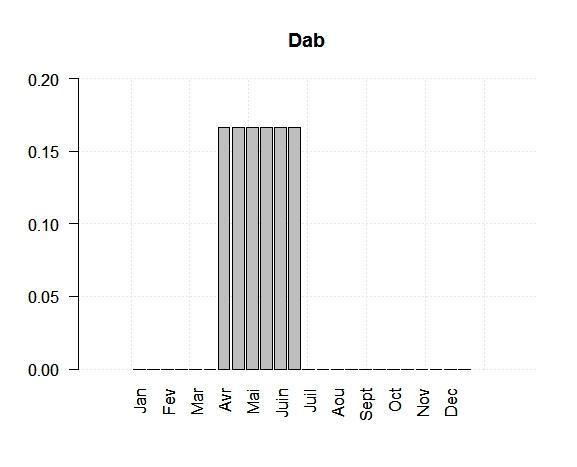

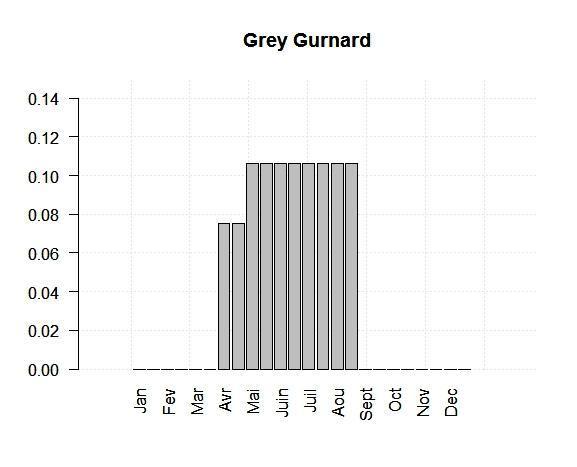

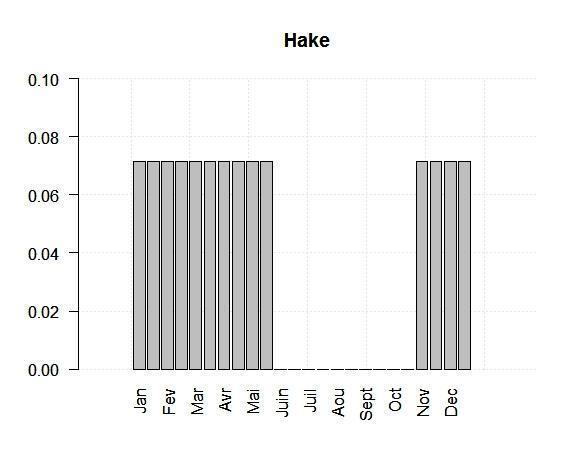

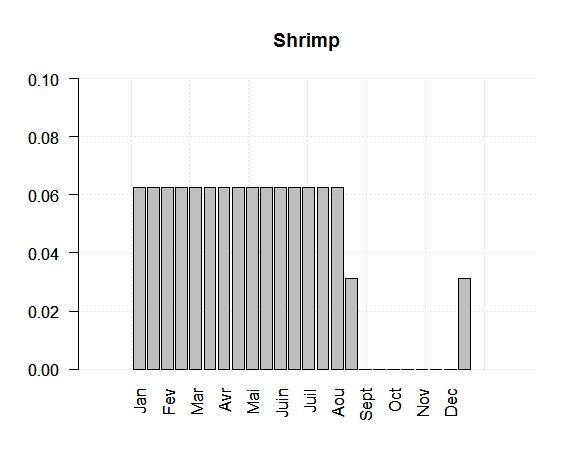

Supporting Information S9: **Monthly maps of LTL and temperature and oxygen variables from POLCOMS-ERSEM averaged on the 2010-2019 period**. The LTL biomasses are vertically integrated and provided as tons per cell. Temperature and oxygen variables are available in the entire water column (vertically integrated) and at the sea bottom. Temperature is in Celsius, Oxygen is in % of saturation.

| LTL groups | Maps |
| --- | --- |
| Micro-phytoplankton | 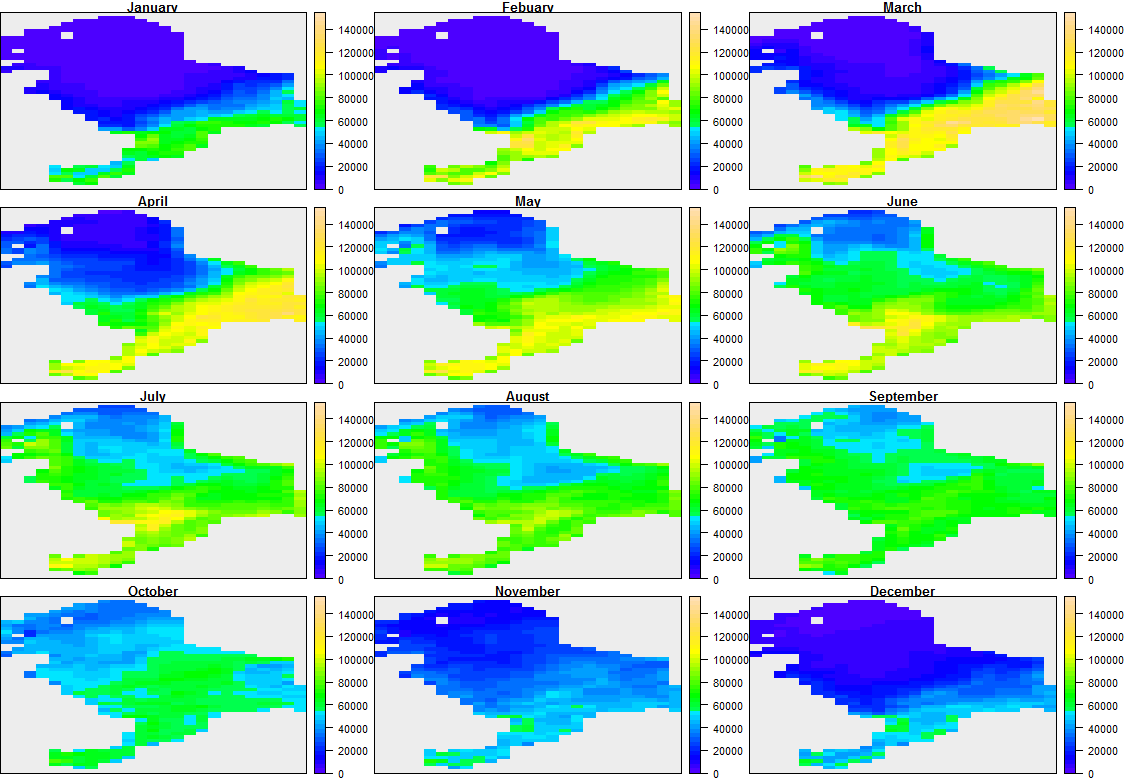 |
| Diatoms | 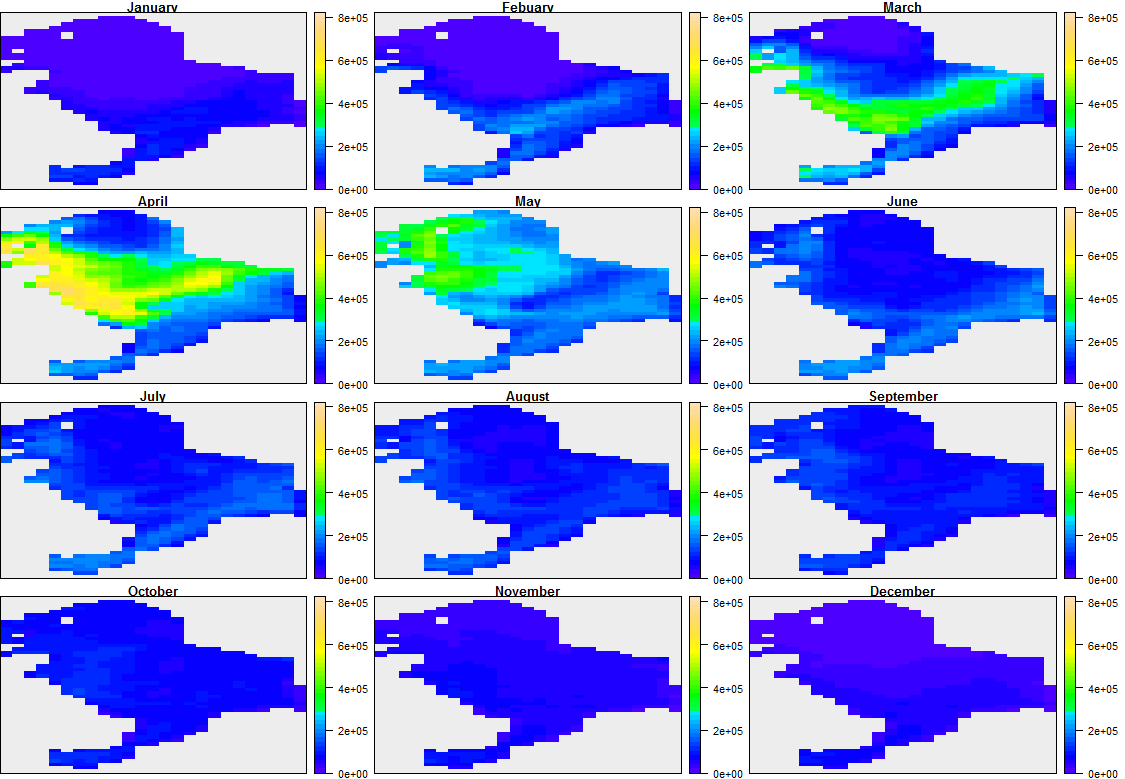 |
| Hetero-trophic flagellates | 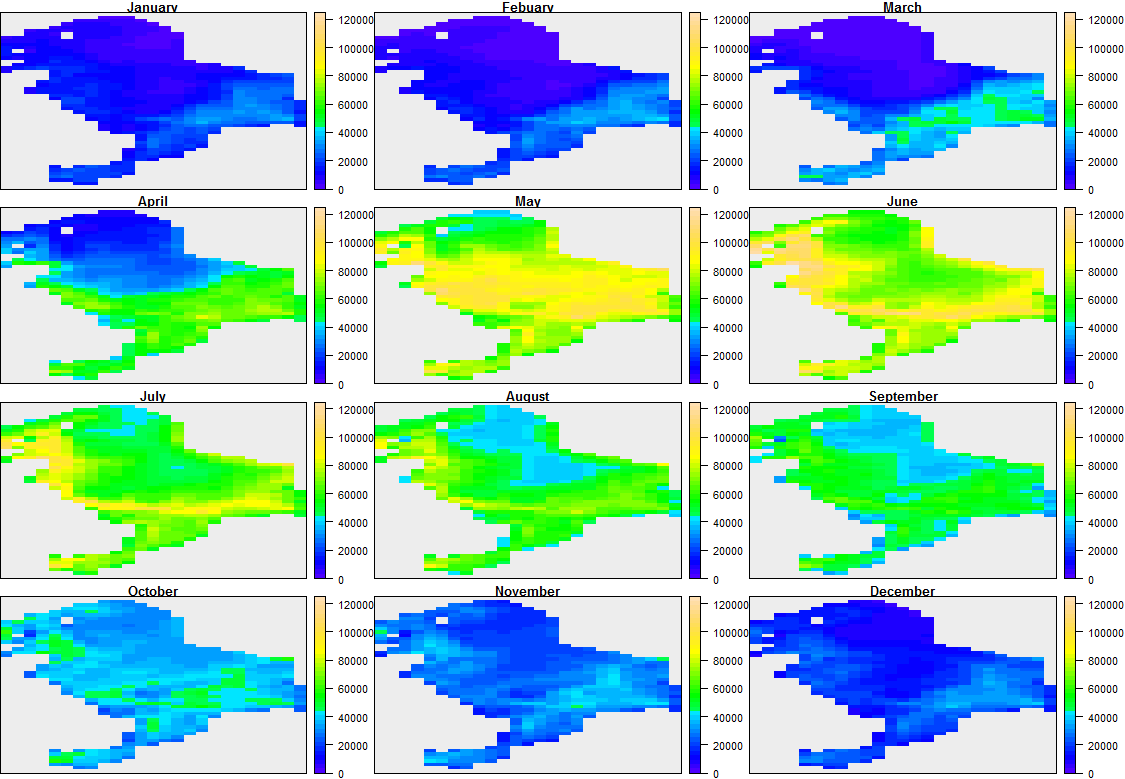 |
| Micro-zooplankton | 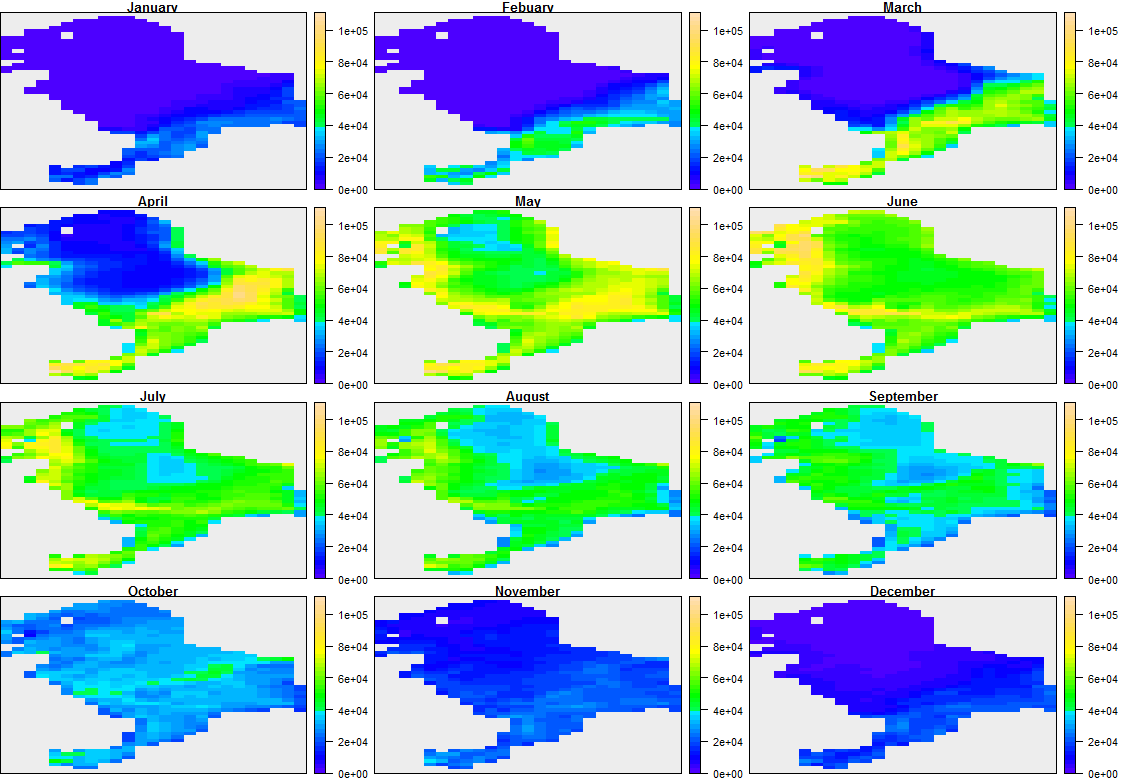 |
| Meso-zooplankton | 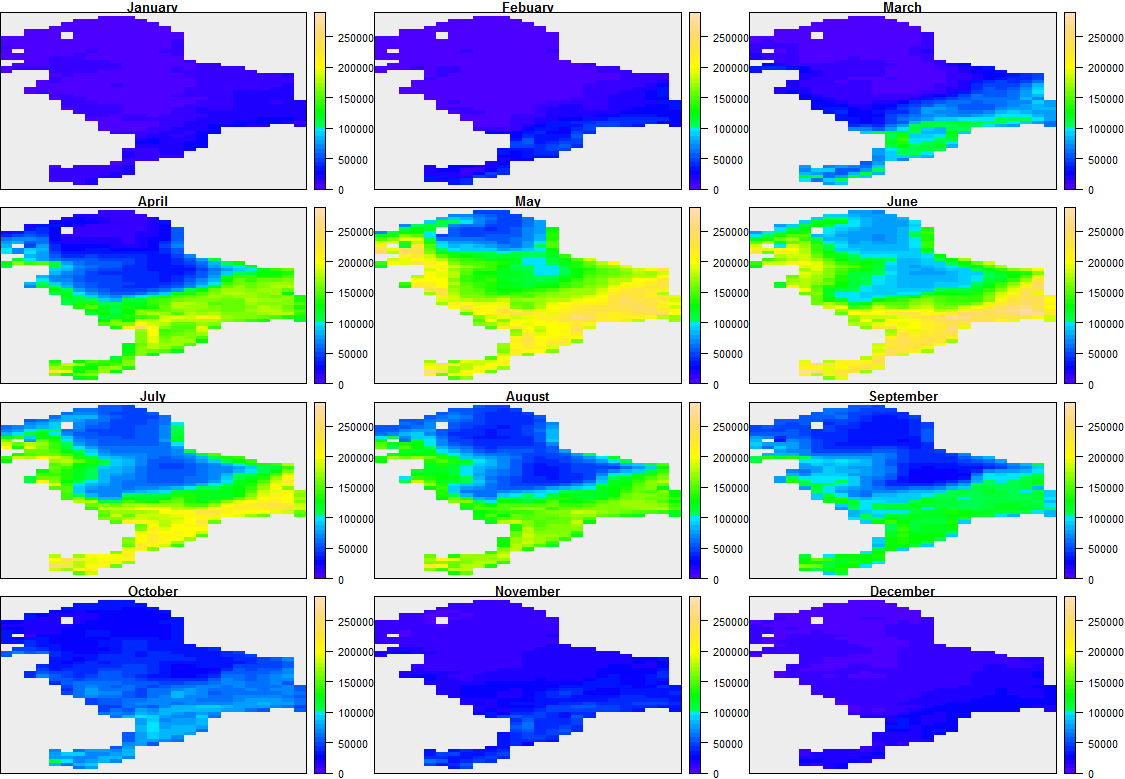 |
| SuspensionFeeders | 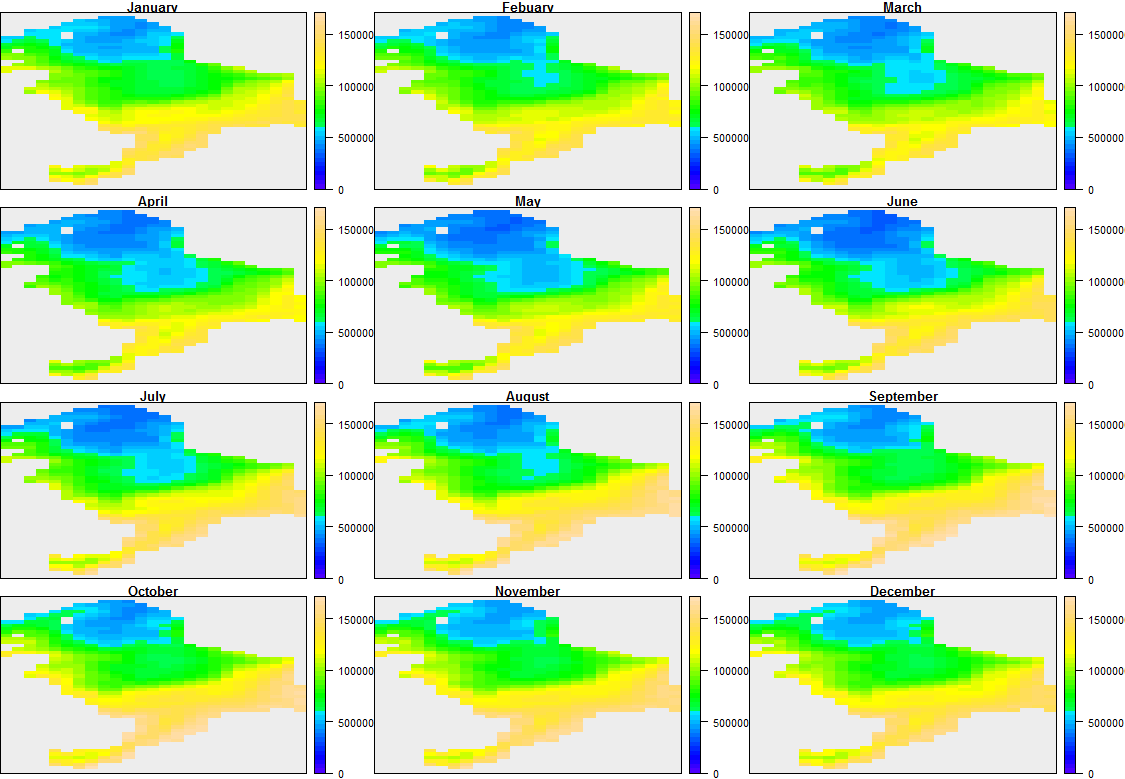 |
| DepositFeeders | 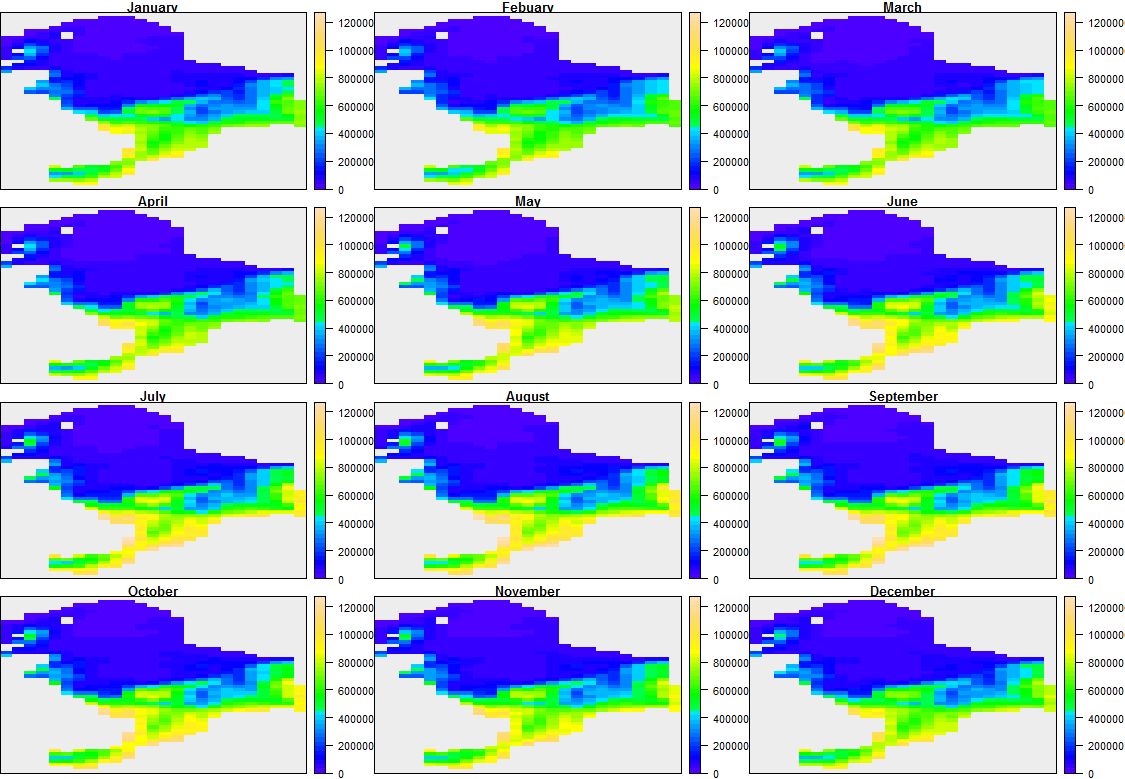 |
| Meiobenthos | 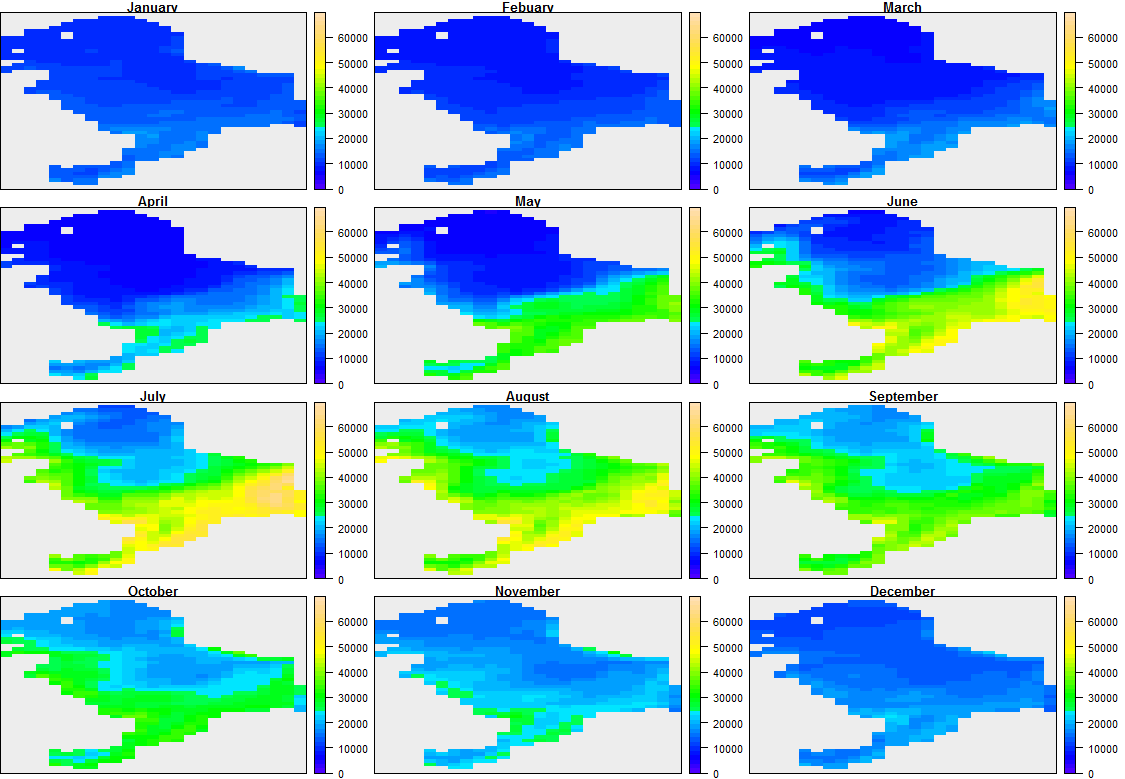 |
| Temperature | Column integration  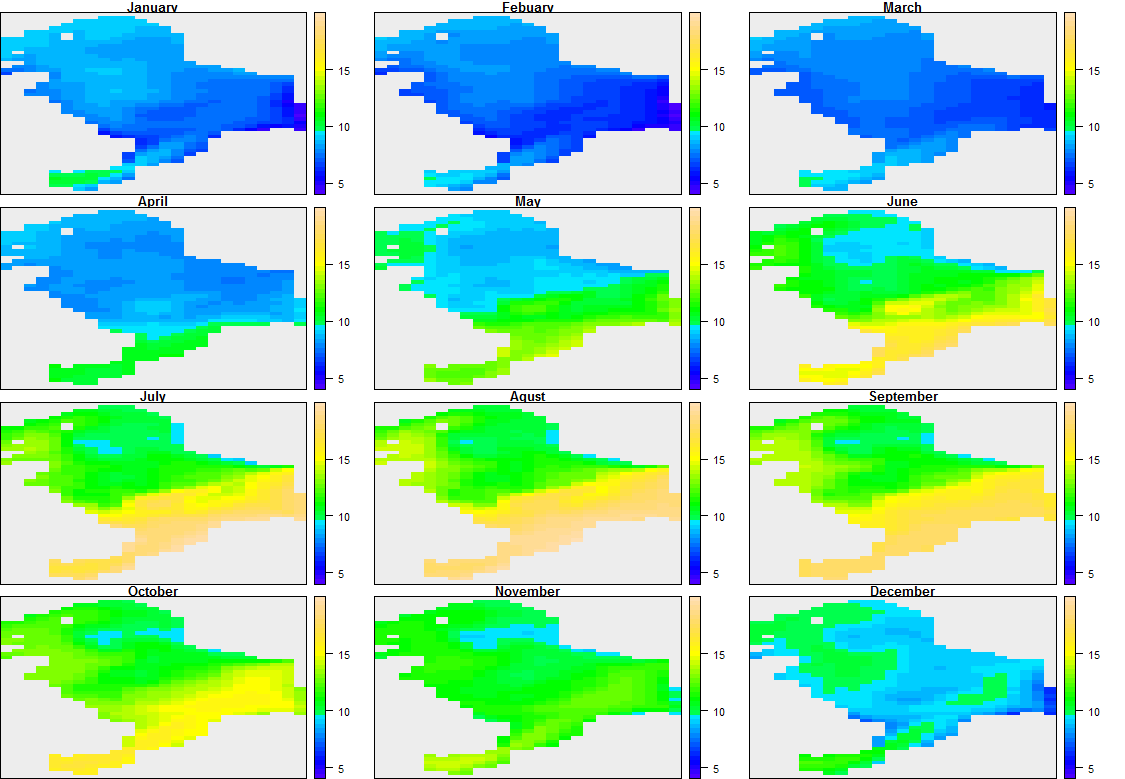  Bottom  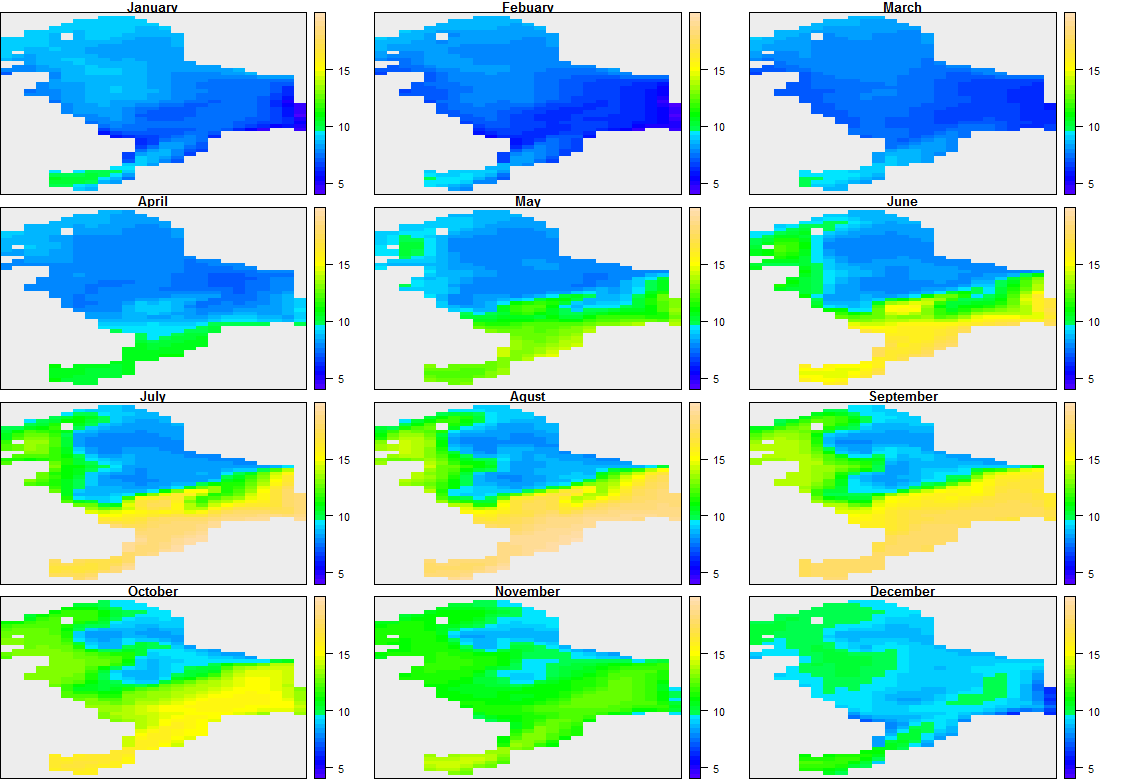 |
| Oxygen | Column integration  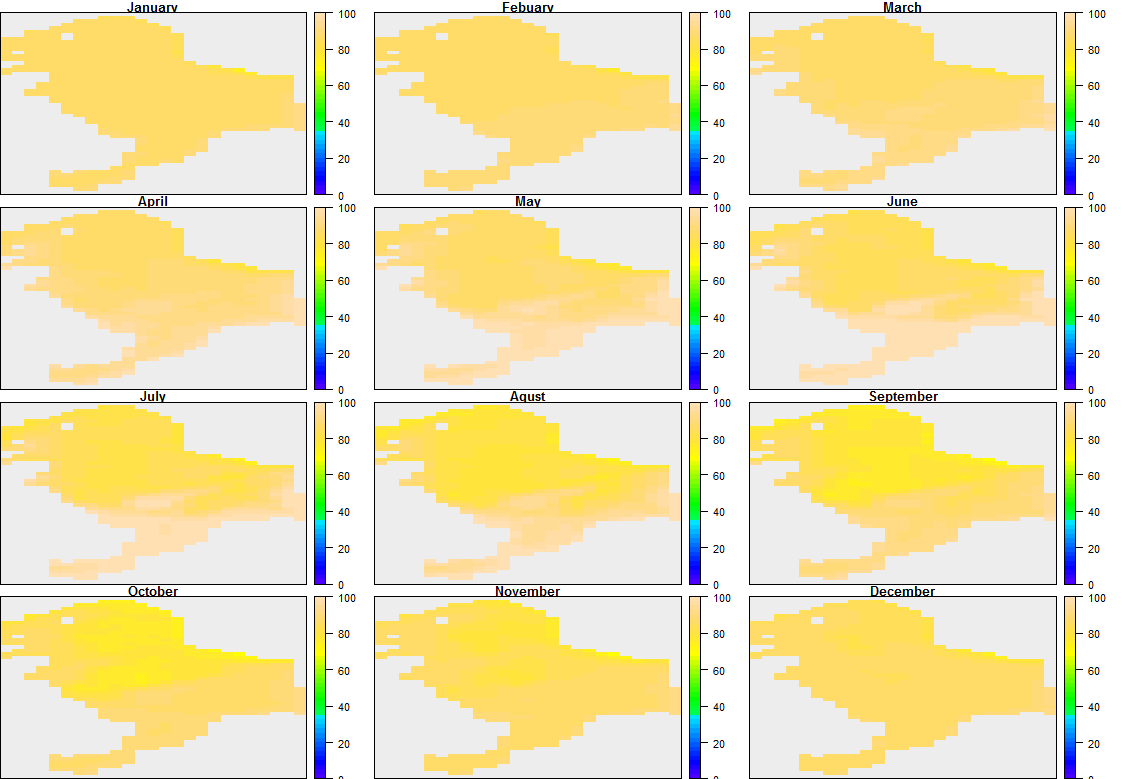  Bottom  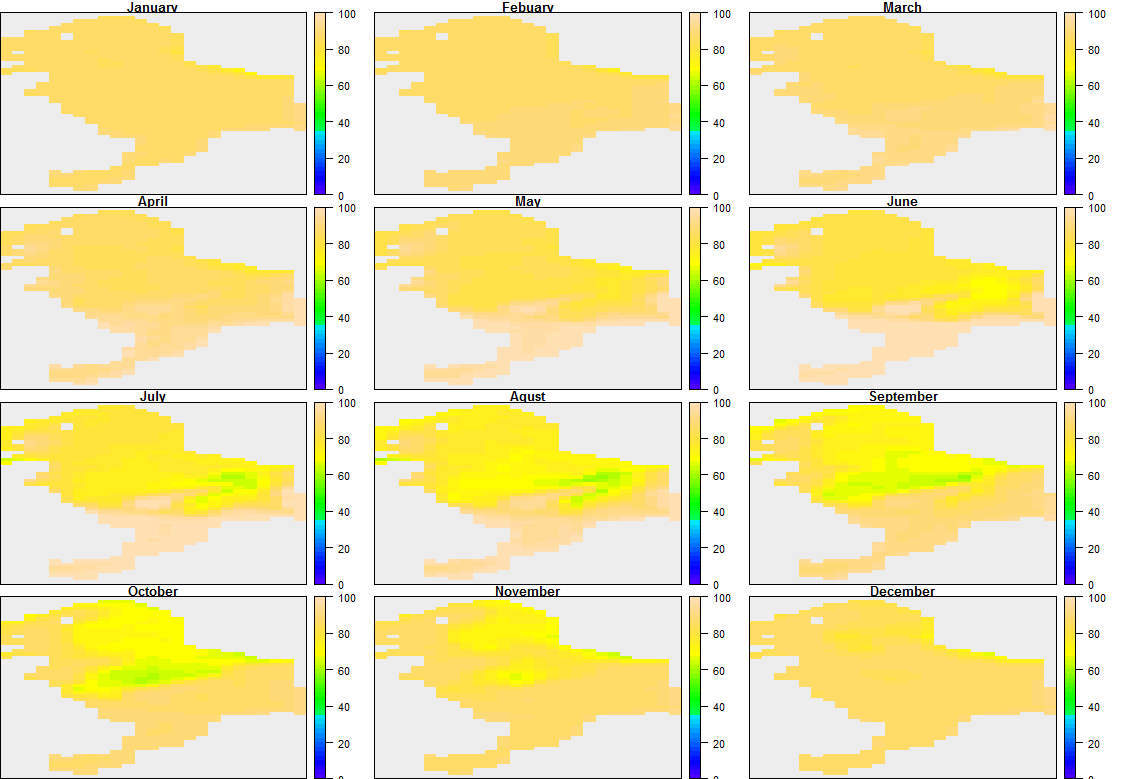 |

Supporting Information S10**:** **Target values for the calibration. The target values were average on the period 2010 to 2019.** Data used to calibrate Bioen-OSMOSE-NS are fisheries landings (ICES, 2019a), size-at-age from scientific surveys (NS-IBTS-Q1, ICES. North Sea International Bottom Trawl Survey (2010-2019). Available online at http://datras.ices.dk) and estimated biomasses for assessed species (ICES, 2016, 2018a, 2018b, 2018c, 2019b) (see section 2.2.3 Calibration for more details).

|  | | Herring | Mackerel | Sandeel | Sprat | N. Pout | Plaice | Sole | Saithe | Cod | Haddock | H. Mackerel | Whiting | Dab | G. Gurnard | Hake | Shrimp |
| --- | --- | --- | --- | --- | --- | --- | --- | --- | --- | --- | --- | --- | --- | --- | --- | --- | --- |
| Biomass (10^3^ tonnes) | | 4 252 | 1 719 | 1 735 | 1 309 | 393 | 969 | 78 | 354 | 206 | 379 |  | 363 | 201 |  | 62 |  |
| Catches (10^3^ tonnes) | | 443 | 285 | 292 | 161 | 57 | 119 | 16 | 76 | 41 | 36 | 21 | 31 | 49 | 7 | 18 | 37 |
| Size-at-age (cm) | 1 | 16.4 | 22.9 |  | 9.3 | 13 | 15.7 | 13 | 22.9 | 23.9 | 22.1 | 11.4 | 18.5 | 11.4 | 13.6 | 33.3 |  |
|  | 2 | 21.9 | 28.1 | 15.9 | 11.4 | 16.4 | 20.2 | 18.2 | 34.4 | 39.1 | 30 | 20 | 24.9 | 14.8 | 18.3 | 44.2 |  |
|  | 3 | 25.2 | 31.4 | 17.5 | 12.8 | 18.4 | 24.4 | 23.2 | 41.7 | 55.8 | 35.3 | 20 | 30.2 | 18.8 | 22.4 | 58.6 |  |
|  | 4 | 27.2 | 33.3 | 17.4 | 13.7 | 19.9 | 27.9 | 26.7 | 48 | 69 | 38.7 | 26 | 34.2 | 22 | 27 | 68.4 |  |
|  | 5 | 28.5 | 34.3 | 17.8 | 14.3 | 20.6 | 30.2 | 27.7 | 54.5 | 77.9 | 40.2 | 34 | 36.6 | 23.5 | 28.7 | 75.8 |  |
|  | 6 | 29.5 | 35 | 18.3 | 14.5 |  | 31.6 | 27.6 | 61.8 | 85.5 | 41.4 | 33 | 37.9 | 25.2 | 27.5 | 77.5 |  |
|  | 7 | 30.1 | 36.1 |  | 14.8 |  | 32.4 | 30.6 | 69.7 | 90.2 | 43.4 | 33.9 | 38.8 | 25.7 | 27.8 | 87.3 |  |
|  | 8 | 30.6 | 36.6 |  | 15 |  | 32.9 | 28.8 | 75.8 | 96.1 | 44.5 | 35.6 | 38.9 | 27.1 | 26.7 | 85.1 |  |
|  | 9 | 30.7 | 37.4 |  | 15 |  | 32.9 | 28.8 | 80.8 | 97.4 | 45.4 | 36.4 | 38.3 | 27.3 | 27.7 | 88.7 |  |
|  | 10 | 31.1 | 38 |  | 15 |  | 33 | 33.2 | 85.2 | 105.1 |  | 36.7 | 38.4 |  | 27.8 |  |  |
|  | 11 | 31.5 | 38.7 |  |  |  | 33.3 | 30.7 | 87.1 | 103 |  | 37 |  |  | 31 |  |  |
|  | 12 | 31.9 | 38.8 |  |  |  | 32.9 | 32.8 | 89.8 |  |  | 37 |  |  | 29 |  |  |
|  | 13 | 32.3 | 38.7 |  |  |  | 34.1 | 31 | 93.3 |  |  |  |  |  |  |  |  |
|  | 14 | 32.5 | 40.1 |  |  |  | 33.4 |  | 97.4 |  |  |  |  |  |  |  |  |
|  | 15 |  |  |  |  |  | 33.6 |  | 94.8 |  |  |  |  |  |  |  |  |
|  | 16 |  |  |  |  |  | 35.7 |  | 98.9 |  |  |  |  |  |  |  |  |

Supporting Information S11: **Fishing selectivity per species.**

Fishing size-selectivity per species. We used fishing mortality rates estimated from stock assessments (ICES, 2016, 2018a, 2018b, 2018c, 2019b). When available, fishing mortality rates per age class were converted to fishing mortality rates per size class using species specific von Bertalanffy growth models. When the only data available is a constant fishing mortality rate for a species and a size at recruitment, a knife-edge selectivity curve is used (sprat, dab, grey gurnard, hake and shrimp).

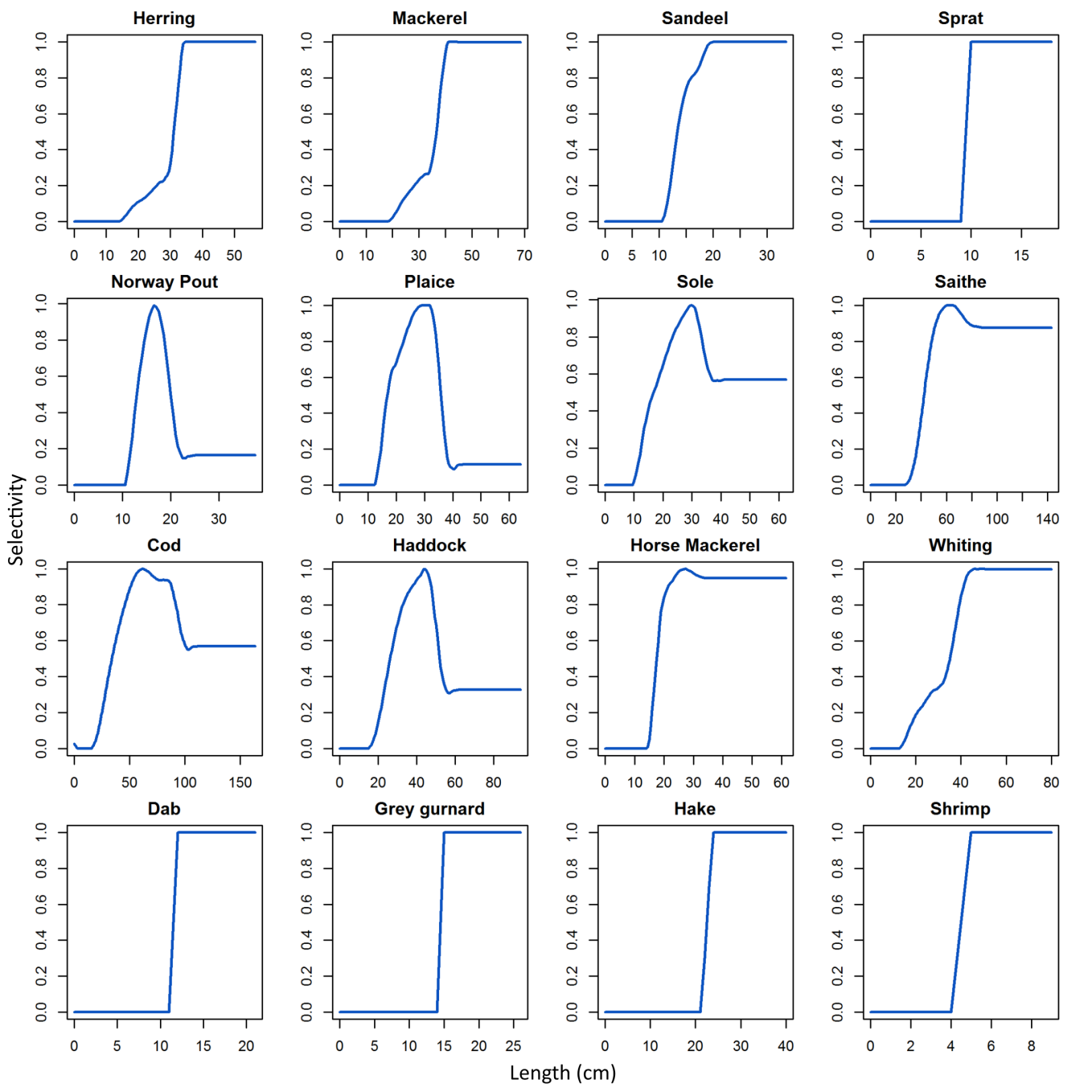

### 1. Supporting Information references

Barot, S., Heino, M., O’Brien, L., and Dieckmann, U. (2004). Estimating reaction norms for age and size at maturation when age at first reproduction is unknown. Evol Ecol Res 6: 659–678

Baxter, I.G. (1958). Fecundities of winter-spring and summer-autumn herring spawners. ICES: C.M. 42:9.

Behrens, J.W., and Steffensen, J.F. (2007). The effect of hypoxia on behavioural and physiological aspects of lesser sandeel, Ammodytes tobianus (Linnaeus, 1785). Mar Biol 150, 1365–1377. https://doi.org/10.1007/s00227-006-0456-4.

Bergstad, O.A., Høines, A.S., and Krüger-Johnsen, E.M. (2001). Spawning time, age and size at maturity, and fecundity of sandeel, Ammodytes marinus, in the northeastern North Sea and in unfished coastal waters off Norway. Aquat. Living Resour. 14:293-301.

Bilgin, S., and Samsun, O. (2006). Fecundity and Egg Size of Three Shrimp Species, Crangon crangon, Palaemon adspersus, and Palaemon elegans (Crustacea: Decapoda: Caridea), off Sinop Peninsula (Turkey) in the Black Sea.

Bohl, H. (1956). On the biology of the dab in the North Sea. ICES: C.M. 5pp.

Boukal, D.S., Dieckmann, U., Enberg, K., Heino, M., and Jørgensen, C. (2014). Life-history implications of the allometric scaling of growth. Journal of Theoretical Biology 359, 199–207. https://doi.org/10.1016/j.jtbi.2014.05.022.

Christiansen, J.S., Fevolden, S.E., Karamushlo, O.V., and Karamushko, L.I. (1997). Reproductive traits of marine fish in relation to their mode of oviposition and zoogeographic distribution. ICES CM 1997/CC. 14 p.

Claireaux, G., Webber, D.M., Lagardère, J.-P., and Kerr, S.R. (2000). Influence of water temperature and oxygenation on the aerobic metabolic scope of Atlantic cod (Gadus morhua). Journal of Sea Research 44, 257–265. https://doi.org/10.1016/S1385-1101(00)00053-8.

Clarke, A., and Johnston, N.M. (1999). Scaling of metabolic rate with body mass and temperature in teleost fish. Journal of Animal Ecology 13.

Cohen, D.M., Inada, T., Iwamoto, T., and Scialabba, N. (1990). Gadiform fishes of the world (Order Gadiformes). An annotated and illustrated catalogue of cods, hakes, grenadiers and other gadiform fishes known to date. FAO Fish. Synop. 125(10). Rome: FAO. Vol. 10., 442 p.

Coull, K.A., Johnston, R., and Rogers, S.I. (1998). Fisheries Sensitivity Maps in British Waters. 63.

Dahlke, F., Wohlrab, S., Butzin, M., and Pörtner, H.-O. (2020). Experimental data compilation, thermal tolerance and thermal responsiveness of fish species and life stages (PANGAEA).

van Damme, C.J.G., Bolle, L.J., Fox, C.J., Fossum, P., Kraus, G., Munk, P., Rohlf, N., Witthames, P.R., and Dickey-Collas, M. (2009). A reanalysis of North Sea plaice spawning-stock biomass using the annual egg production method. ICES J. Mar. Sci. 66:1999-2011.

De Silva, S.S. (1973). Food and feeding habits of the herring Clupea harengus and the sprat C. sprattus in inshore waters of the west coast of Scotland. Marine Biology 20, 282–290. https://doi.org/10.1007/BF00354272.

Dickson, K.A. (2002). Temperature effects on sustained swimming in mackerel.

Dupont-Prinet, A., Pillet, M., Chabot, D., Hansen, T., Tremblay, R., and Audet, C. (2013). Northern shrimp (Pandalus borealis) oxygen consumption and metabolic enzyme activities are severely constrained by hypoxia in the Estuary and Gulf of St. Lawrence. Journal of Experimental Marine Biology and Ecology 448, 298–307. https://doi.org/10.1016/j.jembe.2013.07.019.

Eltink, A., and Vingerhoed, B. (1989). The total fecundity of Western horse mackerel (Trachurus trachurus L.). ICES:C.M. H(44):11p.

Fox, C.J., Planque, B., and Darby, C.D. (2000). Synchrony in the recruitment time-series of plaice (Pleuronectes platessa L) around the United Kingdom and the influence of sea temperature. J. Sea Res. 44:159-168.

Geist, S.J., Ekau, W., and Kunzmann, A. (2013). Energy demand of larval and juvenile Cape horse mackerels, Trachurus capensis, and indications of hypoxia tolerance as benefit in a changing environment. Mar Biol 160, 3221–3232. https://doi.org/10.1007/s00227-013-2309-2.

Hattab, T., Albouy, C., Lasram, F.B.R., Somot, S., Le Loc’h, F., and Leprieur, F. (2014). Towards a better understanding of potential impacts of climate change on marine species distribution: a multiscale modelling approach. Global Ecology and Biogeography 23, 1417–1429.

Hehir, I. (2003). Age, growth and reproductive biology og whiting Merlangius merlangus in the Celtic Sea. Galway-Mayo Institute of Technology. Master thesis. 210p.

Hislop, J.R.G. (1966). A note on the fecundity of whiting in the North Sea. ICES C.M. 19:9.

ICES (2012). Report of the Working Group on the Assessment of Demersal Stocks in the North Sea and Skagerrak (WGNSSK), 27 April - 03 May 2012, ICES Headquarters, Copenhagen. ICES CM 2012/ACON:13. 1346 p.

ICES (2016). Report of the Benchmark Workshop on Sandeel (WKSand 2016). (31 October - 4 November. No. ICES CM 2016/ACOM:33).

ICES (2018a). Report of the Herring Assessment Working Group for the Area South of 62°N (HAWG) (29-31 January 2018 and 12-20 March 2018).

ICES (2018b). Report of the Working Group for the Bay of Biscay and the Iberian Waters Ecoregion (WGBIE) (3-10 May 2018. No. ICES CM 2018/ACOM:12).

ICES (2018c). Report of the Working Group on the Assessment of Demersal Stocks in the North Sea and Skagerrak (WGNSSK) (24 April - 3 May 2018).

ICES (2019a). Catches in FAO area 27 by country, species, area and year as provided by the national authorities. Source: Eurostat/ICES data compilation of catch statistics - ICES 2019, Copenhagen. Version: 16-09-2019.

ICES (2019b). Working group on widely distributed stocks (WGWIDE). ICES Scientific Reports. 1:36. 948 pp. http://doi.org/10.17895/ices.pub.5574.

Koli, L. (1990). Suomen kalat. [Fishes of Finland]. Werner Söderström Osakeyhtiö. Helsinki. 357 p. (in Finnish).

Kopp, D., Lefebvre, S., Cachera, M., Villanueva, M.C., and Ernande, B. (2015). Reorganization of a marine trophic network along an inshore–offshore gradient due to stronger pelagic–benthic coupling in coastal areas. Progress in Oceanography 130, 157–171. https://doi.org/10.1016/j.pocean.2014.11.001.

Lefrançois, C., and Claireaux, G. (2003). Influence of ambient oxygenation and temperature on metabolic scope and scope for heart rate in the common sole Solea solea. Marine Ecology Progress Series 259, 273–284. https://doi.org/10.3354/meps259273.

Lester, N.P., Shuter, B.J., and Abrams, P.A. (2004). Interpreting the von Bertalanffy model of somatic growth in fishes: the cost of reproduction. Proceedings of the Royal Society of London B: Biological Sciences 271, 1625–1631. https://doi.org/10.1098/rspb.2004.2778.

Macer, C.T. (1976). Observations on the maturity and fecundity of mackerel (Scomber scombrus L.). ICES: C.M. H:6(mineo).

MacKenzie, B.R., and Köster, F.W. (2004). Fish Production and Climate: Sprat in the Baltic Sea. Ecology 85, 784–794.

Mackinson, S., and Daskalov, G. (2007). An ecosystem model of the North Sea to support an ecosystem approach to fisheries management: description and parameterisation. 200.

Mehault, S., Dominguez-Petit, R., Cervino, S., and Saborido-Rey, F. (2010). Variability in total egg production and implications for management of the southern stock of European hake. Fish. Res. 104(1-3):111-122.

Meskendahl, L. (2013). Metabolic rates and feeding behaviour of sprat, Sprattus sprattus L. PhD. University of Hamburg.

Muus, B.J., and Nielsen, J.G. (1999). Sea fish. Scandinavian Fishing Year Book, Hedehusene, Denmark. 340 p.

Narberhaus, I., Krause, J., and Bernitt, U. (2012). Threatened biodiversity in the German North and Baltic seas. Naturschutz und Biologische Vielfalt, Heft 117. Federal Agency for Nature Conservation, Bonn, Germany.

Oh, C.-W., Hartnoll, R.G., and Nash, R.D.M. (2001). Feeding ecology of the common shrimp Crangon crangon in Port Erin Bay, Isle of Man, Irish Sea. Marine Ecology Progress Series 214, 211–223. https://doi.org/10.3354/meps214211.

Pinnegar, J.K. (2014). DAPSTOM - An Integrated Database & Portal for Fish Stomach Records. Version 4.7. Centre for Environment, Fisheries & Aquaculture Science, Lowestoft, UK. February 2014, 39pp.

Quéro, J.C., Desoutter, M., and Lagardère, F. (1986). Soleidae. p. 1308-1324. In P.J.P. Whitehead, M.-L. Bauchot, J.-C. Hureau, J. Nielsen and E. Tortonese (eds.) Fishes of the North-eastern Atlantic and the Mediterranean. UNESCO, Paris. Vol. 3.

Quince, C., Abrams, P.A., Shuter, B.J., and Lester, N.P. (2008). Biphasic growth in fish I: Theoretical foundations. Journal of Theoretical Biology 254, 197–206. https://doi.org/10.1016/j.jtbi.2008.05.029.

Sonina, M.A. (1971). La fécondité de l´églefin arcto-norvegien (Melanogrammus aeglefinus L.). ICES: C.M. F(16):11.

Steinhausen, M.F., Steffensen, J.F., and Andersen, N.G. (2005). Tail beat frequency as a predictor of swimming speed and oxygen consumption of saithe (Pollachius virens) and whiting (Merlangius merlangus) during forced swimming. Marine Biology 148, 197–204. https://doi.org/10.1007/s00227-005-0055-9.

Sundbly, S., Kristiansen, T., Nash, R., and Johannessen, T. (2017). Dynamic Mapping of North Sea Spawning – Report of the KINO Project - Semantic Scholar.

Van der Land, M.A. (1991). Distribution of flatfish eggs in the 1989 egg surveys in the southeastern North Sea, and mortality of plaice and sole eggs. Netherlands Journal of Sea Research 27, 277–286. https://doi.org/10.1016/0077-7579(91)90030-5.

West, G.B., Brown, J.H., and Enquist, B.J. (1997). A General Model for the Origin of Allometric Scaling Laws in Biology. Science 276, 122–126. <https://doi.org/10.1126/science.276.5309.122>.

WGCRAN (2015). Report of the Working Group on Crangon Fisheries and Life History.

Witthames, P.R., Greer Walker, M., Dinis, M.T., and Whiting, C.L. (1995). The geographical variation in the potential annual fecundity of dover sole Solea solea (L.) from European shelf waters during 1991. Netherlands Journal of Sea Research 34, 45–58. https://doi.org/10.1016/0077-7579(95)90013-6.

Yoneda, M., and Wright, P.J. (2004). Temporal and spatial variation in reproductive investment of Atlantic cod Gadus morhua in the northern North Sea and Scottish west coast. Mar. Ecol. Prog. Ser. 276:237-248.
